# Selective inhibitors of JAK1 targeting a subtype-restricted allosteric cysteine

**DOI:** 10.1101/2022.01.31.478302

**Authors:** Madeline E. Kavanagh, Benjamin D. Horning, Roli Khattri, Nilotpal Roy, Justine P. Lu, Landon R. Whitby, Jaclyn C. Brannon, Albert Parker, Joel M. Chick, Christie L. Eissler, Ashley Wong, Joe L. Rodriguez, Socorro Rodiles, Kim Masuda, John R. Teijaro, Gabriel M. Simon, Matthew P. Patricelli, Benjamin F. Cravatt

## Abstract

The JAK family of non-receptor tyrosine kinases includes four subtypes (JAK1, JAK2, JAK3, and TYK2) and is responsible for signal transduction downstream of diverse cytokine receptors. JAK inhibitors have emerged as important therapies for immuno(onc)ological disorders, but their use is limited by undesirable side effects presumed to arise from poor subtype selectivity, a common challenge for inhibitors targeting the ATP-binding pocket of kinases. Here, we describe the chemical proteomic discovery of a druggable allosteric cysteine present in the non-catalytic pseudokinase domain of JAK1 (C817) and TYK2 (C838), but absent from JAK2 or JAK3. Electrophilic compounds selectively engaging this site block JAK1-dependent transphosphorylation and cytokine signaling, while appearing to act largely as “silent” ligands for TYK2. Importantly, the allosteric JAK1 inhibitors do not impair JAK2-dependent cytokine signaling and are inactive in cells expressing a C817A JAK1 mutant. Our findings thus reveal an allosteric approach for inhibiting JAK1 with unprecedented subtype selectivity.

---

Dysregulated cytokine signaling is central to the pathophysiology of a wide range of diseases, including autoimmune and inflammatory conditions, cardiovascular, gastrointestinal and neurodegenerative diseases, and cancer^1,2^. More than 50 different cytokines signal through a family of non-receptor Janus tyrosine kinases (JAKs), which, in humans, consists of JAK1, JAK2, JAK3, and TYK2^1,3^. JAKs associate with the intracellular tail of specific cytokine receptors and are activated by receptor-induced dimerization to phosphorylate themselves in *trans*, the receptor, and downstream signaling proteins, including the STAT family of transcription factors. The specific combination of JAK enzymes and STAT transcription factors that are activated by a given cytokine is cell-type and context-dependent, allowing the JAK-STAT system to regulate diverse biological and disease processes^3^.

The key role of JAK-STAT pathways in immunology and cancer has motivated the pursuit of JAK inhibitors, and many pan-JAK inhibitors have been described^1,4^. These compounds have provided preclinical and clinical evidence that inhibiting JAK-STAT signaling can alleviate aberrant cytokine responses and have established JAKs as important therapeutic targets^1,4^. There are currently seven FDA-approved JAK inhibitors for the treatment of rheumatoid arthritis, psoriasis, ulcerative colitis, atopic dermatitis, and/or myeloproliferative diseases (e.g. polycythemia, leukemia and GVHD), and one compound (baracitinib) that has emergency use authorization (EUA) for COVID-19^4^. All FDA-approved JAK inhibitors act by an orthosteric mechanism, meaning that they bind to the conserved ATP pocket of the kinase domain, and, even though individual compounds have differing relative selectivity profiles across the JAK family, they all inhibit more than one JAK isoform with moderate-to-high potency (IC_50_ < 1 μM)^5–12^. This lack of selectivity has important translational implications, as there is growing concern over an array of adverse side effects caused by JAK inhibitors^1,3,4,13^, including dose-limiting cytopenias thought to be due to inhibition of JAK2-mediated growth factor receptor signaling^14^, an increased risk of cardiovascular events and infections, dyslipidemia, and elevated liver enzymes^3^. In 2021, these concerns prompted the FDA to place a “black box” warning on JAK inhibitors indicated for chronic conditions such as rheumatoid arthritis^15^.

The lack of subtype selectivity of FDA-approved JAK inhibitors may contribute to their adverse clinical effects^16,17^, and there is accordingly considerable interest in the discovery of JAK inhibitors with improved specificity. Such compounds might not only constitute next-generation therapeutics, but would also serve as valuable research tools to better understand the unique contributions made by each JAK isoform to physiology and disease. Subtype-selective JAK inhibitors have been pursued by multiple strategies. For example, covalent inhibitors of JAK3 have been developed, such as ritlecitinib, that target a cysteine (C909) uniquely found in the activation loop of this kinase compared to other JAKs^18,19^. While this approach achieves specificity for JAK3 over other JAKs, ritlecitinib cross-reacts with TEC family kinases, which also share a cysteine at an equivalent position. JAKs are distinguished from many other kinases by having an additional non-catalytic pseudokinase (JH2) domain that regulates kinase activity and is a hotspot for gain- or loss-of-function mutations^2,20^. Notably, compounds binding to the ATP pocket of the JH2 domain of TYK2 have been found to inhibit this kinase with remarkable functional selectivity over JAK1-JAK3^21,22^, and one of these agents, – BMS-986165 (deucravacitinib) – is in late-stage clinical development for autoimmune disorders^21,23,24^.

In contrast to the progress made on subtype-restricted JAK3 and TYK2 inhibitors, selective JAK1 inhibitors are still lacking. Although some orthosteric JAK1 inhibitors have been reported that display improved subtype selectivity, these compounds (e.g., abrocitinib, filgotinib) still generally show substantial cross-reactivity with JAK2 (e.g., < 1 μM IC_50_ values), depending on the biochemical or cellular assay employed^12,25,26^. The generation of highly selective inhibitors of JAK1 is an important objective, as several lines of evidence indicate that JAK1 blockade contributes to the efficacy of pan-JAK inhibitors in chronic autoimmune disorders. For instance, gain-of-function *JAK1* mutations promote multi-organ immune dysregulation^27^ and are associated with specific types of cancer (e.g., leukemia^28,29^ and gynecological tumors^30^), while deleterious mutations cause severe immunosuppression in humans^31^. Additionally, JAK1 is broadly expressed and plays essential and non-redundant roles downstream of class II, γc, and gp130 cytokines^32^, many of which are dysregulated in inflammatory diseases^2^. Nonetheless, the precise contribution of JAK1 to homeostatic immune function and disease remains only partly understood due to a lack of genetic models and selective chemical tools. *JAK1* deletion is perinatal lethal to mice^32^, and consequently, much of our understanding of JAK1 biology has relied on studies with conditional knockout mice lacking JAK1 in specific cell types^33–35^, JAK1-deficient human cell lines^20,36–39^ and/or non-selective orthosteric inhibitors^16,36,40^.

Here, we describe the chemical proteomic discovery of a ligandable allosteric cysteine in the pseudokinase domain of JAK1 (C817) and TYK2 (C838) but absent from JAK2 and JAK3. We optimize an electrophilic compound VVD-118313 (**5a**) that engages JAK1_C817 and TYK2_838 with high potency and proteome-wide selectivity and show that this agent blocks JAK1 signaling in human cancer cell lines and primary immune cells, while sparing JAK2-dependent pathways. VVD-118313 does not inhibit signaling of a C817A JAK1 mutant and appears to act as a silent ligand for TYK2 in the context of primary immune cells. Mechanistic studies indicate that VVD-118313 does not inhibit the catalytic activity of purified JAK1, but potently blocks JAK1 trans-phosphorylation in cells. Integrating our findings with previous work on allosteric TYK2 inhibitors, such as BMS-986165, points to the potential for leveraging multiple druggable pockets in the pseudokinase domain of JAKs to develop inhibitors with unprecedented subtype selectivity.

## Results

### Discovery of a ligandable cysteine in the JAK1/TYK2 pseudokinase domain

Previous activity-based protein profiling (ABPP) studies to assess the interactions of electrophilic small-molecule fragments with cysteines in primary human T cells uncovered a ligandable cysteine shared by JAK1 (C817) and TYK2 (C838)^41^. Both cysteines were substantially engaged by chloroacetamide (KB02) and acrylamide (KB05) fragments (**Extended Data Fig. 1a**)^41^, as determined by mass spectrometry (MS)-ABPP experiments that monitored electrophile-dependent changes in iodoacetamide-desthiobiotin (IA-DTB) reactivity of > 10,000 cysteines in the human T-cell proteome (**Fig. 1a**). Other quantified JAK1 and TYK2 cysteines were unaffected in their IA-DTB reactivity by KB02 or KB05 treatment (**Fig. 1b** and **Extended Data Fig. 1b**).

**Figure 1.**
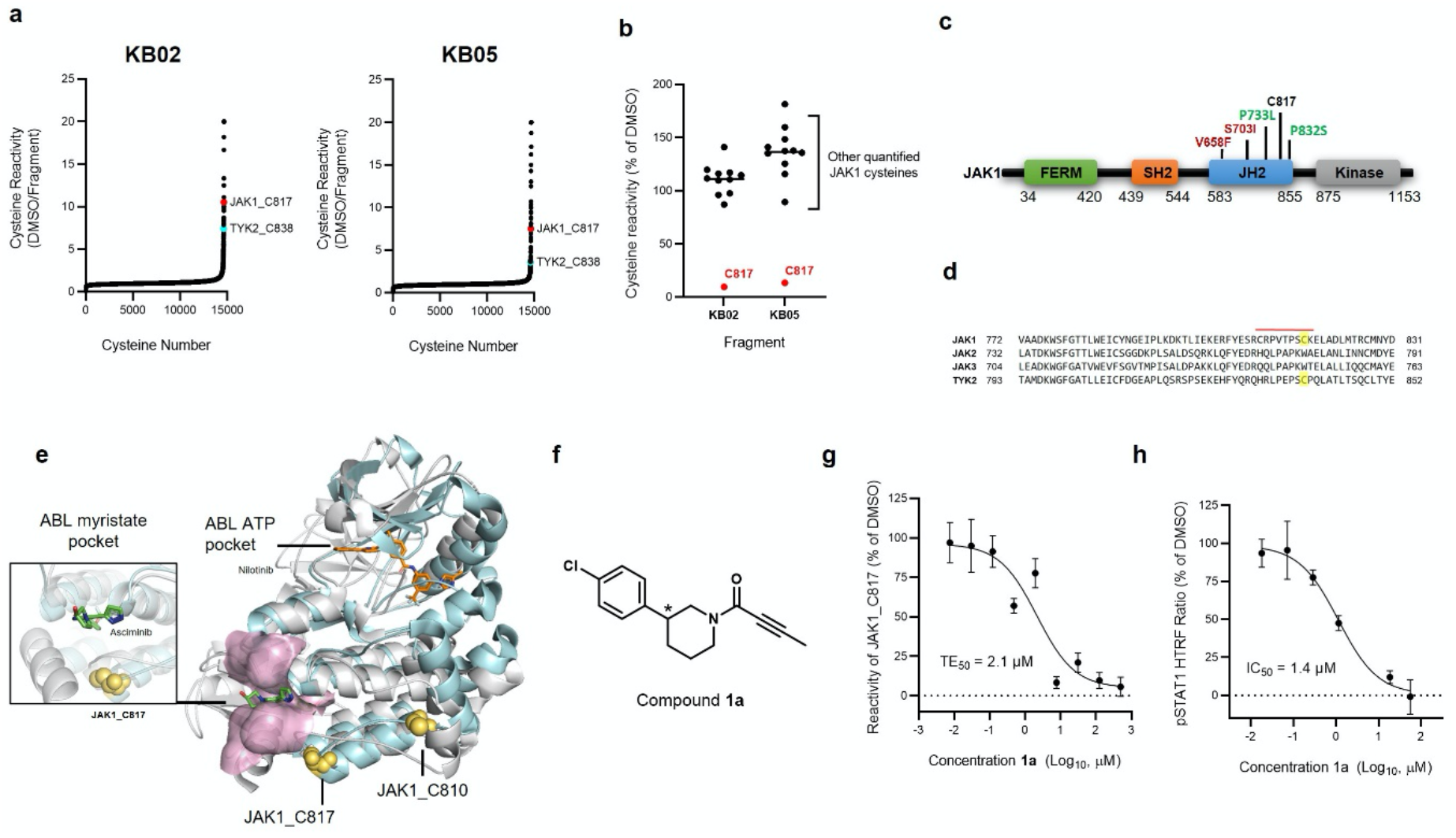
Discovery of a ligandable cysteine in the JAK1/TYK2 pseudokinase domain. **a**, Competition ratios of IA-DTB-labeled and enriched cysteine-containing peptides quantified in MS-ABPP experiments performed using proteomes from human T cells treated *in situ* with the cysteine-reactive small-molecule fragments KB02 and KB05 (50 μM, 1 h) or DMSO control. **b**, Relative MS3 signal intensity values for all quantified IA-DTB-labeled, cysteine-containing peptides in JAK1 in KB02- or KB05-treated T cells compared to DMSO-treated T cells. The KB02- and KB05-liganded cysteine C817 is highlighted in red. Horizontal black bars indicate the median signal intensity for all other quantified JAK1 cysteines. For **a** and **b**, data are mean values combined from soluble and particulate proteomic analyses from two independent experiments performed previously^41^. **c**, Domain structure of JAK1. C817 and select gain-(red) or loss- (green) of-function mutations in the pseudokinase (JH2) domain are highlighted. **d**, Partial amino acid sequence alignment of human JAK family proteins. Electrophilic fragment-liganded cysteines in JAK1 (C817) and TYK2 (C838) are highlighted in yellow. Red bar indicates the tryptic peptide containing JAK1_C817. **e**, Overlay of the x-ray crystal structures of the JAK1 JH2 domain (PDB 4L00) and ABL kinase domain (PDB 5MO4), highlighting the proximity of JAK1 C817 (yellow spheres) to the ABL myristate-binding pocket (pink). The allosteric ABL inhibitor asciminib (green) and orthosteric inhibitor nilotinib (orange) are show in stick representations. **f**, Structure of compound **1a** (*single stereoisomer, absolute configuration not assigned). **g**, Concentration-dependent engagement of JAK1_C817 by **1a** determined by targeted MS-ABPP experiments (1 h compound treatment of human PBMC or Jurkat cell proteome). TE, target engagement. **h**, Concentration-dependent inhibition of IFNα-stimulated STAT1 phosphorylation (pSTAT1) by **1a** in human PBMCs. Cells were treated with **1a** for 2 h followed by 100 ng/mL of IFNα for 30 min, lysed, and pSTAT1 signals measured by HTRF. Data for **g**, **h** are mean values ± S.D. from n = 3 (**g**) or n = 2 (**h**) independent experiments.

JAK1_C817 and TYK2_C838 are located in the catalytically inactive pseudokinase (JH2) domain shared across the JAK family (**Fig. 1c**). The JH2 domain has been found to regulate the kinase activity of the JH1 domain through allosteric mechanisms and is a hotspot for gain- or loss-of-function mutations (**Fig. 1c**)^2,20,42–44^. We noted that other JAK family members – JAK2 and JAK3 – did not share the ligandable cysteine (**Fig. 1d**). A closer examination of the JAK1 JH2 crystal structure in comparison to other kinase structures revealed that C817 is in the C-lobe proximal to a pocket formed by helices αE-F and αH-I, which, in the structurally related kinase ABL, binds an auto-inhibitory *N*-terminal lipid (myristoylation) modification^45–47^ (**Fig. 1e**). This pocket in ABL is targeted by the allosteric inhibitor asciminib, which stabilizes the inactive conformation of the kinase^48^ and has recently been approved for the treatment of chronic myeloid leukemia^49^. Even though asciminib is a reversible inhibitor, and JAK1 and TYK2 are not themselves known to be myristoylated (and do not possess an N-terminal Met-Gly sequence required for myristoylation^50^), the proximity of C817/C838 to a pocket that has been exploited to create allosteric drugs of another kinase encouraged us to further characterize the potential functional impact of electrophilic compounds targeting these cysteines.

### Optimization of covalent allosteric ligands that act as JAK1 inhibitors

We pursued the discovery of more potent and selective covalent ligands for JAK1_C817/TYK2_C838 by screening an internal library of electrophilic compounds using a targeted MS-ABPP assay. This approach furnished an attractive piperidine butynamide fragment hit **1a** (**Fig. 1f**) that showed target engagement values (TE_50_s) of 2.1 μM and 45 μM for JAK1_C817 and TYK2_C838, respectively (**Fig. 1g** and **Table 1**). Using a homogeneous time-resolved fluorescence (HTRF) assay in human peripheral blood mononuclear cells (PBMCs), we also found that **1a** inhibited IFNα-stimulated STAT1 phosphorylation – a JAK1/TYK2-dependent cytokine pathway – with an IC_50_ value of 1.4 μM (**Fig. 1h** and **Table 1**), suggesting that covalent ligands targeting JAK1_C817/TYK2_C838 acted as JAK1 and/or TYK2 inhibitors. The corresponding racemate of **1a** (compound **1**) was ~two-fold less active in both HTRF and TE assays (**Table 1**), indicating a stereochemical preference for engagement of JAK1_C817/TYK2_C838. We next synthesized a focused library of **1a** analogues (**Fig. 2a**) and, based on the greater potency displayed by **1a** for JAK1_C817 over TYK2_C838, screened these compounds for: i) *in vitro* engagement of JAK1_C817 (TE_50_) in human cell proteomes by targeted MS-ABPP; and ii) cell-based functional activity (IC_50_) on JAK1-dependent signaling pathways (IFNα-STAT1, IL-6-STAT3) in human PBMCs. We iteratively improved the potency of compound interactions with JAK1_C817 by three orders of magnitude and observed a strong correlation (R^2^ ~ 0.93-98) between these *in vitro* TE_50_ values and the *in situ* IC_50_ values for blocking STAT phosphorylation (**Fig. 2b, c** and **Table 1**), further supporting that the compounds acted as functional antagonists of JAK1. Key structural modifications contributing to improved potency included the addition of a second chlorine atom at the meta-position of the phenyl ring (e.g., compound **3**) and replacement of the terminal methyl of the butynamide with a pyrrolidine-methylsulfonamide (e.g., compound **5**). The tested compounds generally showed >10-fold greater potency for engagement of JAK1_C817 compared to TYK2_C838 (**Table 1**). Separation of the stereoisomers of compound **5** revealed that the enantiomers(*S, R*)**-5a** and (*R, S*)**-5b** were substantially more potent than the corresponding diastereomers (**5c** and **5d**), blocking STAT phosphorylation with IC_50_ values (~0.03-0.05 μM) that were superior to the pan-JAK inhibitor tofacitinib (**Fig. 2c** and **Table 1**). Compound **5a**, hereafter referred to as VVD-118313, was selected for further functional characterization as the compound showed the strongest overall activity across the assays.

**Table 1.** Engagement and inhibitory activity of covalent ligands targeting JAK1_C817. Engagement (TE_50_, μM, 1 h, *in vitro*) for JAK1_C817 or TYK2_C838 determined by targeted TMT-ABPP in human cell lysates. Data are mean values ± S.D. from two-three independent experiments with the exception of values marked with ^†^, which were from a single experiment. JAK1 inhibition (IC50) determined using HTRF assays measuring IFNα (100 ng/mL, 30 min)-stimulated STAT1 phosphorylation or IL-6 (25 ng/mL, 30 min)-stimulated STAT3 phosphorylation in human PBMCs pretreated with compounds for 2 h. Compounds were tested as single stereoisomers except where noted^§^. Data are mean values ± S.D. from two independent experiments except where noted (^‡^n=3, ^†^n=1). ND – not determined. NA – not applicable for a non-covalent orthosteric inhibitor.

| ID | $TE_{50}$ ($\mu M$)<br>JAK1_C817 | HTRF $IC_{50}$ ($\mu M$) | | $TE_{50}$ ($\mu M$)<br>TYK2_C838 |
| --- | --- | --- | --- | --- |
| | | IFNα-pSTAT1 | IL-6-pSTAT3 | |
| **1**<sup>§</sup> | 4.7 ± 1.3<sup>‡</sup> | 2.4 ± 0.3 | 8.9<sup>†</sup> | 93 ± 10 |
| **1a** | 2.1 ± 0.8<sup>‡</sup> | 1.2 ± 0.2 | 4.4 ± 0.1 | 45 ± 17 |
| **2a** | 0.56 ± 0.37 | 0.30 ± 0.10 | 2.7<sup>†</sup> | ND |
| **3a** | 0.071 ± 0.045<sup>‡</sup> | 0.064 ± 0.009 | 0.32 ± 0.2 | 2.8 ± 1.7 |
| **4**<sup>§</sup> | 0.16 ± 0.14<sup>‡</sup> | 0.10 ± 0.05 | 1.3 ± 1.1 | 0.65<sup>†</sup> |
| **5**<sup>§</sup> | 0.017 ± 0.004 | ND | ND | 0.28<sup>†</sup> |
| (S,R)-5a | 0.008 ± 0.002 | 0.032 ± 0.001 | 0.046 ± 0.002 | 0.21 ± 0.01 |
| (R,S)-5b | 0.016 ± 0.003 | 0.033 ± 0.001 | 0.053 ± 0.003 | 0.19<sup>†</sup> |
| (S,S)-5c | 0.092 ± 0.040 | 0.33 ± 0.30 | 2.4 ± 1.9 | 2.2<sup>†</sup> |
| (R,R)-5d | 0.077 ± 0.031 | 0.16 ± 0.06 | 1.5 ± 0.5 | 1.8<sup>†</sup> |
| Tofacitinib | NA | 0.10 ± 0.01 | 0.27 ± 0.05 | NA |

**Figure 2.**
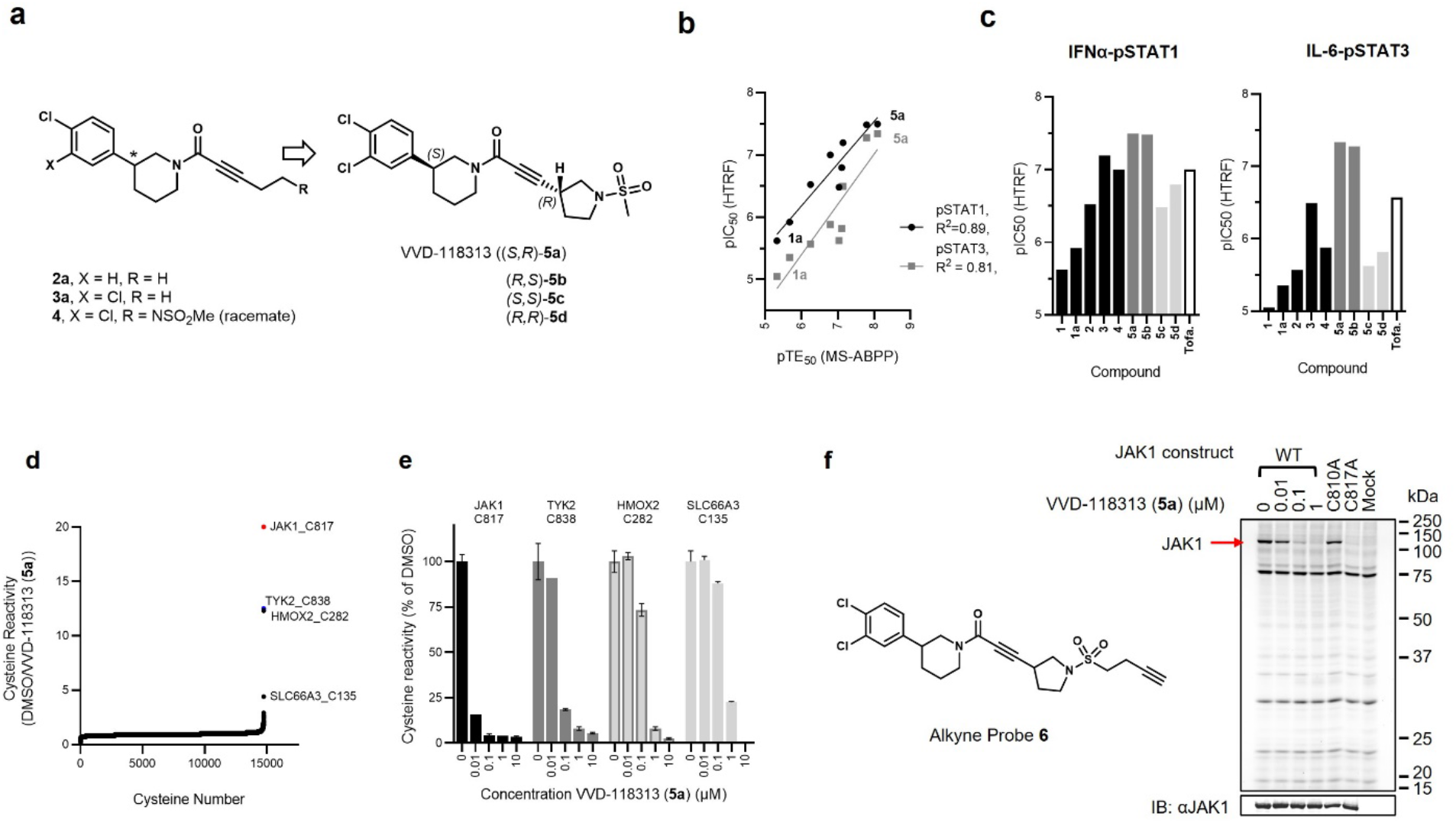
Optimization of covalent allosteric JAK1 inhibitors. **a**, Structures of VVD-118313 (compound **5a**), stereoisomers (**5b-d**), and key precursors (**2a**, **3a**, and **4**). Compounds **2a**, **3a**, and **5a-d** were tested as single stereoisomers. Absolute configuration is shown where known. **b**, Correlation between engagement of JAK1_C817 (pTE50) determined for compounds as described in **Fig. 1g** and inhibition of cytokine-induced STAT1/3 phosphorylation (pSTAT1/pSTAT3; HTRF pIC_50_) determined for compounds tested as described in **Fig. 1h** and **Table 1**. R^2^ values comparing pTE_50_ values to pIC_50_ values for pSTAT1 (black circles) and pSTAT3 (gray squares) were determined by linear regression. **c**, pIC_50_ values for inhibition of IFNα-stimulated STAT1 and IL-6-stimulated STAT3 phosphorylation for representative compounds determined as described in **Fig. 1h** and **Table 1**. For **b**, **c**, Data are mean –log-transformed values from two independent experiments. **d,** Global cysteine reactivity profile for VVD-118313 (**5a**) (1 μM, 3 h, *in situ*) in primary human PBMCs. Data represent mean ratio values (DMSO/VVD-118313) for IA-DTB-labeled, cysteine-containing peptides quantified from two replicate cell treatment experiments analyzed in a single MS-ABPP experiment. Ratio values for JAK1_C817 (red) and TYK2_C838 (blue) are highlighted. Quantified cysteines with ratios ≥ 4 (≥ 75% engagement) are marked. **e**, Concentration-dependent reactivity profiles for cysteines engaged by VVD-118313 in human PBMCs (0.01-10 μM, 3 h, *in situ*). Data are mean values from VVD-118313-treated cells shown as a percentage of DMSO-treated cells from two replicate cell treatment experiments analyzed in a single MS-ABPP experiment. **f**, *Left*, structure of alkyne probe **6**. *Right*, gel-ABPP experiment showing labeling of recombinant WT-JAK1 and C810A-JAK1, but not C817A-JAK1, expressed in 22Rv1 cells with alkyne probe **6** (0.1 μM, 2 h, *in situ*). The labeling of WT-JAK1 is blocked by pretreatment with VVD-118313 (0.01-1 μM, 2 h, *in situ*). Below, western blot showing JAK1 expression in gel-ABPP experiment. Data are from a single experiment representative of two independent experiments.

### VVD-118313 selectively inhibits JAK1 through engagement of C817

We next evaluated the broader proteomic reactivity of VVD-118313 by untargeted MS-ABPP in human PBMCs. Across >14,000 quantified cysteines, JAK1_C817 was the most potently engaged site by VVD-118313 (0.01 – 10 μM, 3h), followed by TYK2_C838, with both cysteines showing near-complete blockade in their IA-DTB reactivity in cells treated with 0.1 μM of VVD-118313 (**Fig. 2d, e** and **Supplementary Dataset 1**). Two additional cysteines (HMOX2_C282, SLC66A3_C135) were engaged by VVD-118313 when tested at a 10-fold higher concentration (1 μM; **Fig. 2d, e**). Similar results were obtained in MS-ABPP experiments that analyzed the *in vitro* proteome-wide reactivity of VVD-118313 in PBMC lysates, where JAK1_C817 was again the most potently engaged cysteine, followed by TOR4A_C21, a site that was also engaged *in situ*, albeit more weakly, and TYK2_C838 (**Extended Data Fig. 2** and **Supplementary Dataset 1**). Taken together, these chemical proteomic data support that VVD-118313 is a highly potent and selective covalent ligand for JAK1_C817 and TYK2_C838.

To test whether VVD-118313 inhibits JAK1 through engagement of C817, we recombinantly expressed WT-JAK1 and a C817A-JAK1 mutant in the 22Rv1 human prostate cancer cell line, which has a frameshift mutation in the *JAK1* gene and consequently only expresses JAK2 and TYK2 (JAK3 is immune cell-restricted in its expression)^3^. We also evaluated a C810A-JAK1 mutant, as the quantified C817 tryptic peptide in our MS-ABPP experiments also contained C810, a residue that is not conserved in TYK2 (**Fig. 1d**) and is further away than C817 from the pocket predicted to bind electrophilic compounds (**Fig. 1e**). We first treated 22Rv1 cells expressing the JAK1 variants with an alkynylated analogue of VVD-118313 (alkyne probe **6** (0.1 μM, 2 h); **Fig. 2f**) and, after cell lysis, detected **6**-labeled proteins by copper-catalyzed azide-alkyne cycloaddition (CuAAC)^51^ with a rhodamine (Rh)-azide reporter group, followed by SDS-PAGE and in-gel fluorescence scanning. Alkyne probe **6** reacted with WT- and C810A-JAK1, but not C817A-JAK1, and the labeling of WT-JAK1 was blocked in a concentration-dependent manner by pre-treatment with VVD-118313 (**Fig. 2f**). We interpret these data to indicate that VVD-118313 site-specifically engages JAK1 at C817.

JAK1 mediates STAT phosphorylation downstream of different cytokine receptors by heterodimerizing with other JAK family members (**Fig. 3a**)^3^. We selected a representative subset of these pathways (IFNα-STAT1 and IL-6-STAT3), along with a JAK2-mediated pathway (prolactin (PRL)-STAT5), to evaluate the functional effects of VVD-118313 on the activity of recombinantly expressed WT and mutant forms of JAK1 in 22Rv1 cells. We first verified that recombinant WT-JAK1 and the C810A and C817A mutants equivalently rectified intrinsic defects in IFNα and IL-6 signaling in parental 22Rv1 cells^37,52^, as reflected by the greater IFNα or IL-6-stimulated STAT1/3 phosphorylation in cells expressing these JAK1 variants compared to mock cells (**Fig. 3b** and **Extended Data Fig. 3a**). We also noted that all of the JAK1 variant-expressing cells displayed a similar degree of constitutive phosphorylation of the JAK1 JH1 activation loop (Y1034/Y1035) that was not further increased by cytokine treatment (**Fig. 3b**). VVD-118313 (2 μM, 2 h) blocked IFNα-simulated STAT1 and IL-6-stimulated STAT3 phosphorylation in WT- or C810A-JAK1-expressing 22Rv1 cells, but not in C817A-JAK1-expressing cells (**Fig. 3b, c**). In contrast, the orthosteric JAK inhibitor tofacitinib equivalently blocked IFNα-simulated STAT1 phosphorylation in cells expressing WT-, C810A-, or C817A-JAK1 (**Fig. 3b, c**). Interestingly, VVD-118313 also completely blocked the constitutive phosphorylation of WT- and C810A-JAK1 but did not affect the phosphorylation of C817A-JAK1 (**Fig. 3b, d**). In contrast, tofacitinib only partly (~50%) reduced phosphorylation of all JAK1 variants (**Fig. 3b, d**). VVD-118313 and tofacitinib further differed in their effects on JAK2-mediated signaling, where VVD-118313 was inactive, while tofacitinib fully inhibited PRL-induced STAT5 phosphorylation (**Fig. 3b, c**). Concentration-dependent analyses revealed that VVD-118313 maximally inhibited IFNα-STAT1 and IL-6-STAT3 phosphorylation (> 80% in each case) in WT- or C810A-JAK1-expressing 22Rv1 cells at ~0.2 μM, while showing negligible impact (< 10%) in cells expressing C817A-JAK1 up to 2 μM (**Fig. 3e** and **Extended Data Fig. 3b, c**). VVD-118313 inhibited WT- and C810A-JAK1 phosphorylation with even greater potency than STAT1/STAT3 phosphorylation, showing maximal activity (> 90% blockade) at 0.05 μM (**Fig. 3e**), which could indicate that only a small fraction of residually phosphorylated and activated recombinant JAK1 is required to support signal transduction in IFNα/IL-6-stimulated 22Rv1 cells. Together, these data indicate that VVD-118313 acts as a selective allosteric inhibitor of JAK1 through covalent engagement of C817.

**Figure 3.**
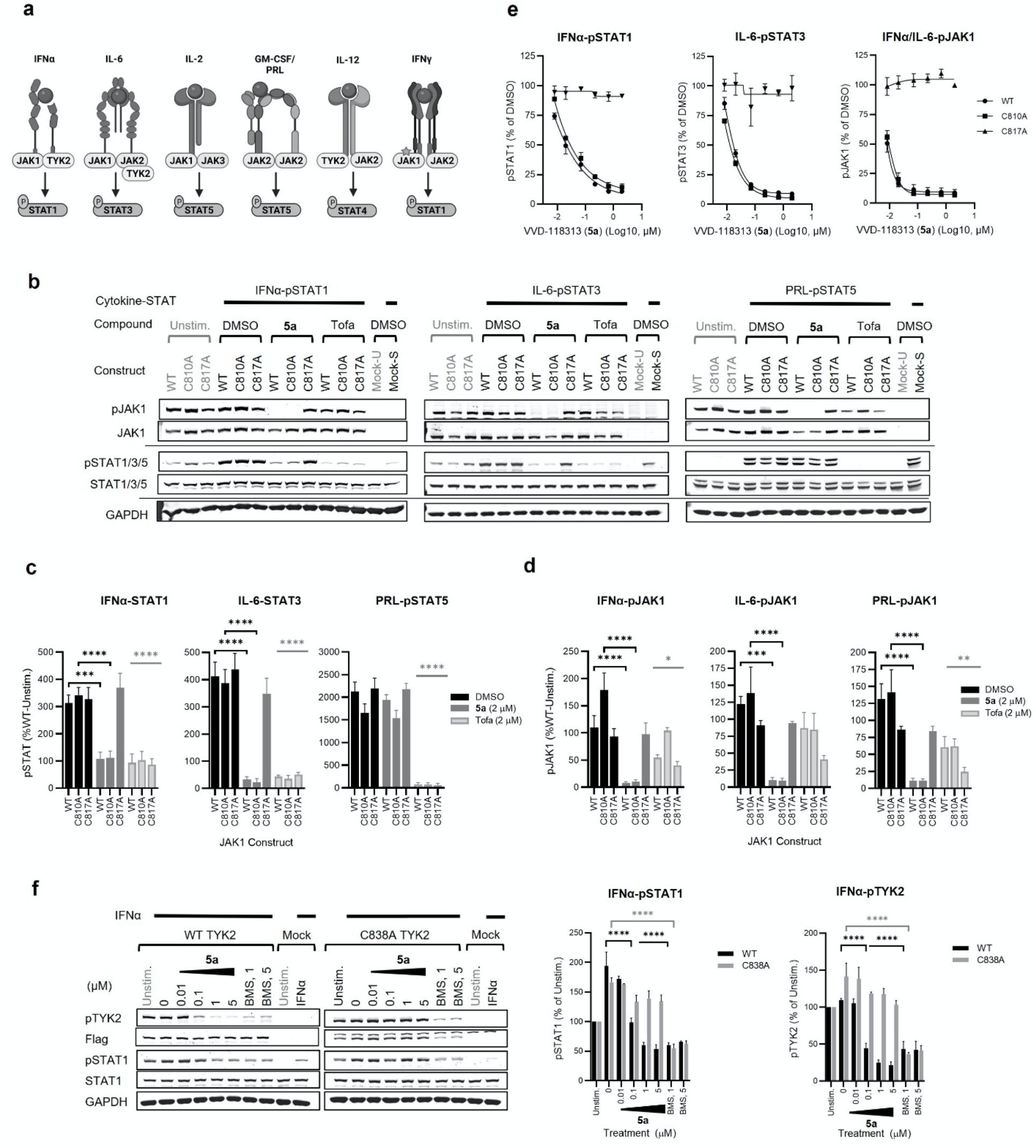
VVD-118313 inhibits JAK1 through engagement of C817. **a,** Representative cytokine signaling pathways involving different JAK family members. *JAK1 serves a scaffolding role for IFNg-STAT1 signaling^31,38,54^. **b**, Western blots showing effects of VVD-118313 (**5a**) and the pan-JAK inhibitor tofacitinib (Tofa) on JAK1 phosphorylation (pJAK1; Y1034/Y1035 phosphorylation detected with (D7N4Z) Rabbit mAb #74129, CST) and IFNα-stimulated STAT1 (JAK1-dependent), IL-6-stimulated STAT3 (JAK1-dependent), and prolactin (PRL)-stimulated STAT5 (JAK2-dependent) phosphorylation in 22Rv1 cells expressing WT-, C810A-, or C817A-JAK1. Cells were treated with compounds (2 μM) for 2 h and then stimulated with IFNα (100 ng/mL, 30 min), IL-6 (50 ng/mL, 30 min) or PRL (15 ng/mL, 15 min) prior to analysis. **c**, **d**, Quantification of pSTAT1/3/5 (**c**) and pJAK1 (**d**) signals from (**b**). Signal intensities were normalized relative to the unstimulated WT-JAK1-transfected control for each experiment. Data are mean values ± S.E.M. from three independent experiments. Significance was determined by two-way ANOVA with Dunnett’s post-hoc test and reported relative to stimulated, DMSO-treated control of respective JAK1 construct. ****P<0.0001, ***P<0.001, **P<0.01, *P<0.05. **e**, Concentration-dependent effects of VVD-118313 (**5a**) on IFNα-stimulated pSTAT1 (left), IL-6-stimulated pSTAT3 (middle), and pJAK1 (integrated from both IFNα- and IL-6-stimulations) in 22Rv1 cells expressing WT-JAK1. Data are mean values ± SEM from two (pSTAT1, pSTAT3) or three (pJAK1) independent experiments. See **Extended Data Fig. 3b, c** for corresponding western blots. **f**, Concentration-dependent effects of VVD-118313 (**5a**; 0.01 – 5 μM, 2 h) and BMS-986165 (BMS, 1 or 5 μM, 2 h) on TYK2 phosphorylation (pTYK2) and IFNα-stimulated STAT1 phosphorylation in 22Rv1 cells expressing recombinant WT-TYK2 or a C838A-TYK2 mutant. *Left*, representative western blots. *Middle and right*, quantification of pSTAT1 (middle) and pTYK2 (right) signals normalized to unstimulated control cells expressing WT-TYK or C838A-TYK2. Data are mean values ± S.E.M from three independent experiments. Significance was determined by two-way ANOVA with Dunnett’s post-hoc test. P-values are only shown for the lowest concentration of each compound that displayed significance for inhibition of pSTAT1 and pTYK2. ****P<0.0001, ***P<0.001.

To determine if VVD-118313 also inhibited TYK2 signaling, we first confirmed that alkyne probe **6** reacted with recombinantly expressed WT-TYK2, but not a C838A-TYK2 mutant, in 22Rv1 cells, and that pre-treatment with VVD-118313 blocked compound **6** reactivity with WT TYK2 (**Extended Data Fig. 4a**). We did not observe a signal for endogenous TYK2 in these experiments (e.g., in mock 22Rv1 cells), indicating that the expression level of TYK2 in 22Rv1 cells was too low to visualize with alkyne probe **6**. 22Rv1 cells expressing recombinant WT- or C838A-TYK2 displayed modestly increased IFNα-induced STAT1 phosphorylation compared to mock 22Rv1 cells (**Extended Data Fig. 4b**), suggesting that atypical TYK2 homodimers and/or TYK2-JAK2 heterodimers can partly support IFNα signaling in these cells^37,52,53^. VVD-118313 (0.01 - 5 μM, 2 h) inhibited IFNα-STAT1 phosphorylation in 22Rv1 cells expressing WT-TYK2, but not C838A-TYK2, while the TYK2 inhibitor BMS-986165 blocked IFNα-STAT1 phosphorylation in both cell populations (**Fig. 3f**). We further noted that VVD-118313 and BMS-986165 blocked the weaker IFNα/IL-6-stimulated STAT1/STAT3 signaling in mock 22Rv1 cells, which lack JAK1 expression (**Extended Data Fig. 4c-f**) suggesting that these pathways are mediated by endogenous TYK2. Similar to what we observed for JAK1, VVD-118313 inhibited phosphorylation of the activation loop of WT-, but not C838A-TYK2 (**Fig. 3f**). BMS-986165 also suppressed TYK2 phosphorylation; however, this effect was independent of C838 (**Fig. 3f**). These data support that VVD-118313 can site-specifically inhibit the signaling of TYK2, at least in the context of a cell line where JAK1 is absent.

### VVD-118313 selectively inhibits JAK1 signaling in primary immune cells

We next evaluated the activity of VVD-118313 (**5a**) and its mixture of stereoisomers (compound **5**) in primary human immune cells and found that both compounds inhibited IFNα-pSTAT1, IL-6-pSTAT3, and IL-2-pSTAT5 pathways in a concentration-dependent manner (**Fig. 4a-c**). At concentrations of 0.1 and 1 μM - where VVD-118313 fully engaged JAK1_C817 in human PBMCs (**Fig. 2d, e**) and substantially blocked IFNα, IL-6, and IL-2 signaling (**Fig. 4a-c**) - no effects on GM-CSF/JAK2-mediated STAT5 phosphorylation were observed (**Fig. 4d**). When tested at 10 μM, VVD-118313 showed modest inhibitory effects on GM-CSF/JAK2-STAT5 phosphorylation (**Fig. 4d**). Considering that 10 μM of VVD-118313 is a 100-fold greater concentration than that required to fully engage JAK1_C817, we interpret the modest inhibitory activity on GM-CSF/JAK2-STAT5 phosphorylation to reflect an off-target activity and note that several cysteines in the human PBMC proteome were substantially engaged by this compound at 10 μM (**Extended Data Fig. 5a** and **Supplementary Dataset 1**). VVD-118313 showed the strongest inhibitory effect on IFNα-stimulated STAT1 phosphorylation (>50% and >90% inhibition at 0.01 and 0.1 μM, respectively), followed by IL-6-pSTAT3 (~80-85% inhibition at 0.1-1 μM) and IL-2-pSTAT5 (~70% inhibition at 0.1-1 μM) pathways. Compound **5** behaved similarly to VVD-118313, with the expected reduction in potency (**Fig. 4a-d**) that is in accordance with TE50 values measured by MS-ABPP (**Table 1**). The pan-JAK inhibitor tofacitinib blocked all of the evaluated cytokine-JAK/STAT pathways at a concentration of 1-2 μM (**Fig. 4a-d**). Finally, we found that VVD-118313 did not block TYK2-dependent IL-12-STAT4 signaling in human PBMC-derived T-blasts, which was inhibited by both BMS-986165 and tofacitinib (**Fig. 4e**). This result differed from the inhibitory activity displayed by VVD-118313 in JAK1-null 22Rv1 cells, where the compound suppressed TYK2-dependent STAT1 phosphorylation and suggests that under more physiological settings, VVD-118313 does not act as a functional antagonist of TYK2.

**Figure 4.**
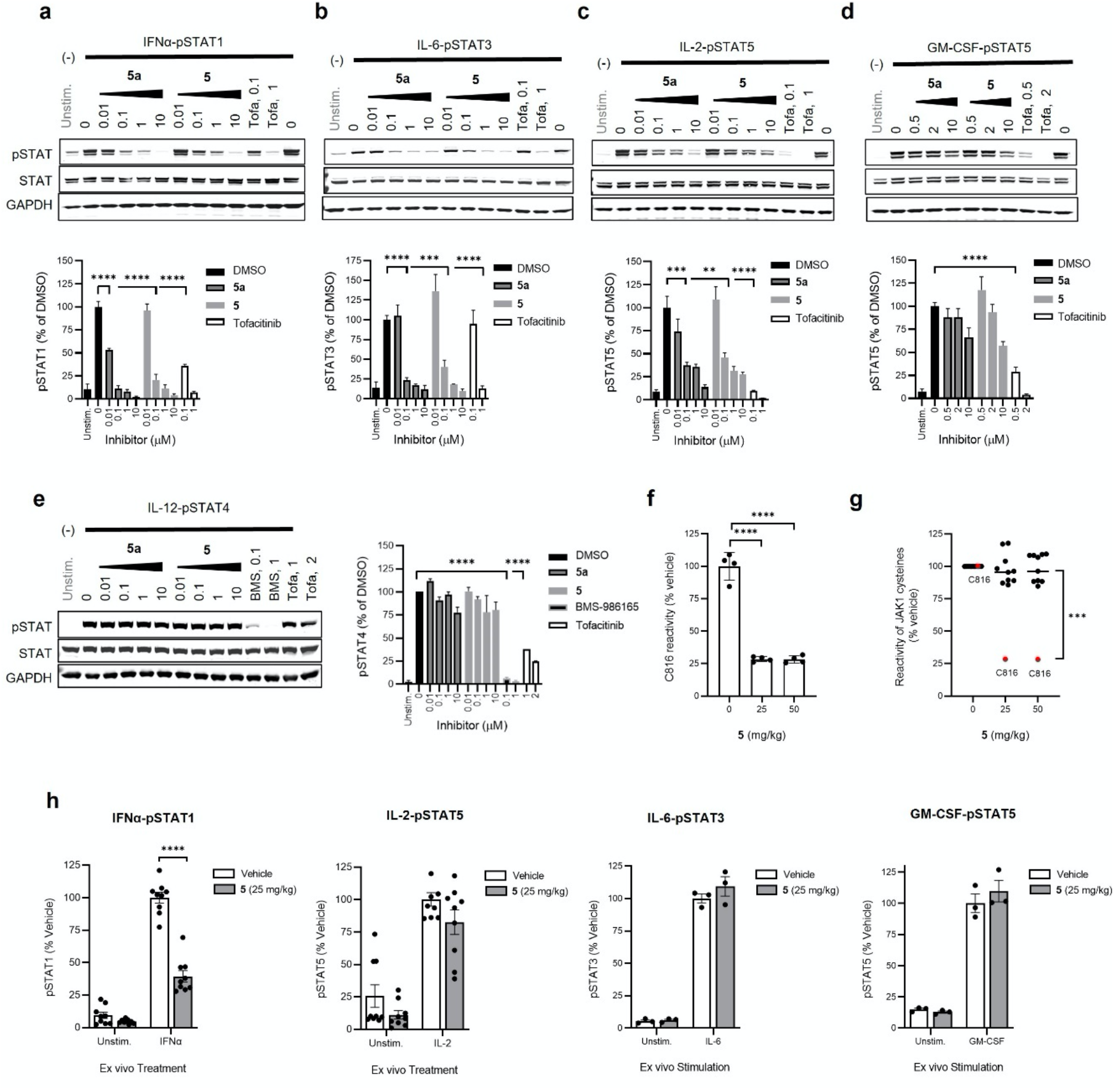
VVD-118313 selectively inhibits JAK1 signaling in primary human immune cells and mice. **a-d**, Effects of VVD-118313 (**5a**), stereoisomeric mixture **5**, and tofacitinib (Tofa) on JAK-STAT signaling pathways in human PBMCs. PBMCs were treated with compounds at the indicated concentrations for 2 h prior to stimulation with IFNα (**a**; 100 ng/mL, 30 min), IL-6 (**b**; 25 ng/mL, 30 min), IL-2 (**c**; 20 U/mL, 15 min), or GM-CSF (**d**; 0.5 mg/mL, 15 min). *Upper*, representative western blots. *Lower*, Quantification of pSTAT signal intensities shown as a percent of the stimulated DMSO-treated control cells for each assay. Data are mean values ± S.E.M. from three (IL-6, IL-2) or four (IFNα, GM-CSF) independent experiments. Significance determined by one-way-ANOVA with Dunnett’s post-hoc test. P-values are only shown for the lowest concentration of each compound that displayed significance for inhibition of pSTAT. All higher concentrations were similarly statistically significant. ****P<0.0001, ***P<0.001, **P<0.01, *P<0.05. **e**, Effects of VVD-118313 (**5a**), compound **5**, BMS-986195 (BMS), and tofacitinib (Tofa) on IL-12-stimulated STAT4 phosphorylation in phytohemagglutinin (PHA-P)/IL-2-generated PBMC-derived T-blasts. Cells were treated with compounds at the indicated concentrations for 2 h prior to stimulation with IL-12 (12.5 ng/mL, 15 min). Data are mean values ± S.E.M. normalized to DMSO controls from three independent experiments and significance was determined as for (**a-d**). **f**, **g**, Reactivity profiles for JAK1_C816 (**f**) and all quantified JAK1 cysteines (**g**) from proteomic lysates of spleen tissue from mice treated with vehicle (dose 0 mg/kg) or compound **5** (25 or 50 mg/kg, s.c. 2 × 4 h). Data are mean values ± S.D. from compound **5**-treated mice shown as a percentage of vehicle-treated mice (n = 4 animals/group analyzed in a single targeted MS-ABPP). In **g**, bars represent median reactivity values for all JAK1 cysteines other than C816. Significance determined by one-way ANOVA with Dunnett’s post hoc test. ****P<0.0001. **h**, *Ex vivo* cytokine stimulation of splenocytes from mice treated with vehicle or **5** (25 mg/kg, s.c., 2 × 4 h); IFNα (100 ng/mL, 30 min), IL-2 (20 U/mL, 15 min), IL-6 (10 ng/mL, 30 min) or GM-CSF (10 ng/mL, 15 min). Data are mean values ± S.E.M., from three (IFNα, IL-2) or one (IL-6, GM-CSF) independent experiments, each containing n = 3 mice per treatment group. Significance determined by two-way ANOVA with Šidák’s post hoc test. ****P<0.0001.

We were also interested in evaluating whether the covalent allosteric inhibitors were capable of engaging and inhibiting JAK1 *in vivo*. We first confirmed by MS-ABPP that VVD-118313 selectively and completely engaged C816 of mouse JAK1 (the corresponding residue to human JAK1_C817) in splenocyte proteomic lysates at concentrations as low at 0.01 μM (**Extended Data Fig. 5b** and **Supplementary Dataset 1**). Across >11,000 quantified sites, only a single additional cysteine BACH1_C438 was engaged > 70% by 1 μM of VVD-118313. Mouse TYK2 was not targeted by VVD-118313 because this protein possesses a serine residue (S858) in the position corresponding to human TYK2_C838. As we observed in human immune cells, VVD-118313 and compound **5** inhibited both IFNa-dependent STAT1 and IL-2-dependent STAT5 phosphorylation in mouse splenocytes at concentrations as low at 0.01-0.1 μM (**Extended Data Fig. 6a, b**). In contrast, VVD-118313 produced only minimal effects (< 30%) on GM-CSF-STAT5 or IL-12-STAT4 phosphorylation signaling up to 2 μM test concentration (**Extended Data Fig. 6c, d**). One unexpected observation was that VVD-118313 did not inhibit IL-6-STAT3 signaling in mouse splenocytes (**Extended Data Fig. 6e**), which contrasted with the robust inhibition of this pathway observed in human PBMCs (**Fig. 4b**). Since tofacitinib retained inhibitory activity against IL-6-STAT3 signaling in mouse splenocytes (**Extended Data Fig. 6e, h**), we speculate that this cytokine pathway may be regulated by different JAK subtypes in human PBMCs versus mouse splenocytes. In support of this hypothesis, we found that BMS-986165 suppressed IL-6-stimulated STAT3 phosphorylation in mouse splenocytes to an even greater degree than the pan-JAK inhibitors tofacitinib and upadacitinib (**Extended Data Fig. 6f**), suggesting that TYK2 may contribute more substantially than JAK1 to IL-6-stimulated STAT3 phosphorylation in mouse splenocytes.

We next performed *in vivo* studies using compound **5**, which was chosen over VVD-118313 due to the comparative ease of scaled-up synthesis and the comparable functional activity of the mixture of stereoisomers in primary immune cells (**Fig. 4a-d**). Initial pharmacokinetic studies revealed that compound **5** exhibited a short half-life (0.36 h) and rapid clearance in mice (112 mL/min/kg) (**Extended Data Table 1**). Nonetheless, we hypothesized that the covalent mechanism of action of **5** may overcome its suboptimal pharmacokinetic properties to still allow for substantial engagement of JAK1_816 *in vivo*. The compound was administered subcutaneously to mice at 25 or 50 mg/kg (or vehicle control) in a two-dose protocol with a 4 h interval between doses. At 4 h after the second dose, mice were sacrificed and spleen tissue analyzed by targeted MS-ABPP, which confirmed ~75% engagement of JAK1_C816 at both 25 and 50 mg/kg of compound **5**, while other cysteines in JAK1 were unaffected (**Fig. 4f, g**). We also found that splenocytes from compound **5**-treated mice showed substantial impairments in IFNa-dependent STAT1 phosphorylation compared to splenocytes from vehicle-treated mice (**Fig. 4h** and **Extended Data Fig 7**). In contrast, IL-2-dependent STAT5 phosphorylation, which was incompletely blocked by VVD-118313 or compound **5** in cultured human (**Fig. 4c**) and mouse (**Extended Data Fig. 6b**) immune cells, was not substantially altered in splenocytes from compound **5**-treated mice (**Fig. 4h** and **Extended Data Fig 7**), suggesting that insufficient JAK1 engagement occurred *in vivo* to impact this pathway. Finally, consistent with our cultured immune cell studies, IL-6-STAT3 and GM-CSF-STAT5 signaling were unaffected in splenocytes isolated from compound **5**-treated mice (**Fig. 4h** and **Extended Data Fig 7**).

Taken together, our data indicate that covalent ligands engaging human JAK1_C817 (or mouse JAK1_C816) selectively disrupt JAK1-dependent cytokine signaling in human and mouse immune cells and can serve as chemical probes for both cellular and *in vivo* studies.

### VVD-118313 blocks JAK1 transphosphorylation in a C817-dependent manner

The more extensive blockade of JAK1 phosphorylation by covalent allosteric inhibitors compared to orthosteric inhibitors (**Fig. 3c**) pointed to distinct mechanisms of action for each class of compounds. Also consistent with this premise, VVD-118313 did not inhibit the catalytic activity of recombinant purified JAK1 (aa 438 - 1154, J01-11G, SignalChem) in a peptide substrate assay (VA7207, Promega), whereas tofacitinib displayed a strong inhibitory effect (**Fig. 5a**). We next explored the potential mechanistic basis for VVD-118313 blockade of JAK1 phosphorylation by evaluating this compound in 22Rv1 cells co-expressing differentially epitope-tagged catalytically active (FLAG-tagged WT or C817A) or inactive (HA-tagged K908E or C817A/K908E) variants of JAK1. We first found that, in the absence of VVD-118313, WT-JAK1 - FLAG and C817A-JAK1-FLAG were robustly auto-phosphorylated when individually expressed in 22Rv1 cells, while the K908E-JAK1-HA and C817A/K908E-JAK1-HA variants did not show evidence of phosphorylation (**Fig. 5b**). Co-expression with either catalytically active JAK1-FLAG variant (WT or C817A) led to clear *trans*-phosphorylation of either inactive JAK1-HA variant (K908E or C817A/K908E), although the magnitude of this trans-phosphorylation activity was noticeably higher in cells expression WT-JAK1-FLAG versus C817A-JAK1-FLAG and weakest in cells co-expressing both C817A-JAK1-FLAG and C817A/K908E-JAK1-HA (**Fig. 5b, c**). VVD-118313 (2 μM, 2 h) completely blocked trans-phosphorylation of either inactive JAK1-HA variant (K908E or C817A/K908E) in cells expressing active WT-JAK1-FLAG, but did not affect transphosphorylation of inactive JAK1-HA variants in cells expressing active C817A-JAK1-FLAG (**Fig. 5b, d**). We also found that BMS-986165, which has been shown to bind the JAK1 JH2 domain *in vitro* (IC_50_ ~ 1 nM) and inhibit JAK1-dependent cytokine signaling (IL-6 and IL-2) with an IC50 of ~ 0.5 μM^21^, blocked JAK1 *trans*-phosphorylation in a C817-independent manner (**Extended Data Fig. 8a**). We interpret these data to indicate that the inhibition of *trans*-phosphorylation of JAK1 contributes to the allosteric mechanism of action of VVD-118313 and that this effect required the presence of C817 on the donor (phosphorylating), but not the recipient (phosphorylated) JAK1 variant. That the C817A mutation, on its own, partly suppressed trans-phosphorylation (**Fig. 5b, c**), despite not impairing cytokine-induced STAT phosphorylation (**Fig. 3c**) can be further interpreted to suggest: i) the potential for allosteric communication between the C817 pocket of the JH2 domain and the JH1 kinase domain of JAK1, even in the absence of exogenous inhibitor; and ii) only a modest fraction of phosphorylated JAK1 is required to fully support cytokine-dependent signaling in the 22Rv1 cell system, as noted above (**Fig. 3b-e**).

**Figure 5.**
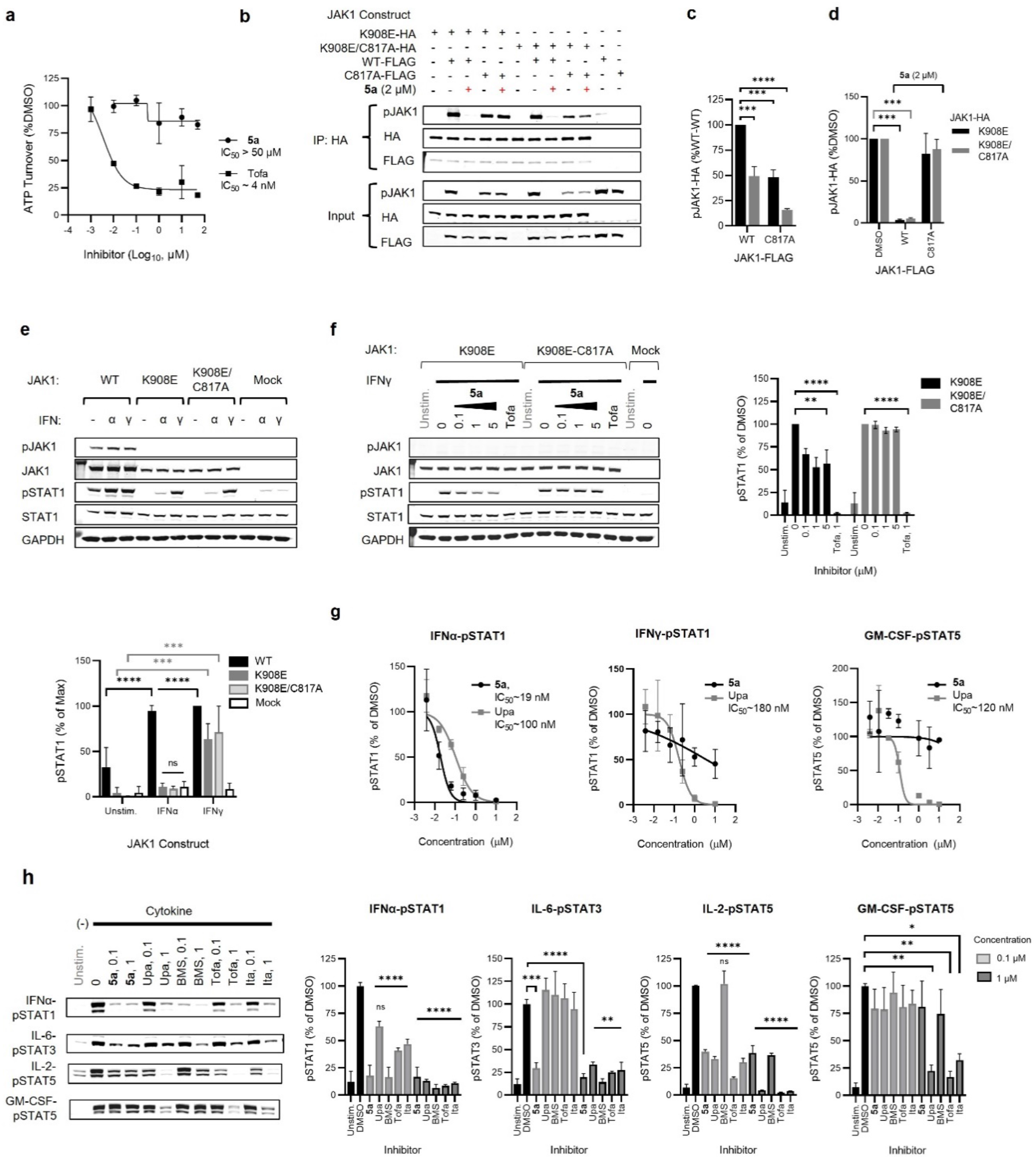
Mechanistic properties and distinct activity profile of allosteric JAK1 inhibitors. **a**, Substrate assay quantifying ATP turnover by recombinant purified JAK1 (residues 438-1154 fused to GST) treated with DMSO, VVD-118313 (**5a**) or tofacitinib (Tofa) (0.001 - μM, 30 min) prior to addition of an IRS-1 peptide substrate (0.2 μg/mL) and ATP (50 μM, 1 h). Data are mean values ± S.D. from two independent experiments. **b**, Western blots measuring JAK1 phosphorylation (pJAK1) from anti-HA immunoprecipitations (IPs) of HA-tagged kinase dead (K908E) JAK1 (WT or C817A mutant) expressed in 22Rv1 cells alongside catalytically active FLAG-tagged JAK1 (WT or C817A mutant). **c**, **d**, Quantification of pJAK1 signals from anti-HA-IPs shown in **b**. Panel **c** shows pJAK1 signals from anti-HA IPs from DMSO-treated cells coexpressing the indicated combinations of K908E-JAK1-HA or C817A/K908E-JAK1-HA with WT-JAK1-FLAG or C817A-JAK1-FLAG. Panel **d** shows pJAK1 signals for the same anti-HA IPs from VVD-118313 (**5a**; 2 μM, 2 h)-treated 22Rv1 cells. Data in **c** are normalized to signals in 22Rv1 cells expressing K908E-JAK1-HA and WT-JAK1-FLAG. Data in **d** are normalized to signals in DMSO-treated control cells Data for **c** and **d** are mean values ± S.E.M. from three independent experiments. Significance was determined by two-way ANOVA with Dunnett’s post-hoc test. ***P<0.001, ****P<0.0001. **e,** *Upper*, western blots showing that both K908E- and K908E/C817A-JAK1 mutants support IFNγ-stimulated (50 ng/mL, 30 min), but not IFNα-stimulated (100 ng/mL, 30 min) STAT1 phosphorylation (pSTAT1) in 22Rv1 cells. WT-JAK1 supports both cytokine pathways. *Lower*, quantification of western data. Data are mean values ± S.E.M. from three independent experiments. pSTAT1 signals were normalized to the maximum signal, which was generated by IFNγ-stimulated WT-JAK1 transfected cells. Significance was determined by two-way ANOVA with Tukey’s post-hoc test. ****P<0.0001, ***P<0.001. ns – no significance difference between IFNα-stimulated K908E-JAK1-HA- or K908E/C817A-JAK1-HA-expressing cells and mock cells. **f**, *Left*, western blots showing the effects of VVD-118313 (**5a**; 0.1-5 μM, 2 h) and tofacitinib (Tofa; 1 μM, 2 h) on IFNγ-dependent STAT1 phosphorylation (pSTAT1) in 22Rv1 cells expressing K908E-JAK1-HA or K908E/C817A-JAK1-HA. *Right*, quantification of western data. Data are mean values ± S.E.M. from three independent experiments. Significance determined by two-way ANOVA with Dunnett’s post-hoc test. *P<0.05, **P<0.01, ****P<0.0001, ns – non-significant. **g**, Concentration-dependent effects of VVD-118313 (**5a**) or upadacitinib (Upa) on IFNα-dependent STAT1, IFNγ-dependent STAT1, and GM-CSF-dependent STAT5 phosphorylation in human PBMCs. pSTAT signals were normalized to the DMSO-treated cytokine-stimulated control in each assay. Dose-response curves are mean values ± S.D. of two biological replicates used to estimate IC50 values by fitting data to a 4PL model. **h**, *Left*, western blots comparing the effects of VVD-118313 (**5a**) and a panel of JAK inhibitors on the indicated cytokine-stimulated pSTAT pathways in human PBMCs. *Right*, quantification of western data. Data are mean values ± S.E.M. from two (IL-6) or three (IFNα, IL-2, GM-CSF) independent experiments. Significance determined by one-way ANOVA with Šidák’s post-hoc test. *P<0.05, ****P<0.0001, ns – non-significant

### VVD-118313 shows a distinct functional selectivity profile among JAK inhibitors

In addition to supporting cytokine signaling through phosphorylation of downstream substrates (e.g., STATs), JAK kinases have also been found to play scaffolding roles in these signal transduction pathways. For instance, both JAK1 and JAK2 participate in the IFNγ-STAT1 pathway, but only the catalytic activity of JAK2 is required for STAT1 phosphorylation, while the JAK1 pseudokinase domain is thought to serve a scaffolding function^31,54^. We confirmed that IFNγ promoted STAT1 phosphorylation in 22Rv1 cells expressing recombinant WT-, K908E-, or C817A/K908E-JAK1, but not in mock-transfected cells (**Fig. 5e**). In contrast, IFNα-stimulated STAT1 phosphorylation was only supported by catalytically active WT-JAK1, but not the K908E or C817A/K908E mutants (**Fig. 5e**). We found that VVD-118313 produced an ~40% partial blockade of IFNγ-stimulated STAT1 in K908E-JAK1-expressing 22Rv1 cells, but not in C817A/K908E JAK1-expressing cells (**Fig. 5f**). In contrast, tofacitinib (1 μM, 2h) fully inhibited IFNγ-stimulated STAT1 phosphorylation in 22Rv1 cells expressing either K908E-JAK1 or C817A/K908E-JAK1 (**Fig. 5f**), which presumably reflects the blockade of endogenous JAK2 activity by this pan-JAK inhibitor. We further evaluated the impact of VVD-118313 on JAK1 scaffolding function in primary human immune cells. We specifically compared the concentration-dependent effects of VVD-118313 and the JAK1/JAK2 inhibitor upadacitinib^10,55^ on IFNα-STAT1, IFNγ-STAT1, and GM-CSF-STAT5 signaling pathways in human PBMCs, which revealed that upadacitinib blocked all three pathways with similar efficacy (> 90%) and potency (IC50 values of ~0.10-0.18 μM), while VVD-118313 fully blocked IFNα-mediated phosphorylation of STAT1 with an IC_50_ value of ~0.02 μM, partially blocked IFNγ-mediated phosphorylation of STAT1 with an Imax of ~50%, and did not inhibit GM-CSF-mediated phosphorylation of STAT5 (**Fig. 5g** and **Extended Data Fig. 8b**).

To further explore the distinct pharmacological profile of VVD-118313, we compared the compound to other JAK inhibitors in a panel of cytokine-induced STAT phosphorylation assays in human PBMCs. Consistent with our other results (**Fig. 4a-d**), VVD-118313, at both test concentrations (0.1 and 1 μM), produced a near complete blockade of JAK1-dependent IFNα-STAT1 and IL-6-STAT3 signaling and a partial inhibition of IL-2-STAT5 signaling, while showing negligible effects on JAK2-dependent GM-CSF-STAT5 signaling (**Fig. 5h**). This profile was differentiated from the other tested JAK inhibitors – tofacitinib, upadacitinib or itacitinib – all of which showed weaker potency than VVD-118313 in the IFNα-STAT1 and IL-6-STAT3 signaling assays, but much greater activity in the IL-2-STAT5 and GM-CSF-STAT5 assays (**Fig. 5h** and **Extended Data Fig. 8c**). The allosteric TYK2 inhibitor BMS-986165 displayed its greatest potency, as expected, in suppressing IFNα-STAT1 signaling, followed by IL-6-STAT3 and IL-2-STAT5 signaling, while being inactive against GM-CSF-STAT5 signaling (**Fig. 5h** and **Extended Data Fig. 8c**). Finally, we attempted to measure the effects of VVD-118313 and other JAK inhibitors on JAK1 phosphorylation, but the signals were too low to visualize in human PBMCs.

Taken together, our studies in primary immune cells illuminate a unique pharmacological profile of allosteric covalent ligands engaging JAK1_C817 compared to other JAK inhibitors that includes: i) the strong blockade of cytokine pathways, like IFNα-STAT1 and IL-6-STAT3 signaling, that depend on the catalytic functions of JAK1; ii) the partial inhibition of cytokine pathways that depend on the scaffolding functions of JAK1 (IFNγ-STAT1) or the activity of multiple JAK subtypes (IL-2-STAT5^36^); and iii) the sparing of cytokine pathways that depend predominantly on the activity of other JAK subtypes (GM-CSF-STAT5 (JAK2), IL-12-STAT4 (TYK2)).

## Discussion

Despite the potential benefits afforded by allosteric over orthosteric kinase inhibitors, which include not only improvements in selectivity due to interactions with less conserved pockets, but also avoidance of direct ATP competition for binding, the identification of ligandable and functional allosteric sites remains challenging^56–58^. Allostery is often context-dependent and, therefore, may not be detected in more conventional high-throughput assays with purified kinases and simple peptide substrates, especially if these assays only use truncated catalytic domains^59^. Existing allosteric kinase inhibitors have largely been discovered serendipitously or with detailed knowledge of endogenous regulatory mechanisms^56–58^. Examples include the “type III” inhibitors of MEK1/2^60^, LIMK^61^, and AKT^62^, which bind to a pocket adjacent to the ATP-binding site, and the “type IV” inhibitors of Bcr-Abl^45,46,63,64^, MAPKs^65^, and receptor tyrosine kinases^66,67^, which bind to allosteric sites distal to the ATP-binding pocket. We have shown here that chemical proteomics offers a distinct way to discover allosteric inhibitors of kinases.

The profiling of simple electrophilic fragments provided initial evidence of covalent ligandability of a cysteine that is found in the JH2 pseudokinase domain of JAK1 (C817) and TYK2 (C838), but not in JAK2 or JAK3. This insight was then efficiently progressed to inhibitory chemical probes by the coordinated use of targeted chemical proteomic and cell-based functional assays, furnishing an advanced compound VVD-118313 that site-specifically engages and inhibits JAK1 in primary immune cells with low-nanomolar potency, while showing negligible effects on JAK2-dependent signaling pathways. Despite also engaging TYK2_C838 at a concentration < 1 μM, VVD-118313 had a more subtle functional impact on this kinase, as we did not observe inhibition of TYK2-dependent cytokine signaling by this compound in human immune cells. These data indicate that VVD-118313 acts as a functionally selective JAK1 inhibitor and overcomes the long-standing challenge confronted by orthosteric inhibitors of targeting JAK1 while sparing JAK2. As JAK3 only forms heterodimers with JAK1, we did not assess whether VVD-118313 independently inhibits JAK3 activity in cells, but we do not anticipate cross-reactivity with JAK3 because it lacks the corresponding liganded cysteine.

Our initial mechanistic studies indicate that VVD-118313 may inhibit JAK1 by blocking trans-phosphorylation of the activation loop of this kinase. This effect was much stronger for VVD-118313 compared to orthosteric JAK inhibitors, and we even observed some attenuation of JAK1 transphosphorylation for the C817A mutant. Our data thus point to a strong potential for allosteric regulation of JAK1 phosphorylation by the VVD-118313-binding pocket. Considering this pocket mirrors the myristate-binding pocket of ABL^45–47^, it is tempting to speculate that endogenous metabolites might also bind to JAK1 at this site to regulate its function. We also wonder how many additional kinases may possess this ligandable pocket and prove amenable to a similar mode of allosteric small-molecule regulation. In this regard, structure-based alignment of 497 human kinase domains reveals a diversity of cysteines at various positions proximal to the Bcr-Abl myristate pocket^68^.

The remarkable proteome-wide selectivity displayed by VVD-118313 for JAK_C817 across more than 14,000 quantified cysteines in human and mouse immune cell proteomes supports the broader utility of this compound as a cellular probe to investigate the specific biological functions of JAK1. Indeed, using VVD-118313, we discovered that JAK1 makes differential contributions to IL-6-STAT3 signaling in human PBMCs versus mouse splenocytes, a finding that may have been obscured in past experiments with other JAK1 inhibitors due to their lack of subtype selectivity. We also found that the TYK2 inhibitor BMS-986165 was noticeably more potent in blocking IL-6-STAT3 signaling in mouse splenocytes compared to human PBMCs (**Extended Data Fig. 6f** and **Fig. 5h**). Thus, by using a combination of allosteric inhibitors with high subtype selectivity, we have provided evidence for species and/or immune cell type differences in the relative contributions of JAK family members to an important cytokine signaling pathway. Curiously, splenocytes from TYK2^-/-^ mice have been reported to have unperturbed IL-6-STAT3 signaling^69,70^, which could indicate that, in this setting, JAK1 compensates for chronic TYK2 disruption. VVD-118313 should also help to illuminate JAK1 contributions to other signaling pathways, including, for instance, PI3K/AKT, and MAPK/ERK/p38 and p38 kinase signaling^71–73^.

Projecting forward, we believe that, while VVD-118313 was capable of inhibiting JAK1 in mice, the full utility of this chemical probe for *in vivo* studies would benefit from improvements in its pharmacokinetic properties. We also wonder if further exploration of the SAR might uncover compounds that show greater functional activity for TYK2, which could provide an additional class of useful chemical probes that act as dual allosteric JAK1/TYK2 inhibitors. From a translational perspective, it is enticing to consider the possibility that covalent allosteric JAK1 inhibitors may circumvent some of the systemic toxicities associated with pan-JAK inhibition in humans. Alternative ways to address this problem have been brought forward, including tissue-restricted JAK inhibitors, such as gut-restricted izencitinib (TD-1473) for ulcerative colitis^74^, and lung-restricted nezulcitinib (TD-0903) for COVID-19^75^; however, these compounds have not yet displayed efficacy in clinical studies in humans^76,77^, pointing to the potential need for systemic exposure. Of course, it is possible that selective allosteric JAK1 inhibitors may also sacrifice a proportion of the efficacy observed with systemic pan-JAK inhibitors, but we believe our cellular studies showing that VVD-118313 matches the activity of pan-JAK inhibitors in suppressing multiple human cytokine pathways (e.g., IFNα, IL-6) should encourage further pursuit of highly selective JAK1 inhibitors for immunological disorders.

Finally, we believe that our findings provide another compelling example of the utility of chemical proteomics for the discovery of small molecules that act by unconventional mechanisms^41,78–82^. Chemical proteomic platforms like ABPP have a distinct advantage of evaluating compounds against thousands of sites on endogenously expressed proteins and can thus uncover ligandable pockets that may be missed by more conventional assays performed with purified proteins or protein domains. Nonetheless, chemical proteomics is still principally a binding assay and interpreting how newly discovered small-molecule interactions affect the functions of proteins can be technically challenging. Here, we benefited from the availability of robust cell-based activity assays for JAK1 and, in particular, structural information that emphasized the potential functionality of a conserved pocket adjacent to the covalently liganded C817 residue^47^. As the structures of more full-length proteins are solved or accurately predicted^83,84^, the integration of this information with global small-molecule interaction maps furnished by chemical proteomics should facilitate the discovery of additional cryptic functional and druggable allosteric pockets on a broad range of proteins.

## Supporting information

Chemistry Methods

Supplementary Figures and Biology Methods

## Acknowledgements

This work was supported by the N.I.H. (R35 CA231991) and a Sir Henry Wellcome Postdoctoral Fellowship (210890/Z/18/Z) awarded to (M.E.K). We thank K. Yao, P. Gao, F. Zhang, X. Li, W. Wu, X. Jia and M. Xu for their contribution to the synthetic chemistry and B. Melillo for guidance with chemical analysis.

