## Supplementary material for "Selective inhibitors of JAK1 targeting a subtype-restricted allosteric cysteine": Chemistry Methods

#### Synthetic and analytical chemistry

##### General methods

All chemicals, including anhydrous solvents, were obtained from commercial suppliers (including Alfa, Adamas, Sigma-Aldrich, Pharmablock and Bidepharm) and used without further purification. Reactions were monitored by TLC and LCMS. $^{1}H$ and $^{13}C$ NMR spectra were obtained on Bruker 400 NMR spectrometer or Bruker AVIII HD 600 NMR spectrometer equipped with a 5 mm CPDCH (C-H) CryoProbes. All $^{1}H$ and $^{13}C$ NMR experiments are reported in δ units parts per million (ppm) and are listed relative to residual signals for $CHCl_3$ (7.26 ppm/ $^{1}H$, 77.16/$^{13}C$) or DMSO (2.50/$^{1}H$, 39.52/$^{13}C$). Two peaks were observed for all carbons, indicative of the presence of two amide rotational isomers. Mass measurements for high-resolution mass spec (HRMS) were performed on a Waters Xevo G2-XS TOF calibrated against sodium formate clusters and using a LeuEnk lockmass. Expected monoisotopic masses were calculated using MassLynx 4.1 and the m/z values for calibrant and lockmass were MassLynx-default values and were required to be within 5 ppm error. HPLC purifications were performed using a Phenomenex Luna C18 column using the following solvent systems unless otherwise noted: Neutral – solvent A: 10 mM $NH_4HCO_3$ in water; solvent B: acetonitrile; Acidic – solvent A: 0.1% (v/v) TFA in water; solvent B: 0.1% (v/v) TFA in acetonitrile. All other solvent systems are described as v/v, unless otherwise noted. SFC separations were performed as described, using the same conditions for preparative and analytical analysis. Analytic traces provided for compounds **2a**, **3a**, **5a**, **5b**, **5c**, **5d**. Specific optical rotations (0.01 g 100 $ml^{-1}$) were measured in a 100 mm quartz cell using a Anton Paar MCP 100 polarimeter at 20 °C.

Abbreviations: THF – tetrahydrofuran; KHMDS – potassium hexamethyldisilazide; $PPh_3$ – triphenylphosphine; TFA – trifluoroacetic acid; T3P - 2,4,6-tripropyl-1,3,5,2,4,6-trioxatriphosphorinane-2,4,6-trioxide; DIEA - *N*,*N*-diisopropylethylamine; BuLi – n-butyllithium; MsCl – methane sulfonyl chloride; TsOH – toluene sulfonic acid; DMP - Dess–Martin periodinane

Preparation of compound **1**

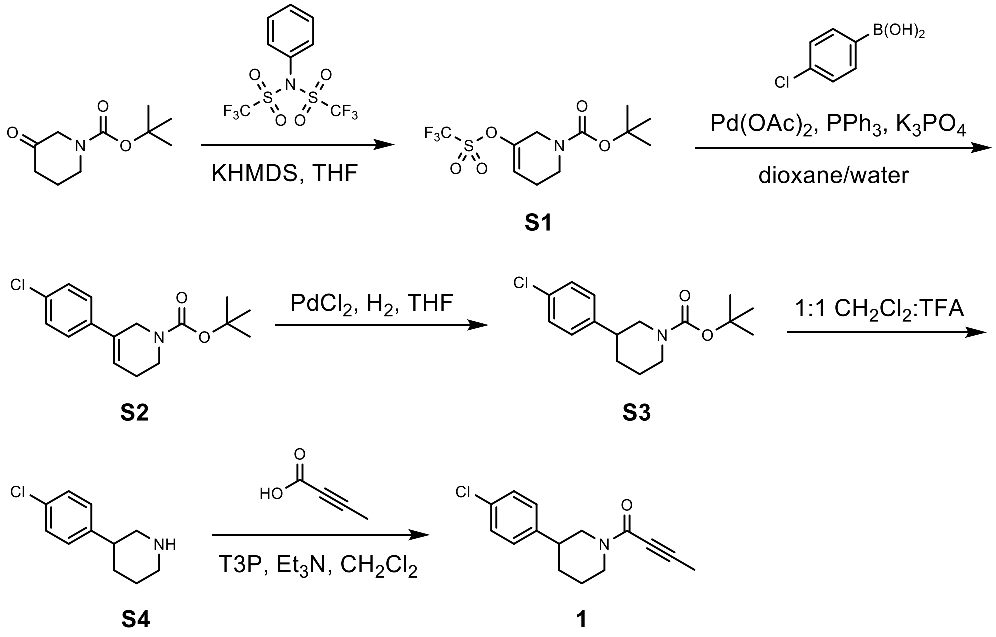


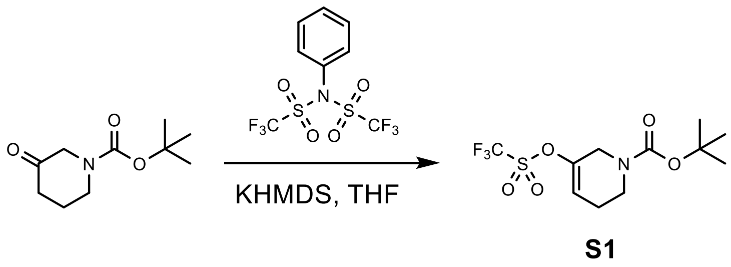


**tert-Butyl 5-(trifluoromethylsulfonyloxy)-3,6-dihydro-2H-pyridine-1-carboxylate (S1)**

To a solution prepared by diluting 1 M KHMDS in THF (63 mL, 63 mmol) with THF (100 mL) at -78 °C under $N_2$ (15 PSI) was added a solution of tert-butyl 3-oxopiperidine-1-carboxylate (10.0 g, 50.2 mmol) in THF (100 mL) dropwise and the resulting mixture was stirred at -78 °C for 1.5 hours. A solution of N-phenylbis (trifluoromethanesulfonimide) (18.83 g, 52.7 mmol) in THF (100 mL) was added to the mixture dropwise at -78 °C and the mixture was stirred at -78 °C for 1 hour. The reaction mixture was allowed to warm to room temperature. After 1 hour, the mixture was quenched with a 10% NaOH solution (300 mL) and extracted with ethyl acetate (2 x 200 mL). The combined organic phases were washed with brine (500 mL), dried over $Na_2SO_4$, filtered, concentrated and purified by flash chromatography (0-10% v/v ethyl acetate in petroleum ether) to give tert-butyl 5-(trifluoromethylsulfonyloxy)-3,6-dihydro-2H-pyridine-1-carboxylate (**S1**) (13.00 g, 392.38 mmol) as a light yellow oil. Spectral characterization matched those found in the literature[1].

**$^1$H NMR** (400 MHz, $CDCl_3$): δ= 5.93 (br s, 1 H), 4.05 (br d, *J*=1.76 Hz, 2 H), 3.53 (br s, 2 H), 2.31 (br d, *J*=4.11 Hz, 2 H), 1.48 (s, 9 H) ppm. 16/16 protons observed/expected.

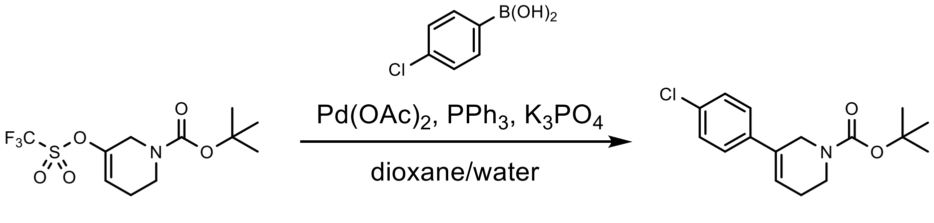


**S1** **S2**

**tert-Butyl 5-(4-chlorophenyl)-3,6-dihydro-2H-pyridine-1-carboxylate (S2)**

A solution of tert-butyl 5-(trifluoromethylsulfonyloxy)-3,6-dihydro-2H-pyridine-1-carboxylate (1.00 g, 3.018 mmol) (**S1**), 4-chlorophenylboronic acid (0.47 g, 3.02 mmol), $Pd(OAc)_2$ (76.8 mg, 0.302 mmol), $PPh_3$ (158.3 mg, 0.604 mmol) and $K_3PO_4$ (0.83 g, 2 eq., 6.0 mmol) in a 10:1 mixture of 1,4-dioxane/water (33 mL, 0.092 M) was stirred at 80 °C for 2 hours. TLC (5:1 petroleum ether:ethyl acetate; Rf: starting material 0.7, product 0.8) showed the reaction was complete. The mixture was concentrated and purified by flash chromatography (0-10% v/v ethyl acetate in petroleum ether) to give tert-butyl 5-(4-chlorophenyl)-3,6-dihydro-2H-pyridine-1-carboxylate (**S2**) (0.55 g, 1.872 mmol, 62.0% yield) as a yellow oil.

**$^1$H NMR** (400 MHz, $CDCl_3$): δ = 7.30 (s, 3 H), 7.24 - 7.27 (m, 1 H), 6.19 (dt, *J*=3.96, 2.23 Hz, 1 H), 4.24 (br s, 1 H), 3.52 - 3.58 (m, 2 H), 2.39 - 2.50 (m, 2 H), 1.88 - 1.99 (m, 1 H), 1.50 (s, 9 H) ppm. 20/20 protons observed/expected.

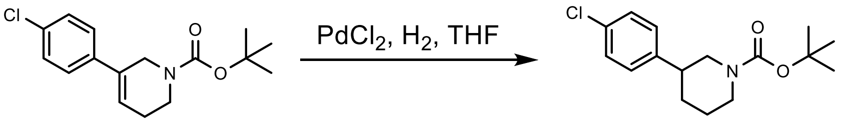


**S2** **S3**

**tert-Butyl 3-(4-chlorophenyl)piperidine-1-carboxylate (S3)**

A mixture of tert-butyl 5-(4-chlorophenyl)-3,6-dihydro-2H-pyridine-1-carboxylate (300.0 mg, 1.021 mmol) (**S2**) and $PdCl_2$ (20.0 mg) in THF (10 mL, 0.102 M) was stirred under $H_2$ (15 PSI) at 25 °C for 1 hour. TLC (5:1 petroleum ether:ethyl acetate; Rf: starting material 0.8, product 0.75) showed the reaction was complete. The reaction was concentrated and purified by acidic prep-HPLC (60-90% v/v solvent B) to give tert-butyl 3-(4-chlorophenyl)piperidine-1-carboxylate (**S3**) (130 mg, 0.434 mmol, 43% yield) as a white solid. Spectral characterization matched those found in the literature[2].

**$^1$H NMR** (400 MHz, $CDCl_3$): δ 7.28 - 7.34 (m, 2 H), 7.22 - 7.28 (m, 2 H), 4.09 (br d, *J*=13.30 Hz, 2 H), 2.83 (br s, 2 H), 2.65 (tt, *J*=11.34, 3.53 Hz, 1 H), 2.65 (tt, *J*=11.34, 3.53 Hz, 1 H), 1.98 (br d, *J*=12.92 Hz, 1 H), 1.54 - 1.82 (m, 3 H), 1.46 (s, 9 H) ppm. 23/22 protons observed/expected.

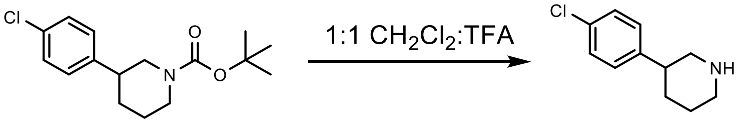


**S3** **S4**

##### 3-(4-Chlorophenyl)piperidine (S4)

A mixture of tert-butyl 3-(4-chlorophenyl) piperidine-1-carboxylate (130.0 mg, 0.44 mmol) (**S3**) in $CH_2Cl_2$ (2 mL) and TFA (2 mL) was stirred at 25 °C for 0.5 hours. TLC (1:1 petroleum ether:ethyl acetate; Rf: starting material 0.8, product 0.1) showed the reaction was complete. The reaction was concentrated directly to give 3-(4-chlorophenyl)piperidine (**S4**) (80.0 mg, 0.409 mmol, 93.0% yield) as a yellow oil.

**$^1$H NMR** (400 MHz, $CDCl_3$): δ 7.34 -7.39 (m, 2 H), 7.24 - 7.32 (m, 2 H), 3.36 - 3.48 (m, 2 H) 2.92 - 3.14 (m, 3 H), 1.98 - 2.13 (m, 2 H), 1.72 - 1.95 (m, 2 H), ppm. 13/14 protons observed/expected.

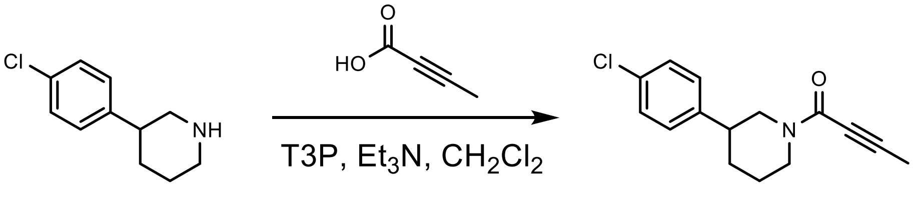


**S4** **1**

##### 1-[3-(4-Chlorophenyl)-1-piperidyl]but-2-yn-1-one (1)

To a solution of 3-(4-chlorophenyl)piperidine (80.0 mg, 0.409 mmol) (**S4**), $Et_3N$ (82.8 mg, 0.818 mmol), and but-2-ynoic acid (68.7 mg, 0.818 mmol) in $CH_2Cl_2$ (4 mL, 0.102 M) was added T3P (260.0 mg, 0.818 mmol) at 25 °C and the reaction was stirred at 25 °C for 0.5 hour. TLC (1:1 petroleum ether:ethyl acetate; Rf: starting material 0.1, product 0.4) showed the reaction was complete. The reaction was concentrated and purified by prep-TLC (1:1 petroleum ether:ethyl acetate) to give 1-[3-(4-chlorophenyl)-1-piperidyl]but-2-yn-1-one (80.7 mg, 0.307 mmol, 75% yield) 1-[3-(4-chlorophenyl)-1-piperidyl]but-2-yn-1-one (**1**) as a colorless oil.

**$^1$H NMR** (400 MHz, DMSO-$d_6$) δ = 7.42 – 7.36 (m, 2H), 7.36 – 7.28 (m, 2H), 4.41 – 4.15 (m, 2H), 3.31 – 3.09 (m, 1H), 2.82 – 2.55 (m, 2H), 2.00 (d, J = 22.5 Hz, 3H), 1.90 (dd, J = 12.6, 4.7 Hz, 1H), 1.74 (ddddd, J = 34.7, 18.7, 12.5, 6.3, 3.4 Hz, 2H), 1.45 (dtt, J = 37.7, 12.6, 4.0 Hz, 1H). 16/16 protons observed/expected.

**$^{13}$C NMR** (101 MHz, DMSO- $d_6$) δ = 151.88, 151.85, 142.10, 142.00, 131.26, 131.15, 128.99, 128.94, 128.49, 128.42, 89.48, 89.30, 73.02, 72.94, 52.21, 46.52, 46.44, 42.20, 41.08, 40.72, 31.47, 31.23, 25.63, 24.60, 3.34, 3.26. Apparent mixture of two amide rotational isomers.

**HRMS** (*m/z*): $[M+H]^+$ calculated for C15H16ClNO, 262.0993; found, 262.1009.

Preparation of compound **1a**

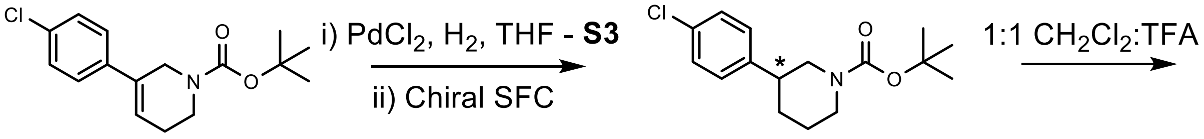


**S2** **S3a**

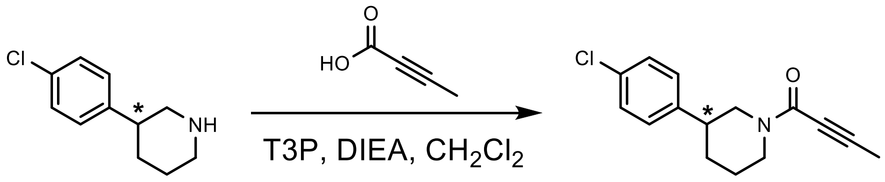


**S4a** **1a**

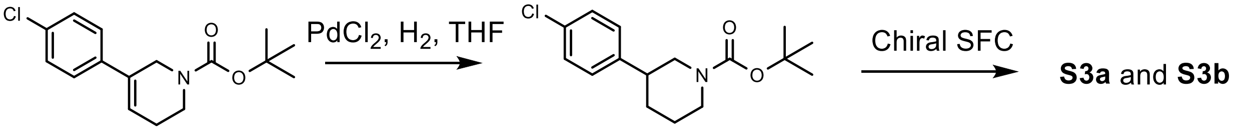


**S2** **S3** (racemate)

**tert-Butyl (3S)-3-(4-chlorophenyl) piperidine-1-carboxylate (S3a)**

A mixture of tert-butyl 5-(4-chlorophenyl)-3,6-dihydro-2H-pyridine-1-carboxylate (1.50 g, 5.106 mmol) (**S2**) and $PdCl_2$ (30.0 mg) in THF (25 mL, 0.117 M) was stirred under $H_2$ (15 Psi) at 25 °C for 1 hour. TLC (10:1 petroleum ether:ethyl acetate; Rf: starting material 0.6, product 0.5) showed the reaction was complete. The reaction was concentrated to give crude tert-butyl 3-(4-chlorophenyl)piperidine-1-carboxylate (1.50 g, 5.071 mmol, 99% yield) as a yellow oil. The crude product (300 mg) was purified by prep-TLC (10:1 petroleum ether:ethyl acetate; Rf: product 0.4) to give 100 mg of pure product (**S3**) which was then separated by chiral SFC (ChiralPak AD-3 liquid phase:[A-CO2; B-MeOH(0.05%IPAm, v/v)] B%: 10%-40%)) indicated a single enantiomer to enantiomers **S3a** (50.0 mg, 0.169 mmol) and **S3b** (50.0 mg, 0.169 mmol). The NMR spectra of **S3a** and **S3b** were identical.

**$^1$H NMR** (400 MHz, $CDCl_3$): δ= 7.32 - 7.35 (m, 2 H), 7.17 (s, 2 H), 4.13 - 4.17 (m, 2 H), 2.59 - 2.82 (m, 3 H), 2.00 (br d, *J*=8.53 Hz, 1 H), 1.70 - 1.80 (m, 1 H), 1.57 - 1.67 (m, 2 H), 1.47 (s, 9 H) ppm. 22/22 protons observed/expected.

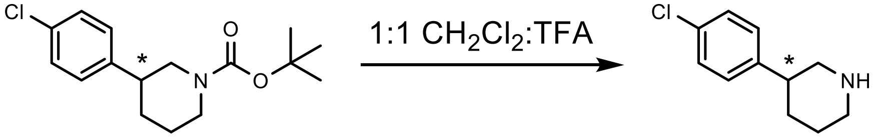


**S3a** **S4a**

##### 3-(4-Chlorophenyl)piperidine (S4a)

A mixture of tert-butyl 3-(4-chlorophenyl) piperidine-1-carboxylate (**S3a**) (50.0 mg, 0.169 mmol) in $CH_2Cl_2$ (2 mL) and TFA (2 mL) was stirred at 25 °C for 0.5 hours. TLC (10:1 petroleum ether:ethyl acetate; Rf starting material 0.30, product 0.00) showed the reaction was complete. The reaction was concentrated directly to give (3S)-3-(4-chlorophenyl)piperidine (**S4a**) (32.0 mg, 0.164 mmol, 97% yield) as a yellow oil.

**$^1$H NMR** (400 MHz, $CDCl_3$): δ = 1.76 - 1.93 (m, 2 H), 2.07 (br s, 2 H), 2.98 - 3.15 (m, 3 H) 3.38 - 3.52 (m, 2 H), 7.29 - 7.34 (m, 2 H), 7.36 - 7.42 (m, 2 H) ppm. 13/14 protons observed/expected.

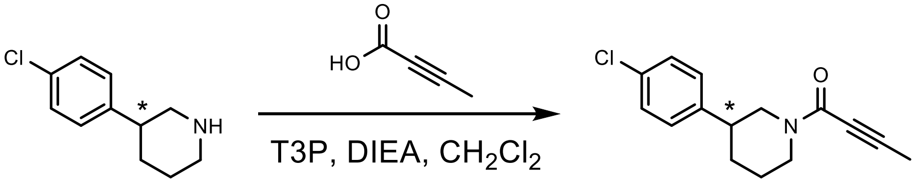


**S4a** **1a**

##### 1-(3-(4-Chlorophenyl)-1-piperidyl)but-2-yn-1-one (1a)

To a solution of 3-(4-chlorophenyl)piperidine (**S4a**) (33.0 mg, 0.169 mmol), and 2-butynoic acid (28.4 mg, 0.337 mmol) in $CH_2Cl_2$ (3 mL) was added T3P (165.0 mg, 0.337 mmol) and DIEA (45.0 mg, 0.337 mmol) at 25 °C and the reaction was stirred at 25 °C for 0.5 hour. TLC (1:1 petroleum ether:ethyl acetate; Rf starting material 0.10, product 0.60) showed the reaction was complete. The reaction was concentrated and purified by Prep-TLC (1:1 petroleum ether:ethyl acetate; Rf starting material 0.10, product 0.60) to give 1-[(3R)-3-(4-chlorophenyl)-1-piperidyl]but-2-yn-1-one (20.7 mg, 0.079 mmol, 47% yield) 1-(3-(4-Chlorophenyl)-1-piperidyl)but-2-yn-1-one (**1a**) as a white solid. Spectral characterization matched compound **1**.

**$^1$H NMR** (400 MHz, DMSO-$d_6$) δ = 7.47 – 7.27 (m, 4H), 4.28 (td, J = 24.9, 13.9 Hz, 2H), 3.40 – 3.31 (m, 1H), 3.19 (dt, J = 30.7, 12.5 Hz, 1H), 2.91 – 2.42 (m, 2H), 2.02 (t, J = 15.8 Hz, 3H), 1.89 (d, J = 12.4 Hz, 1H), 1.84 – 1.65 (m, 1H), 1.55 – 1.35 (m, 1H).

$[\alpha]_D^{20}$ = -130 (c 0.01, $CHCl_3$)

Preparation of compound **2**

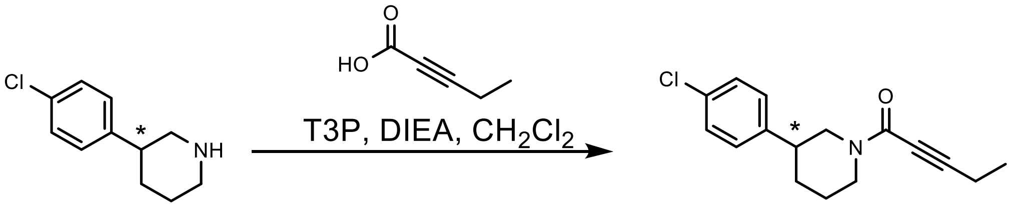

**S4a** **2a**

**1-(3-(4-Chlorophenyl)-1-piperidyl)pent-2-yn-1-one (2a)**

To a solution of 3-(4-chlorophenyl)piperidine (**S4a**) (50.0 mg, 0.256 mmol) and pent-2-ynoic acid (50.1 mg, 0.511 mmol) in $CH_2Cl_2$ (3 mL, 0.085 M) was added T3P (290.0 mg, 0.456 mmol) and DIEA (80.0 mg, 0.600 mmol) at 25 °C and the reaction was stirred at 25 °C for 0.5 hour. TLC (1:1 petroleum ether:ethyl acetate; Rf: starting material 0.1, product 0.6) showed the reaction was complete. The reaction was concentrated and purified by prep-TLC (1:1 petroleum ether: ethyl acetate) and prep-HPLC (neutral conditions, 55-75% B) to give 1-(3-(4-chlorophenyl)-1-piperidyl)pent-2-yn-1-one (**2a**) (32.5 mg, 0.118 mmol, 46.1% yield) as a yellow oil which was a single enantiomer by analytical chiral SFC (ChiralPak AS liquid phase:[A-CO2; B-Isopropanol(0.05%IPAm, v/v)] B%: 10%-25%)) (ee 97.5%).

**$^1$H NMR** (400 MHz, DMSO-*d*$_6$) δ 7.37 (qd, J = 8.6, 2.7 Hz, 2H), 7.34 – 7.24 (m, 2H), 4.37 – 4.18 (m, 2H), 3.26 (td, J = 11.6, 8.0 Hz, 1H), 3.21 – 3.11 (m, 1H), 2.84 – 2.54 (m, 1H), 2.49 – 2.27 (m, 2H), 1.95 – 1.58 (m, 3H), 1.53 – 1.35 (m, 1H), 1.10 (dtd, J = 25.7, 7.6, 2.9 Hz, 3H). 18/18 protons observed/expected.

**$^{13}$C NMR** (101 MHz, DMSO-*d*$_6$) δ 151.92, 142.13, 142.00, 131.30, 131.18, 128.97, 128.54, 128.45, 94.29, 94.18, 73.17, 73.13, 52.50, 46.61, 46.49, 42.26, 41.11, 40.82, 40.15, 39.94, 39.73, 39.52, 39.31, 39.11, 38.90, 31.26, 25.63, 24.64, 12.86, 11.78, 11.69. Apparent mixture of two amide rotational isomers.

**HRMS** (*m/z*): $[M+H]^+$ calculated for $C_{16}H_{18}ClNO$, 276.1150; found, 276.1162.

$[\alpha]_D^{20}$ = -30 (c 0.01, $CHCl_3$)

Preparation of compound **3**

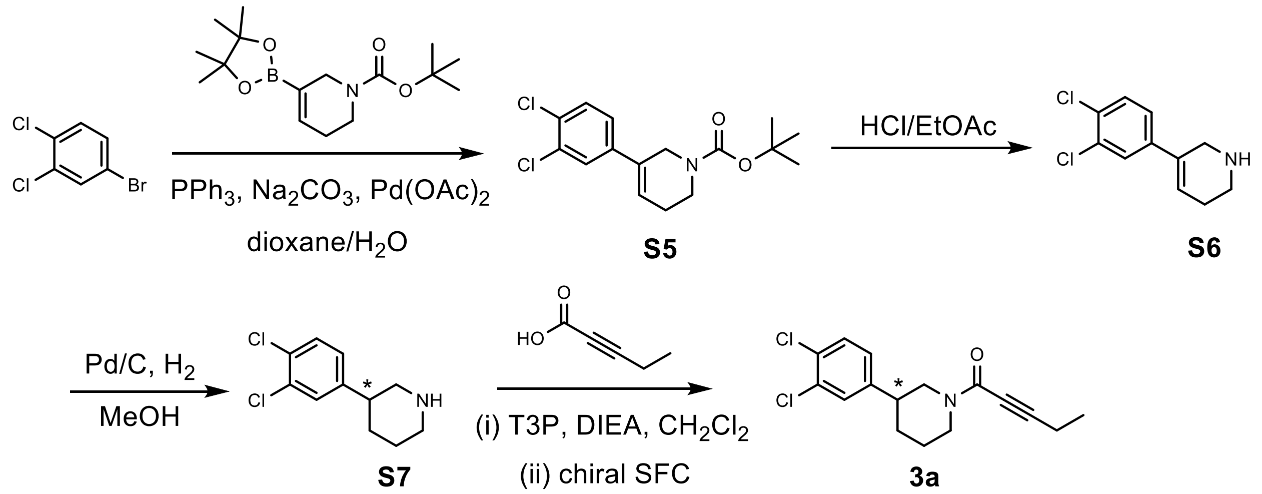


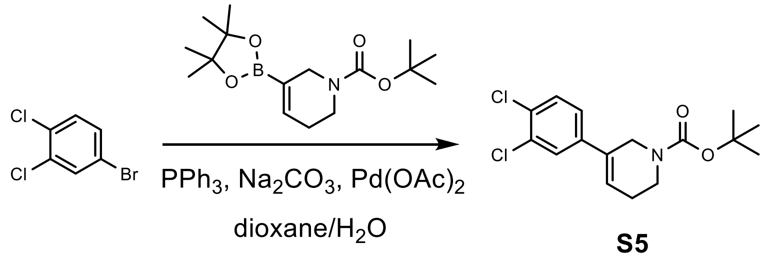


**tert-Butyl 5-(3,4-dichlorophenyl)-3,6-dihydro-2H-pyridine-1-carboxylate (S5)**

To a solution of 1-bromo-3,4-dichlorobenzene (1.00 g, 4.427 mmol, 1 eq.), $Na_2CO_3$ (938.4 mg, 8.854 mmol, 2 eq.) and tert-butyl 5-(4,4,5,5-tetramethyl-1,3,2-dioxaborolan-2-yl)-3,6-dihydro-2H-pyridine-1-carboxylate (1.37 g, 4.427 mmol, 1 eq.) in dioxane/$H_2O$ (5:1, 12 mL) was added $PPh_3$ (11.6 mg, 0.044 mmol, 0.01 eq.) and $Pd(OAc)_2$ (0.003 eq., 3.3 mg) at 20 °C and the mixture was stirred at 100 °C for 16 h under $N_2$. TLC (10:1 petroleum ether:ethyl acetate; Rf starting material 0.8, product 0.5) showed the starting material was consumed completely. The reaction was poured into water (15 mL) and extracted with ethyl acetate (3 x 5 mL). The organic layers were washed with brine (5 mL), dried over $Na_2SO_4$, filtered and concentrated under reduced pressure to give a residue. The residue was purified by flash chromatography (5-10% v/v ethyl acetate in petroleum ether) to give tert-butyl 5-(3,4-dichlorophenyl)-3,6-dihydro-2H-pyridine-1-carboxylate (**S5**) (1.20 g, 3.656 mmol, 82.6% yield) as a colorless oil.

**$^1$H NMR** (400 MHz, $CDCl_3$): δ 7.45 (br s, 1 H), 7.40 (d, *J*=8.8 Hz, 1 H), 7.19 (dd, *J*=1.8, 8.3 Hz, 1 H), 6.22 (tt, *J*=1.9, 4.1 Hz, 1 H), 4.21 (br s, 2 H), 3.54 (t, *J*=5.7 Hz, 2 H), 2.32 (br d, *J*=3.5 Hz, 2 H), 1.50 (s, 9 H) ppm. 19/19 protons observed/expected.

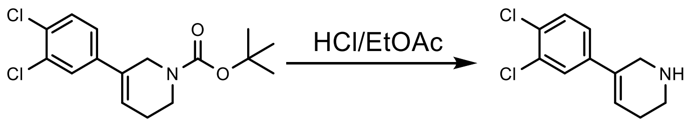


**S5** **S6**

**5-(3,4-Dichlorophenyl)-1,2,3,6-tetrahydropyridine (S6)**

A solution of tert-butyl 5-(3,4-dichlorophenyl)-3,6-dihydro-2H-pyridine-1-carboxylate (**S5**) (300.0 mg, 0.914 mmol) in HCl/ethyl acetate (2M, 3 mL) was stirred at 20 °C for 0.5 hr. TLC (3:1 petroleum ether:ethyl acetate; Rf: starting material 0.45, product 0.0) showed the starting material was consumed completely. The reaction was concentrated under reduced pressure to give crude 5-(3,4-dichlorophenyl)-1,2,3,6-tetrahydropyridine (**S6**) (200.0 mg, 0.877 mmol, 96% yield) as a white solid.

**$^1$H NMR** (400 MHz, $CD_3OD$): δ 7.61 (d, *J*=2.2 Hz, 1 H), 7.54 (d, *J*=8.3 Hz, 1 H), 7.37 (dd, *J*=2.4, 8.6 Hz, 1 H), 6.44 (td, *J*=2.1, 4.1 Hz, 1 H), 4.06 (q, *J*=2.2 Hz, 2 H), 3.37 (t, *J*=6.1 Hz, 2 H), 2.55 - 2.66 (m, 2 H) ppm.10/11 protons observed/expected.

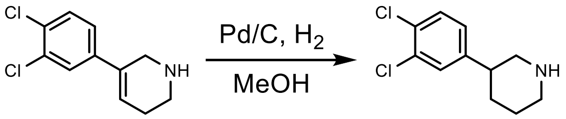


**S6** **S7**

**3-(3,4-Dichlorophenyl)piperidine (S7)**

To a solution of crude 5-(3,4-dichlorophenyl)-1,2,3,6-tetrahydropyridine (**S6**) (200.0 mg, 0.877 mmol) in MeOH (5 mL) was added Pd/C (5%, 10.0 mg) at 20 °C and the mixture was stirred under 15 PSI $H_2$ at the same temperature for 1 hr. The reaction was filtered over celite and concentrated under reduced pressure to give 3-(3,4-dichlorophenyl)piperidine (**S7**) (200.0 mg, 0.869 mmol, 99% yield) as a yellow solid.

**$^1$H NMR** (400 MHz, $CD_3OD$): δ 7.45 - 7.49 (m, 2 H), 7.23 (dd, *J*=2.2, 8.3 Hz, 1 H), 3.33 - 3.47 (m, 2 H), 2.90 - 3.14 (m, 3 H), 1.94 - 2.08 (m, 2 H), 1.68 - 1.93 (m, 2 H) ppm. 12/13 protons observed/expected.

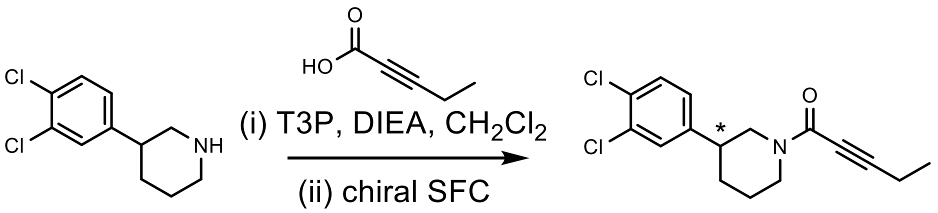


**S7** **3a**

**1-(3-(3,4-Dichlorophenyl)-1-piperidyl)pent-2-yn-1-one (3a)**

To a solution of 3-(3,4-dichlorophenyl)piperidine (**S7**) (200.0 mg, 0.869 mmol), DIEA (337.0 mg, 2.607 mmol) and T3P (1.38 g, 2.173 mmol) in $CH_2Cl_2$ (3 mL) was added pent-2-ynoic acid (93.8 mg, 0.956 mmol) at 0 °C and the mixture was stirred at 20 °C for 0.5 h. The reaction was poured into $H_2O$ (8 mL) and extracted with $CH_2Cl_2$ (3 x 2 mL). The organic layers were washed with brine (2 mL), dried over $Na_2SO_4$, filtered and concentrated under reduced pressure to give a residue. The residue was purified by chiral SFC (OJ column, 10 to 40% isopropanol) to give 1-(3-(3,4-dichlorophenyl)-1-piperidyl)pent-2-yn-1-one (**3a**) (27.7 mg, 0.089 mmol, 10.2% yield) as a white solid and single enantiomer (ee 99%).

**$^1$H NMR** (400 MHz, DMSO-$d_6$) δ 7.65 – 7.53 (m, 2H), 7.31 (ddd, J = 10.4, 8.3, 2.1 Hz, 1H), 4.35 – 4.16 (m, 2H), 3.17 (td, J = 12.7, 2.8 Hz, 1H), 2.84 (dd, J = 12.7, 11.4 Hz, 1H), 2.80 – 2.58 (m, 1H), 2.38 (dq, J = 21.1, 7.5 Hz, 2H), 1.94 – 1.85 (m, 1H), 1.84 – 1.65 (m, 2H), 1.56 – 1.33 (m, 1H), 1.10 (dt, J = 23.4, 7.4 Hz, 3H). 17/17 protons observed/expected.

**$^{13}$C NMR** (101 MHz, DMSO-$d_6$) δ = 151.89, 151.87, 144.33, 144.21, 131.16, 131.09, 130.68, 130.59, 129.27, 129.23, 129.21, 129.15, 127.63, 126.98, 94.28, 94.17, 73.10, 73.06, 52.00, 46.50, 46.03, 41.92, 40.82, 40.71, 40.15, 39.94, 39.73, 39.52, 39.31, 39.10, 38.89, 31.07, 25.49, 24.49, 12.82, 12.79, 11.74, 11.68. Apparent mixture of two amide rotational isomers.

**HRMS** (*m/z*): $[M+H]^+$ calculated for $C_{16}H_{17}Cl_2NO$, 310.0760; found, 310.0777.

$[\alpha]_D^{20}$ = -170 (c 0.01, $CDCl_3$)

<u>Preparation of compound **4**</u>

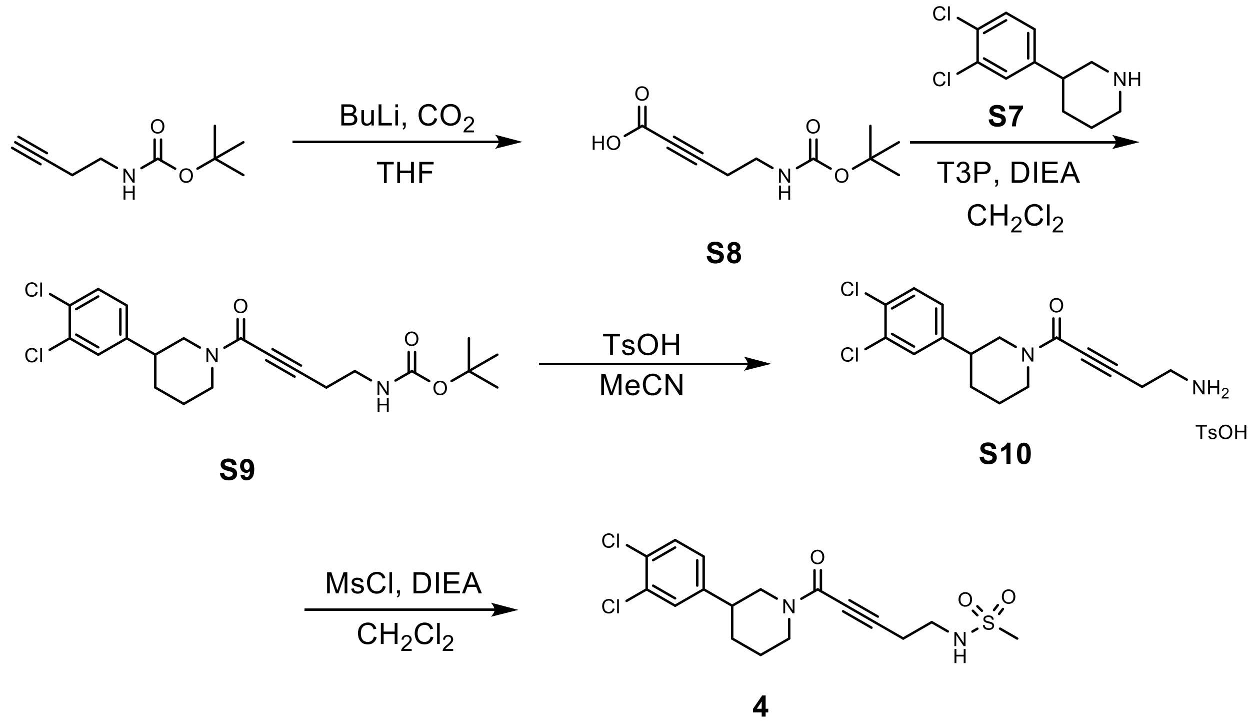

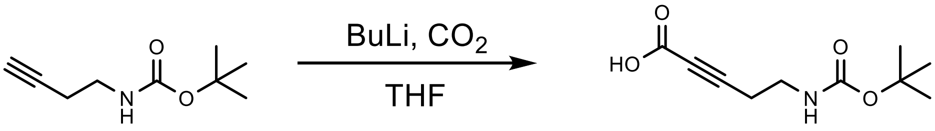


**S8**

**5-(tert-Butoxycarbonylamino)pent-2-ynoic acid (S8)**

To a solution of tert-butyl N-but-3-ynylcarbamate (200.0 mg, 1.182 mmol, 1 eq.) in THF (2 mL) was added dropwise n-BuLi (2.2 eq., 1.05 mL, 2.5 M) at -78 °C. The mixture was stirred at -78 °C for 1 hour. To the mixture was then added dry ice (5 eq., 1.5 g) at -78 °C. The mixture was stirred at -78 °C for 0.5 hours. The temperature of the reaction mixture was gradually raised to room temperature. The reaction mixture was quenched with water (10 mL), and then 2M hydrochloric acid was added to adjust to pH = 3 and the mixture was extracted with ethyl acetate (3 x 3 mL). The combined organic layers were dried over $Na_2SO_4$, filtered and concentrated under reduced pressure to give 5-(tert-butoxycarbonylamino)pent-2-ynoic acid (**S8**) (250 mg, 1.172 mmol, 99% yield) as yellow oil. Spectral characterization matched that found in the literature[3].

**$^1$H NMR** (400 MHz, $CDCl_3$): δ = 3.24-3.41 (m, 2 H), 2.56 (br d, *J*=5.5 Hz, 2 H), 1.41-1.49 (m, 9 H) ppm. 13/14 protons observed/expected.

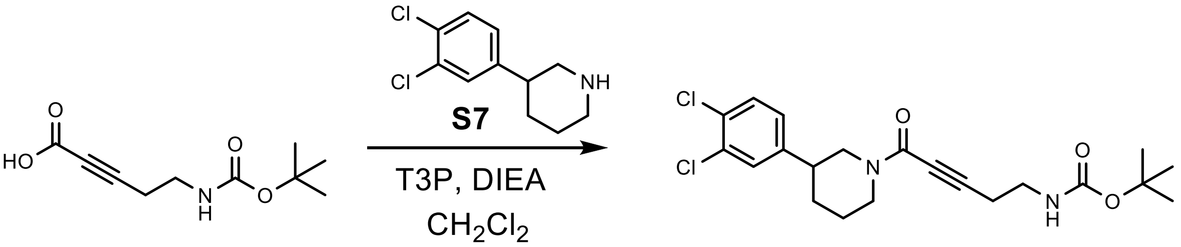


**S8** **S9**

**tert-Butyl N-[5-[3-(3,4-dichlorophenyl)-1-piperidyl]-5-oxo-pent-3-ynyl]carbamate (S9)**

To a solution of 3-(3,4-dichlorophenyl)piperidine (**S7**) (269.8 mg, 1.172 mmol), DIEA (454.6 mg, 3.517 mmol) and T3P (1.87 g, 2.931 mmol) in $CH_2Cl_2$ (4 mL) was added 5-(tert-butoxycarbonylamino)pent-2-ynoic acid (**S8**) (250 mg, 1.172 mmol) at 0 °C and the mixture was warmed to 20 °C and stirred for 0.5 h. TLC (1:1 petroleum ether:ethyl acetate; Rf starting material 0.0, product 0.7) showed the starting material was consumed completely. The reaction was poured into $H_2O$ (30 mL) and extracted with $CH_2Cl_2$ (3 x 8 mL). The organic layers were washed with brine (10 mL), dried over $Na_2SO_4$, filtered and concentrated under reduced pressure to give a residue. The residue was purified by flash chromatography (10-40% v/v ethyl acetate in petroleum ether) to give tert-butyl N-[5-[3-(3,4-dichlorophenyl)-1-piperidyl]-5-oxo-pent-3-ynyl]carbamate (**S9**) (240 mg, 0.564 mmol, 48% yield) as a yellow oil.

**$^1$H NMR** (400 MHz, $CDCl_3$): δ = 7.30-7.40 (m, 2 H), 7.08 (td, *J*=8.6, 2.1 Hz, 1 H), 4.91 (br s, 1 H), 4.56-4.69 (m, 1 H), 4.35-4.49 (m, 1 H), 3.30-3.41 (m, 2 H), 3.04-3.14 (m, 1 H), 2.49-2.66 (m, 4 H), 1.62-1.78 (m, 3 H), 1.45 (s, 9 H) ppm. 25/26 protons observed/expected.

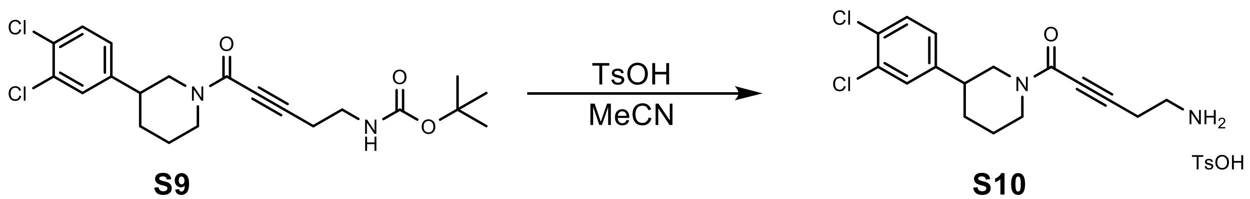


**S9** **S10**

**5-Amino-1-[3-(3,4-dichlorophenyl)-1-piperidyl]pent-2-yn-1-one (S10)**

To a solution of tert-butyl N-[5-[3-(3,4-dichlorophenyl)-1-piperidyl]-5-oxo-pent-3-ynyl]carbamate (**S9**) (200 mg, 0.470 mmol, 1 eq.) in MeCN (2 mL) was added TsOH (81.0 mg, 0.470 mmol, 1 eq.) at 20 °C and the mixture was stirred at 65 °C for 1 hr. TLC (1:1 petroleum ether:ethyl acetate; Rf starting material 0.50, product 0.00) showed the starting material was consumed completely. The reaction was concentrated under reduced pressure to give 5-amino-1-[3-(3,4-dichlorophenyl)-1-piperidyl]pent-2-yn-1-one tosylate salt (**S10**) (150 mg) as white solid which was carried on crude without further purification.

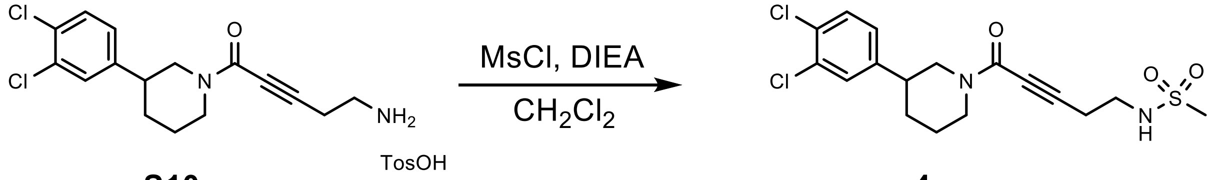


**S10** **4**

**N-[5-[3-(3,4-Dichlorophenyl)-1-piperidyl]-5-oxo-pent-3-ynyl]methanesulfonamide (4)**

To a solution of 5-amino-1-[3-(3,4-dichlorophenyl)-1-piperidyl]pent-2-yn-1-one (**S10**) (90 mg, 0.277 mmol) in $CH_2Cl_2$ (1 mL) was added DIEA (35.8 mg, 0.277 mmol) and MsCl (31.7 mg, 0.277 mmol) at 0 °C and the reaction was stirred at 0 °C for 0.5 hr. The reaction was poured into water (7 mL) and extracted with $CH_2Cl_2$ (3 x 2 mL. The combined organic layers were washed with brine (2 mL), dried over $Na_2SO_4$, filtered and concentrated under reduced pressure to give a residue. The residue was purified by acidic prep-HPLC (column: Nano-micro Kromasil C18 100*30mm 8um) to give N-[5-[3-(3,4-dichlorophenyl)-1-piperidyl]-5-oxo-pent-3-ynyl]methanesulfonamide (56.0 mg, 0.139 mmol, 50% yield over two steps) (**4**) as a yellow oil.

**$^1$H NMR** (600 MHz, CDCl3) δ 7.39 (dd, J = 25.1, 8.3 Hz, 1H), 7.31 (dd, J = 13.1, 2.1 Hz, 1H), 7.07 (ddd, J = 17.3, 8.3, 2.1 Hz, 1H), 5.61 (t, J = 6.4 Hz, 1H), 4.63 – 4.54 (m, 1H), 4.41 – 4.34 (m, 1H), 3.39 – 3.27 (m, 2H), 3.13 – 3.04 (m, 1H), 2.97 (d, J = 19.7 Hz, 3H), 2.74 – 2.55 (m, 4H), 1.95 – 1.81 (m, 1H), 1.72 – 1.50 (m, 1H). 18/20 protons observed/predicted.

**$^{13}$C NMR** (151 MHz, $CDCl_3$) δ 152.87, 142.78, 132.85, 132.71, 131.16, 130.96, 130.87, 130.66, 129.17, 126.73, 126.71, 90.44, 90.12, 77.37, 77.16, 76.95, 75.43, 75.31, 53.39, 47.65, 47.55, 42.75, 41.81, 41.44, 41.34, 41.29, 40.95, 40.92, 31.96, 31.56, 25.99, 24.87, 21.48. Apparent mixture of two amide rotational isomers.

**HRMS** (*m/z*): $[M+H]^+$ calculated for $C_{17}H_{20}Cl_2N_2O_3S$, 403.0644; found, 403.0641.

Preparation of compound **5**

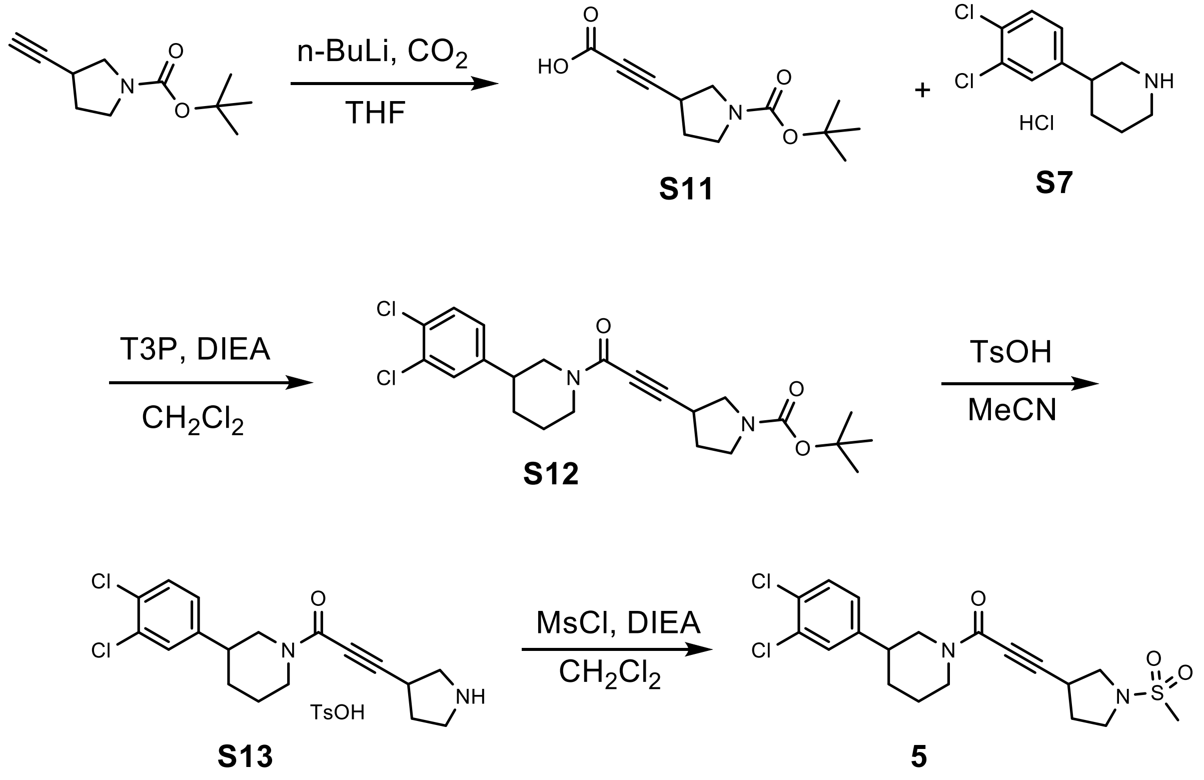


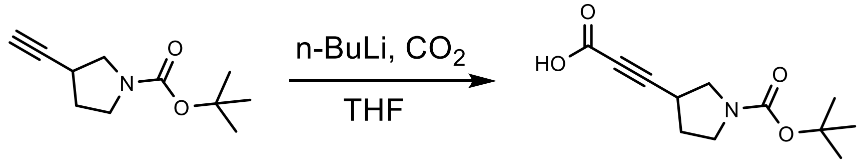


**S11**

**3-(1-tert-Butoxycarbonylpyrrolidin-3-yl)prop-2-ynoic acid (S11)**

n-BuLi (25.6 mL, 64.02 mmol, 1.25 eq.) was added to a solution of tert-butyl 3-ethynylpyrrolidine-1-carboxylate (10.0 g, 51.21 mmol, 1.00 eq.) in THF (100 mL, 0.51 M) dropwise at -65 °C. The mixture was stirred at -65 °C for 1 hour. Then dry ice (50 g) was added to the mixture at -65 °C. The resulting mixture was stirred at -65 °C for 30 min. TLC (5:1 petroleum ether: ethyl acetate; Rf starting material 0.45, product 0.05) showed the reaction was complete. The reaction mixture was quenched with water (100 mL), acidified by 1N HCl aqueous until pH = 3~4 and extracted with EtOAc (3 x 50 mL). The combined organic phase was washed with brine (30 mL), dried over $Na_2SO_4$, filtered and concentrated to give crude 3-(1-tert-butoxycarbonylpyrrolidin-3-yl)prop-2-ynoic acid (**S11**) (12.00 g, 50.20 mmol, 98.0% yield) as a yellow oil. The crude product was used to the next step without further purification.

$^{1}$**H NMR** (400 MHz, $CDCl_3$): δ = 3.33 - 3.73 (m, 4 H), 3.10 (br t, *J*=6.9 Hz, 1 H), 2.12 - 2.26 (m, 1 H), 1.88 (td, *J*=3.2, 6.7 Hz, 1 H), 1.46 (s, 9 H) ppm. 16/17 protons observed/expected.

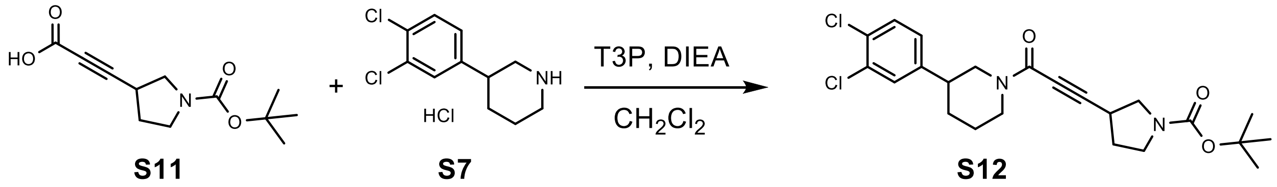


**tert-Butyl 3-[3-[3-(3,4-dichlorophenyl)-1-piperidyl]-3-oxo-prop-1-ynyl]pyrrolidine-1-carboxylate (S12)**

T3P (23.94 g, 37.61 mmol, 1.00 eq.) was added to the mixture of 3-(1-tert-butoxycarbonylpyrrolidin-3-yl)prop-2-ynoic acid (**S11**) (9.00 g, 37.61 mmol, 1.00 eq.), 3-(3,4-dichlorophenyl)piperidine hydrochloride (**S7**) (10.03 g, 37.61 mmol, 1.00 eq.) and DIEA (14.58 g, 112.84 mmol, 3.00 eq.) in $CH_2Cl_2$ (100 mL, 0.38 M). The resulted mixture was stirred at 20 °C for 1 hour. The reaction was poured into water (250 mL) and extracted with $CH_2Cl_2$ (3 x 100 mL). The combined organic layers were washed with brine (50 mL), dried over $Na_2SO_4$, filtered and concentrated under reduced pressure to give a residue. The residue was purified by column chromatography (eluent 0~30% ethyl acetate/petroleum ether gradient). tert-Butyl 3-[3-[3-(3,4-dichlorophenyl)-1-piperidyl]-3-oxo-prop-1-ynyl]pyrrolidine-1-carboxylate (**S12**) (13.00 g, 28.80 mmol, 77.0% yield) was obtained as a yellow oil.

$^{1}$**H NMR** (400 MHz, $CDCl_3$): δ = 7.37 - 7.47 (m, 1 H), 7.31 - 7.35 (m, 1 H), 7.07 (dd, *J*=1.8, 8.2 Hz, 1 H), 4.56 - 4.72 (m, 1 H), 4.31 - 4.40 (m, 1 H), 3.25 - 3.74 (m, 4 H), 3.02 - 3.19 (m, 2 H), 2.59 - 2.76 (m, 2 H), 2.15 - 2.29 (m, 1 H), 1.97- 2.12 (m, 2 H), 1.61 - 1.93 (m, 3 H), 1.46 (d, *J*=12.9 Hz, 9 H) ppm. 28/28 protons observed/expected.

$^{13}$**C NMR** (101 MHz, DMSO-$d_6$) δ 153.32, 151.51, 144.28, 144.14, 131.20, 131.09, 130.67, 130.60, 129.21, 129.17, 127.64, 92.40, 78.62, 50.74, 46.56, 46.08, 44.62, 40.82, 40.15, 39.94, 39.73, 39.52, 39.31, 39.11, 38.90, 31.48, 31.03, 30.89, 30.65, 28.11, 28.02, 25.49, 24.47. Apparent mixture of two amide rotational isomers.

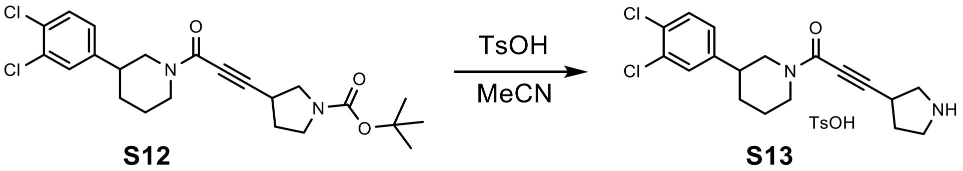


**1-[3-(3,4-Dichlorophenyl)-1-piperidyl]-3-pyrrolidin-3-yl-prop-2-yn-1-one tosylate (S13)**

TsOH (2.67 g, 15.51 mmol, 1.00 eq.) was added to a solution of tert-butyl 3-[3-[3-(3,4-dichlorophenyl)-1-piperidyl]-3-oxo-prop-1-ynyl]pyrrolidine-1-carboxylate (**S12**) (7.00 g, 15.51 mmol, 1.00 eq.) in MeCN (70 mL, 0.22 M). The resulting mixture was stirred at 65 °C for 6 hours. The reaction mixture was concentrated under reduced pressure to give crude 1-[3-(3,4-

dichlorophenyl)-1-piperidyl]-3-pyrrolidin-3-yl-prop-2-yn-1-one 4-methylbenzenesulfonic acid salt (**S13**) (8.00 g) as a yellow oil which was carried on crude without further purification.

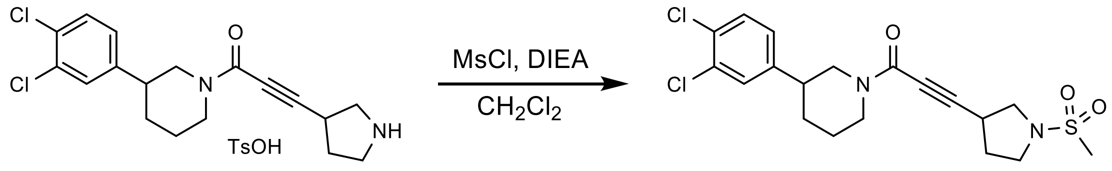


**S13** **5**

**1-(3-(3,4-Dichlorophenyl)piperidin-1-yl)-3-(1-(methylsulfonyl)pyrrolidin-3-yl)prop-2-yn-1-one (5)**

MsCl (1.84 g, 16.05 mmol, 1.20 eq.) was added to the mixture of 1-[3-(3,4-dichlorophenyl)-1-piperidyl]-3-pyrrolidin-3-yl-prop-2-yn-1-one;4-methylbenzenesulfonic acid (**S13**) (7.00 g, 13.37 mmol, 1.00 eq.) and DIEA (5.18 g, 40.12 mmol, 3.00 eq.) in $CH_2Cl_2$ (70 mL, 0.19 M) at 0 °C. The resulting mixture was stirred at 0 °C for 30 min. The reaction mixture was poured into water (200 mL) and extracted with $CH_2Cl_2$ (3 x 50 mL). The combined organic layers were washed with brine (50 mL), dried over $Na_2SO_4$, filtered and concentrated under reduced pressure to give a residue. The residue was purified by column chromatography (0-50% v/v ethyl acetate in petroleum ether) to provide crude product. The crude product was purified by prep-HPLC (column: Agela DuraShell C18 250*70 mm*10 um; liquid phase: [A-TFA/$H_2O$=0.075% v/v; B-ACN] B%: 35%-65%, 22 min]) to yield 1-(3-(3,4-dichlorophenyl)piperidin-1-yl)-3-(1-(methylsulfonyl)pyrrolidin-3-yl)prop-2-yn-1-one (**5**) (4.0 g, 8.46 mmol, 63% yield) as a white solid.

**$^1$H NMR** (400 MHz, DMSO-*d*$_6$) δ 7.63 – 7.55 (m, 2H), 7.31 (ddd, J = 10.5, 8.3, 2.1 Hz, 1H), 4.35 – 4.14 (m, 3H), 3.63 – 3.49 (m, 1H), 3.43 – 3.12 (m, 5H), 2.88 (d, J = 3.6 Hz, 2H), 2.86 – 2.58 (m, 2H), 2.32 – 2.14 (m, 1H), 2.04 – 1.86 (m, 2H), 1.85 – 1.76 (m, 1H), 1.74 – 1.68 (m, 1H), 1.57 – 1.34 (m, 1H). 22/22 protons observed/predicted.

**$^{13}$C NMR** (101 MHz, DMSO-*d*$_6$) δ 151.44, 144.28, 144.11, 131.18, 131.10, 130.68, 130.61, 129.27, 129.21, 129.18, 127.70, 127.64, 91.87, 91.76, 74.70, 74.60, 52.22, 52.14, 46.77, 46.74, 46.59, 46.09, 41.89, 40.83, 40.15, 39.94, 39.73, 39.52, 39.31, 39.10, 38.89, 33.69, 33.59, 31.54, 31.52, 31.03, 30.88, 29.19, 29.14, 25.51, 24.48. Apparent mixture of two amide rotational isomers

**HRMS** (*m/z*): $[M+H]^+$ calculated for $C_{19}H_{23}Cl_2N_2O_3S$, 429.0801; found, 429.0796.

Preparation of VVD-118313 (**5a**)

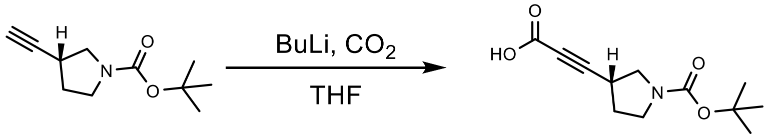

**S11a**

##### 3-[(3R)-1-tert-Butoxycarbonylpyrrolidin-3-yl]prop-2-ynoic acid (S11a)

To a solution of tert-butyl (3R)-3-ethynylpyrrolidine-1-carboxylate (1.20 g, 6.15 mmol, 1 eq.) in THF (12 mL) was added dropwise n-BuLi (1.2 eq., 3 mL, 2.5 M) at -78 °C. The mixture was stirred at -78 °C for 1 hour. Then to the mixture was added dry ice (5 eq., 1.5 g) at -78 °C. The mixture was stirred at -78 °C for 0.5 hour. Then the temperature of the reaction liquid was gradually raised to room temperature. The reaction mixture was quenched by water (30 mL), and then 2M hydrochloric acid was added to adjust the pH to 3 and extracted with ethyl acetate (3 x 10 mL). The combined organic layers were dried over $Na_2SO_4$, filtered and concentrated under reduced pressure to give 3-[(3R)-1-tert-butoxycarbonylpyrrolidin-3-yl]prop-2-ynoic acid (**S11a**) (1.45 g, 6.06 mmol, 99% yield) as a yellow oil.

**$^1$H NMR** (400 MHz, $CDCl_3$): δ = 3.30 - 3.73 (m, 4 H), 3.11 (quin, *J*=6.7 Hz, 1 H), 2.15 - 2.26 (m, 1 H), 2.05 -2.10 (m, 1 H), 1.47 (s, 9 H) ppm. 16/17 protons observed/expected

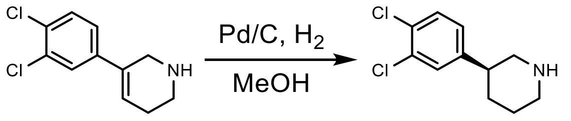

**S6** **S7a**

##### (3S)-3-(3,4-Dichlorophenyl)piperidine (S7a)

To a solution of 5-(3,4-dichlorophenyl)-1,2,3,6-tetrahydropyridine (**S6**) (1.00 g, 3.78 mmol) in MeOH (40 mL) was added Pd/C (10%, 300 mg) at 20 °C and the mixture was stirred at the same temperature for 1 h. The reaction was filtered and concentrated under reduced pressure to give a residue. The residue was purified by Chiral SFC (AD column, 5% to 40% methanol) to give (3S)-3-(3,4-dichlorophenyl)piperidine (**S7a**) (330 mg,1.43 mmol, 38% yield) as a white solid. The absolute stereochemistry of **7a** was assigned by x-ray crystallography (**S. I. Figure 1 and Table 1**).

**$^1$H NMR** (400 MHz, $CDCl_3$): δ = 7.66 – 8.13 (m, 1 H), 7.40 (d, *J*=8.3 Hz, 1 H), 7.31 (d, *J*=2.0 Hz, 1 H), 7.05 (dd, *J*=8.3, 2.1 Hz, 1 H), 3.35 – 3.55 (m, 2 H), 3.14 (tt, *J*=12.2, 3.4 Hz, 1 H), 2.73 – 2.92 (m, 2 H), 2.05 – 2.13 (m, 1 H), 1.92 – 2.04 (m, 2 H), 1.51 – 1.72 (m, 1 H) ppm. 13/13 protons observed/expected.

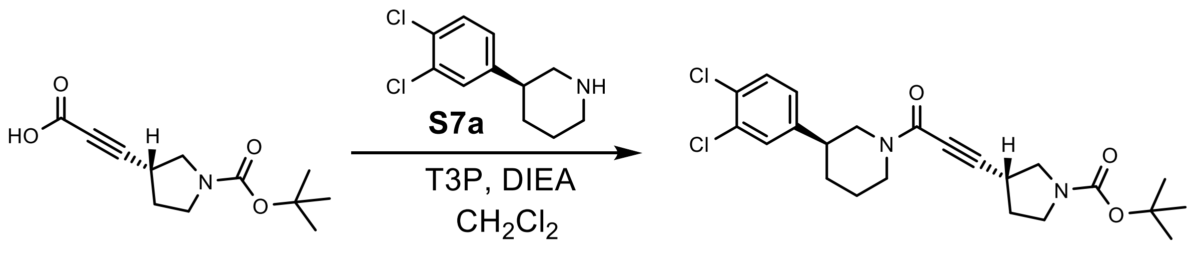

S11a S12a

**tert-Butyl (3S)-3-[3-oxo-3-[(3R)-3-(3,4-dichlorophenyl)-1-piperidyl]prop-1-ynyl]pyrrolidine-1-carboxylate (S12a)**

To a solution of 3-[(3R)-1-tert-butoxycarbonylpyrrolidin-3-yl]prop-2-ynoic acid (**S11a**) (390.0 mg, 1.630 mmol, 1 eq.), (3S)-3-(3,4-dichlorophenyl)piperidine (**S7a**) (375.1 mg, 1.630 mmol, 1 eq.) and DIEA (337.0 mg, 2.607 mmol, 3 eq.) in $CH_2Cl_2$ (3 mL) was added T3P (1.38 g, 2.173 mmol, 2.5 eq.) at 20 °C and the reaction was stirred at 20 °C for 5 min. TLC (1:1 petroleum ether:ethyl acetate; Rf starting material 0.00, product 0.40) showed the starting material was consumed completely. The reaction was poured into $H_2O$ (15 mL) and extracted with $CH_2Cl_2$ (3 x 5 mL). The organic layers were washed with brine (5 mL), dried over $Na_2SO_4$, filtered and concentrated under reduced pressure to give a residue. The residue was purified by flash chromatography (10-50% v/v ethyl acetate in petroleum ether) to give tert-butyl (3S)-3-[3-oxo-3-[(3R)-3-(3,4-dichlorophenyl)-1-piperidyl]prop-1-ynyl]pyrrolidine-1-carboxylate (**S12a**) (300.0 mg, 0.665 mmol, 76.5% yield) as a yellow solid.

**$^1$H NMR** (400 MHz, $CDCl_3$): δ 7.36 – 7.46 (m, 1 H), 7.33 (dd, *J*=3.9, 2.2 Hz, 1 H), 7.07 (dd, *J*=8.3, 1.8 Hz, 1 H), 4.53 – 4.72 (m, 1 H), 4.28 – 4.46 (m, 1 H), 3.25 – 3.84 (m, 4 H), 3.00 – 3.22 (m, 2 H), 2.55 – 2.80 (m, 2 H), 2.13 – 2.30 (m, 1 H), 2.06 – 2.12 (m, 1 H), 1.77 – 2.05 (m, 2 H), 1.59 – 1.76 (m, 2 H), 1.46 (d, *J*=13.2 Hz, 9 H) ppm. 28/28 protons observed/expected.

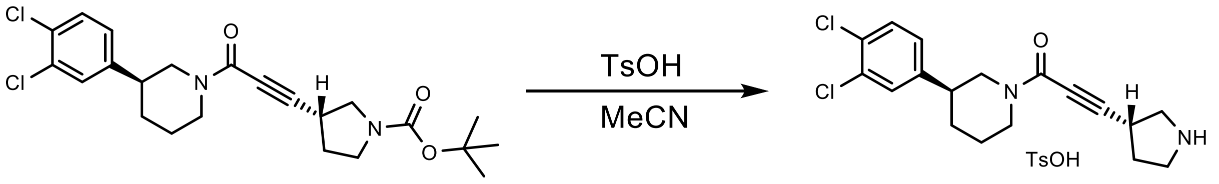

S12a S13a

**1-[(3S)-3-(3,4-Dichlorophenyl)-1-piperidyl]-3-[(3R)-pyrrolidin-3-yl]prop-2-yn-1-one (S13a)**

To a solution of tert-butyl (3R)-3-[3-oxo-3-[(3S)-3-(3,4-dichlorophenyl)-1-piperidyl]prop-1-ynyl]pyrrolidine-1-carboxylate (300.0 mg, 0.665 mmol, 1 eq.) (**S12a**) in MeCN (3 mL) was added TsOH (114.5 mg, 0.665 mmol, 1 eq.) at 20 °C and the mixture was stirred at 65 °C for 6 hrs. TLC (0:1 petroleum ether:ethyl acetate; Rf starting material 0.70, product 0.00) showed the starting material was consumed completely. The reaction was concentrated under reduced pressure to give 1-[(3S)-3-(3,4-dichlorophenyl)-1-piperidyl]-3-[(3R)-pyrrolidin-3-yl]prop-2-yn-1-one (**S13a**) (233 mg) as white solid which was carried forward crude without further purification.

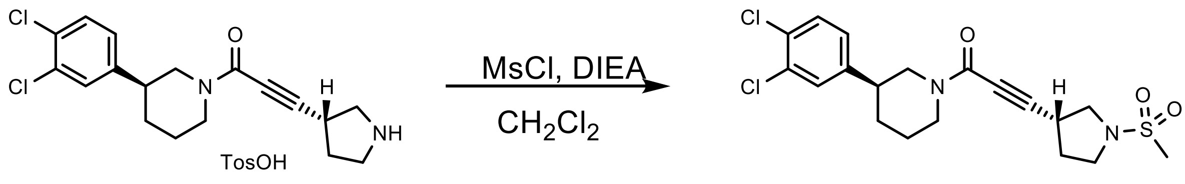


**S13a** **5a**

**1-[(3S)-3-(3,4-Dichlorophenyl)-1-piperidyl]-3-[(3R)-1-methylsulfonylpyrrolidin-3-yl]prop-2-yn-1-one (5a)**

To a solution of 1-[(3S)-3-(3,4-dichlorophenyl)-1-piperidyl]-3-[(3R)-pyrrolidin-3-yl]prop-2-yn-1-one (233.0 mg, 0.663 mmol, 1 eq.) (**S13a**) in $CH_2Cl_2$ (3 mL) was added DIEA (257.2 mg, 1.990 mmol, 3 eq.) and MsCl (91.2 mg, 0.796 mmol, 1.2 eq.) at 0 °C and the mixture was stirred at 0 °C for 0.5 hr. TLC (0:1 petroleum ether:ethyl acetate; Rf starting material 0.00, product 0.45) showed the starting material was consumed completely. The reaction was poured into $H_2O$ (6 mL) and extracted with $CH_2Cl_2$ (3 x 2 mL). The organic layers were washed with brine (2 mL), dried over $Na_2SO_4$, filtered and concentrated under reduced pressure to give a residue. The residue was purified by flash chromatography (10-70% v/v ethyl acetate in petroleum ether) and chiral HPLC (WHELK O1 IH column, liquid phase: [A-CO2$_2$; B-MeOH] 42% B isocratic) to give 1-[(3S)-3-(3,4-dichlorophenyl)-1-piperidyl]-3-[(3R)-1-methylsulfonylpyrrolidin-3-yl]prop-2-yn-1-one (48.6 mg, 0.112 mmol, 16.8% yield over two steps) (**5a**) as a white solid (ee >99%).

**$^1$H NMR** (400 MHz,DMSO-*d$_6$*): δ 7.64 – 7.55 (m, 2H), 7.31 (ddd, *J* = 10.6, 8.4, 2.1 Hz, 1H), 4.35 – 4.14 (m, 2H), 3.56 (ddd, *J* = 22.1, 9.8, 7.0 Hz, 1H), 3.44 – 3.12 (m, 6H), 2.91 (d, *J* = 26.3 Hz, 3H), 2.85 – 2.59 (m, 2H), 2.32 – 2.15 (m, 1H), 2.06 – 1.87 (m, 1H), 1.88 (s, 1H), 1.79 (ddd, *J* = 16.0, 8.1, 3.7 Hz, 1H), 1.74 (s, 1H) ppm. 22/22 protons observed/expected.

**$^{13}$C NMR** (101 MHz, DMSO-*d$_6$*) δ = 151.93, 144.76, 144.59, 131.66, 131.58, 131.17, 131.10, 129.75, 129.71, 129.69, 129.66, 128.19, 128.15, 128.12, 92.35, 92.25, 75.18, 75.08, 52.71, 52.62, 47.26, 47.06, 46.56, 42.39, 41.32, 34.20, 34.14, 34.10, 34.04, 32.03, 31.99, 31.52, 31.37, 29.67, 29.61, 25.99, 24.96. Apparent mixture of two amide rotational isomers

**HRMS** (*m/z*): $[M+H]^+$ calculated for $C_{19}H_{23}Cl_2N_2O_3S$, 429.0801; found, 429.0800.

$[\alpha]_D^{20}$ = -20 (c 0.01, $CHCl_3$)

Preparation of compound **5b**

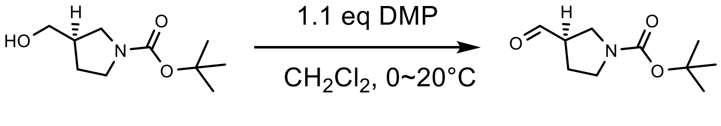


**S14a**

**tert-Butyl (3R)-3-formylpyrrolidine-1-carboxylate (S14)**

To a solution of (R)-3-hydroxymethyl-pyrrolidine-1-carboxylic acid tert-butyl ester (2.00 g, 9.937 mmol, 1.0 eq.) in $CH_2Cl_2$ (40 mL, 0.2484 M) was added Dess–Martin periodinane (4.64 g, 10.931 mmol, 1.1 eq.) at 0 °C and the mixture was stirred at 20 °C for 3 hours. TLC (1:1 petroleum ether:ethyl acetate; Rf: starting material 0.4, product 0.5) showed the reaction was complete. 20 mL of 10% $Na_2S_2O_3$ was added to the mixture and stirred at 20 °C for 0.5 hour. The mixture was separated. The organic phase was washed with sat. $NaHCO_3$ (80 mL) and brine (80 mL), dried over $Na_2SO_4$, filtered, concentrated and purified by flash chromatography (17-67% v/v ethyl acetate in petroleum ether) to give tert-butyl (3R)-3-formylpyrrolidine-1-carboxylate (**S14**) (1.60 g, 8.030 mmol, 80.8% yield) as a colorless oil. Spectral characterization matched those found in the literature[4].

**$^1$H NMR** (400 MHz, $CDCl_3$): δ = 9.70 (d, *J*=1.5 Hz, 1 H), 3.24 - 3.81 (m, 4 H), 3.03 (br s, 1 H), 2.13 (s, 2 H), 1.47 (s, 9 H) ppm. 17/17 protons observed/expected.

**tert-Butyl (3S)-3-ethynylpyrrolidine-1-carboxylate** (**S15a**)

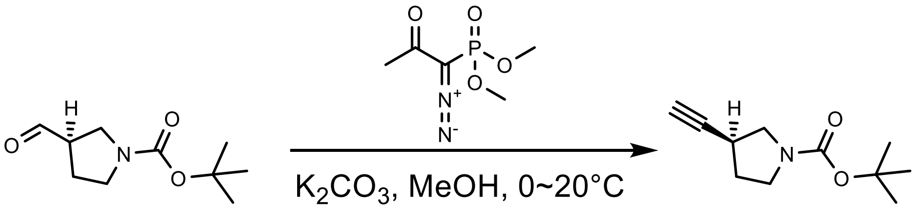


**S14a** **S15a**

To a solution of tert-butyl (3R)-3-formylpyrrolidine-1-carboxylate (**S14a**) (1.50 g, 7.528 mmol, 1.0 eq.) in methanol (15 mL, 0.5019 M) was added dimethyl (1-diazo-2-oxopropyl)phosphonate (1.88 g, 9.787 mmol, 1.3 eq.), followed by $K_2CO_3$ (2.40 g, 2.3 eq., 17.31 mmol) at 0 °C and the reaction was warmed to 20 °C over 1 hour. TLC (5:1 petroleum ether:ethyl acetate; Rf: starting material 0.2, product 0.5) showed the reaction was complete. The mixture was diluted with ethyl acetate (20 mL) and filtered. The filtrate was concentrated and purified by flash chromatography (5-10% v/v ethyl acetate in petroleum ether) to give tert-butyl (3S)-3-ethynylpyrrolidine-1-carboxylate (**S15**) (1.15 g, 5.890 mmol, 78.2% yield) as a white solid.

**$^1$H NMR** (400 MHz, $CDCl_3$): δ = 3.42 - 3.72 (m, 2 H), 3.19 - 3.40 (m, 2 H), 2.86 - 3.03 (m, 1 H), 2.08 - 2.25 (m, 2 H), 1.87 - 2.02 (m, 1 H), 1.35 - 1.56 (m, 9 H) ppm. 17/17 protons observed/expected.

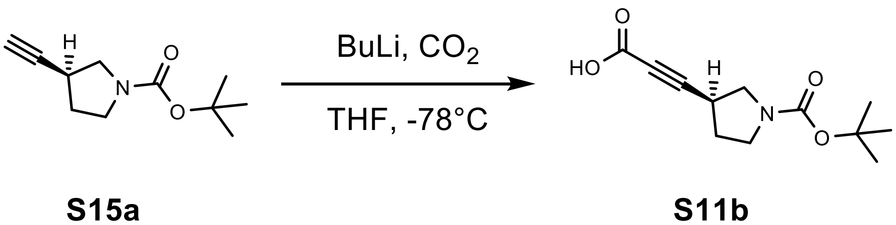


**S15a** **S11b**

**3-[(3S)-1-tert-Butoxycarbonylpyrrolidin-3-yl]prop-2-ynoic acid (S11b)**

To a solution of tert-butyl (3S)-3-ethynylpyrrolidine-1-carboxylate (**S15a**) (600.0 mg, 3.073 mmol, 1.0 eq.) in THF (5.63 mL, 0.5463 M) was added dropwise n-BuLi (1.2 eq., 0.16 mL) at -78 °C. The mixture was stirred at -78 °C for 1 hour. To the resulting mixture was then added dry ice (5.0 eq., 1.5 g) at -78 °C. The mixture was stirred at -78 °C for 0.5 hour. The temperature of the reaction liquid was then gradually raised to room temperature. The reaction mixture was quenched with water (10 mL), and then 2M hydrochloric acid was added to adjust the pH to 3 and the solution was extracted with ethyl acetate (3 x 3 mL). The combined organic layers were dried over $Na_2SO_4$, filtered and concentrated under reduced pressure to give 3-[(3S)-1-tert-butoxycarbonylpyrrolidin-3-yl]prop-2-ynoic acid (**S11b**) (700.0 mg, 2.926 mmol, 95.2% yield) as a yellow oil.

**$^1$H NMR** (400 MHz, $CDCl_3$): δ 3.30 - 3.73 (m, 4 H), 3.11 (quin, *J*=6.7 Hz, 1 H), 2.15 - 2.26 (m, 1 H), 2.05 - 2.10 (m, 1 H), 1.47 (s, 9 H) ppm. 16/17 protons observed/expected.

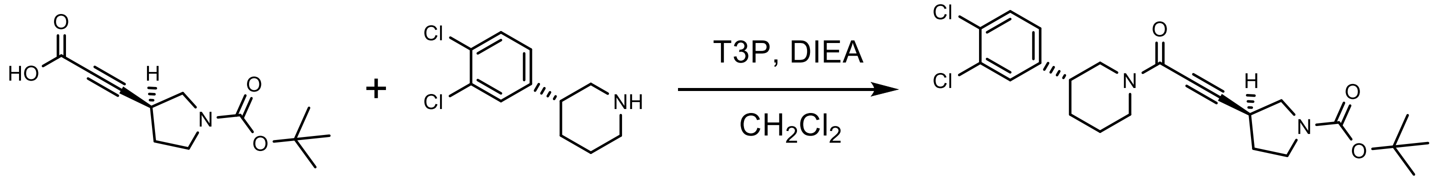


**S11b** **S7b** **S12b**

**tert-Butyl (3S)-3-[3-oxo-3-[(3R)-3-(3,4-dichlorophenyl)-1-piperidyl]prop-1-ynyl]pyrrolidine-1-carboxylate (S12b)**

To a solution of 3-[(3S)-1-tert-butoxycarbonylpyrrolidin-3-yl]prop-2-ynoic acid (**S11b**) (207.9 mg, 0.869 mmol, 1.0 eq.), (3R)-3-(3,4-dichlorophenyl)piperidine (**S7b**) (200.0 mg, 0.869 mmol, 1.0 eq.) and DIEA (337 mg, 2.61 mmol, 3 eq.) in $CH_2Cl_2$ (5mL, 0.17 M) was added T3P (1.38 g, 2.173 mmol, 2.5 eq.) at 20 °C and the reaction was stirred at 20 °C for 0.5 hr. TLC (1:1 petroleum ether:ethyl acetate; Rf: starting material 0, product 0.4) showed the starting material was consumed completely. The reaction was poured into water (15 mL) and extracted with $CH_2Cl_2$ (3 x 5 mL). The organic layers were washed with brine (5 mL), dried over $Na_2SO_4$, filtered and concentrated under reduced pressure to give a residue. The residue was purified by flash chromatography (0-33% v/v ethyl acetate in petroleum ether) to give tert-butyl (3S)-3-[3-oxo-3-[(3R)-3-(3,4-dichlorophenyl)-1-piperidyl]prop-1-ynyl]pyrrolidine-1-carboxylate (**S12b**) (300 mg, 0.665 mmol, 76.5% yield) as a white solid.

**$^1$H NMR** (400 MHz, $CDCl_3$): δ 7.36 - 7.45 (m, 1 H), 7.30 - 7.35 (m, 1 H), 7.06 (dd, *J*=2.2, 8.3 Hz, 1 H), 4.56 - 4.72 (m, 1 H), 4.34 (td, *J*=2.0, 13.2 Hz, 1 H), 3.25 - 3.74 (m, 4 H), 2.99 - 3.21 (m, 2 H), 2.58 - 2.75 (m, 2 H), 2.06 - 2.28 (m, 2 H), 1.78 - 1.94 (m, 1 H), 1.52 - 1.75 (m, 2 H), 1.45 (d, *J*=13.2 Hz, 9 H) ppm. 27/28 protons observed/expected.

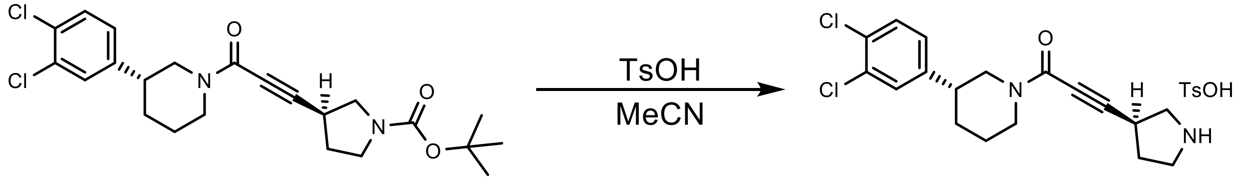


**S12b** **S13b**

**4-Methylbenzenesulfonic acid;1-[ (3R)-3-(3,4-dichlorophenyl)-1-piperidyl]-3-[(3S)-pyrrolidin-3-yl]prop-2-yn-1-one (S13b)**

To a solution of tert-butyl (3S)-3-[3-oxo-3-[(3R)-3-(3,4-dichlorophenyl)-1-piperidyl]prop-1-ynyl]pyrrolidine-1-carboxylate (300.0 mg, 0.665 mmol, 1 eq.) (**S12b**) was added TsOH (125.9 mg, 0.731 mmol, 1.1 eq.) at 25 °C and the mixture was warmed to 65 °C and stirred for 6 hours. TLC (1:1 petroleum ether:ethyl acetate; Rf: starting material 0.4, product 0) showed the reaction was complete. The mixture was concentrated to give 4-methylbenzenesulfonic acid;1-[ (3R)-3-(3,4-dichlorophenyl)-1-piperidyl]-3-[(3S)-pyrrolidin-3-yl]prop-2-yn-1-one (**S13b**) (340 mg) as a white solid which was carried forward crude without further purification.

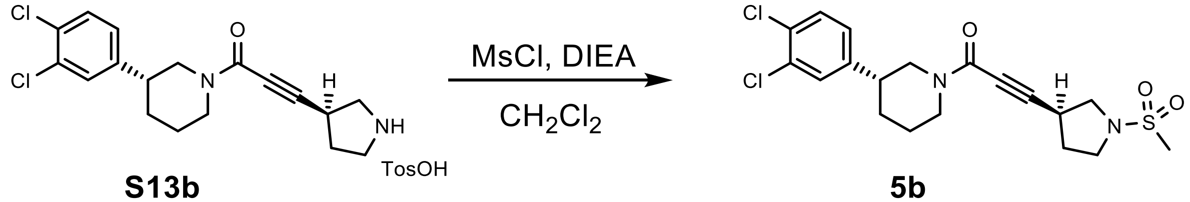


**S13b** **5b**

**1-[(3R)-3-(3,4-Dichlorophenyl)-1-piperidyl]-3-[(3S)-1-methylsulfonylpyrrolidin-3-yl]prop-2-yn-1-one (5b)**

To a solution of 1-[(3R)-3-(3,4-dichlorophenyl)-1-piperidyl]-3-[(3S)-pyrrolidin-3-yl]prop-2-yn-1-one 4-methylbenzenesulfonic acid salt (**S13b**) (340.0 mg, 0.650 mmol, 1.0 eq.) in $CH_2Cl_2$ (5 mL, 0.13 M) was added DIEA dropwise until the pH reached 7. Then to the mixture was added MsCl (111.6 mg, 0.970 mmol, 1.5 eq.) and the mixture was stirred at 20 °C for 5 min. TLC (10:1 dichloromethane:methanol; Rf: starting material 0.5, product 0.8) showed the reaction was complete. The mixture was quenched with water (10 mL) and extracted with $CH_2Cl_2$ (2 x 10 mL). The organic phases were washed with brine (20 mL), dried over $Na_2SO_4$, filtered, concentrated and purified by flash chromatography (67-100% v/v ethyl acetate in petroleum ether) to give a crude product. The crude product was purified by prep-chiral SFC (Chiralpak IE column; liquid phase: [A-CO2; B-isopropanol] 50% B isocratic) to give 1-[(3R)-3-(3,4-dichlorophenyl)-1-piperidyl]-3-[(3S)-1-methylsulfonylpyrrolidin-3-yl]prop-2-yn-1-one (**5b**) (37.5 mg, 0.081 mmol, 12.5% yield) as a white solid (ee 99%).

**$^{1}$H NMR** (600 MHz, CDCl3) δ 7.40 (dd, J = 31.9, 8.3 Hz, 1H), 7.31 (dd, J = 7.6, 2.2 Hz, 1H), 7.07 (ddd, J = 10.5, 8.3, 2.1 Hz, 1H), 4.66 – 4.55 (m, 1H), 4.32 (dddt, J = 14.7, 5.9, 3.8, 2.1 Hz, 1H), 3.65 (ddd, J = 20.4, 10.2, 7.0 Hz, 1H), 3.55 – 3.48 (m, 1H), 3.48 – 3.41 (m, 2H), 3.37 (dd, J = 10.2, 6.0 Hz, 1H), 3.21 (dp, J = 25.4, 6.7 Hz, 1H), 3.14 – 3.05 (m, 1H), 2.90 (s, 2H), 2.85 (s, 1H), 2.73 – 2.59 (m, 2H), 2.29 (tq, J = 19.9, 6.9 Hz, 1H), 2.21 – 2.03 (m, 3H), 1.88 (ddt, J = 30.0, 13.5, 3.0 Hz, 1H), 1.73 – 1.51 (m, 1H). 23/23 protons observed/expected.

**$^{13}$C NMR** (101 MHz, DMSO-*d$_6$*) δ = 151.44, 144.28, 144.11, 131.18, 131.10, 130.68, 130.61, 129.27, 129.21, 129.18, 127.70, 127.64, 91.87, 91.76, 74.70, 74.60, 52.22, 52.14, 46.77, 46.74, 46.59, 46.09, 41.89, 40.83, 40.15, 39.94, 39.73, 39.52, 39.31, 39.10, 38.89, 33.69, 33.59, 31.54, 31.52, 31.03, 30.88, 29.19, 29.14, 25.51, 24.48. Apparent mixture of two amide rotational isomers

**HRMS** (*m/z*): $[M+H]^+$ calculated for $C_{19}H_{23}Cl_2N_2O_3S$, 429.0801; found, 429.0802.

$[\alpha]_D^{20}$ = +10 (c 0.01, $CHCl_3$)

Preparation of compound **5c**

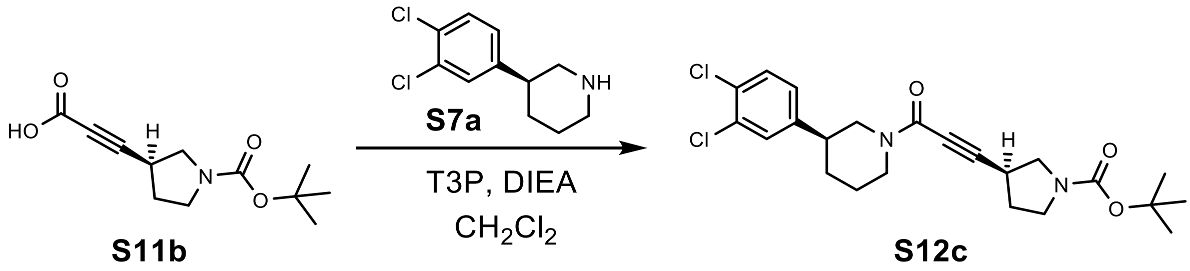


**tert-Butyl (3S)-3-[3-oxo-3-[(3S)-3-(3,4-dichlorophenyl)-1-piperidyl]prop-1-ynyl]pyrrolidine-1-carboxylate (S12c)**

To a solution of 3-[(3S)-1-tert-butoxycarbonylpyrrolidin-3-yl]prop-2-ynoic acid (**S11b**) (207.9 mg, 0.869 mmol, 1 eq.), (3S)-3-(3,4-dichlorophenyl)piperidine (200.0 mg, 0.869 mmol, 1 eq.) (**S7a**) and DIEA (337 mg, 2.61 mmol, 3 eq.) in $CH_2Cl_2$ (3 mL) was added T3P (1.38 g, 2.173 mmol, 2.5 eq.) at 20 °C and the reaction was stirred at 20 °C for 1hr. TLC (1:1 petroleum ether:ethyl acetate; Rf starting material 0.00, product 0.40) showed the starting material was consumed completely. The reaction was poured into $H_2O$ (15 mL) and extracted with $CH_2Cl_2$ (3 x 5 mL). The organic layers were washed with brine (5 mL), dried over $Na_2SO_4$, filtered and concentrated under reduced pressure to give a residue. The residue was purified by flash chromatography (10-50% v/v ethyl acetate in petroleum ether) to give tert-butyl (3S)-3-[3-oxo-3-[(3S)-3-(3,4-dichlorophenyl)-1-piperidyl]prop-1-ynyl]pyrrolidine-1-carboxylate (**S12c**) (300.0 mg, 0.665 mmol, 76.5% yield) as a yellow solid.

**$^1$H NMR** (400 MHz, $CDCl_3$): δ 7.36 - 7.46 (m, 1 H), 7.33 (dd, *J*=3.9, 2.2 Hz, 1 H), 7.07 (dd, *J*=8.3, 1.8 Hz, 1 H), 4.53 - 4.72 (m, 1 H), 4.28 - 4.46 (m, 1 H), 3.25 - 3.84 (m, 4 H), 3.00 - 3.22 (m, 2 H), 2.55 - 2.80 (m, 2 H), 2.13 - 2.30 (m, 1 H), 2.06 - 2.12 (m, 1 H), 1.77 - 2.05 (m, 2 H), 1.59 - 1.76 (m, 2 H), 1.46 (d, *J*=13.2 Hz, 9 H) ppm. 28/28 protons observed/expected.

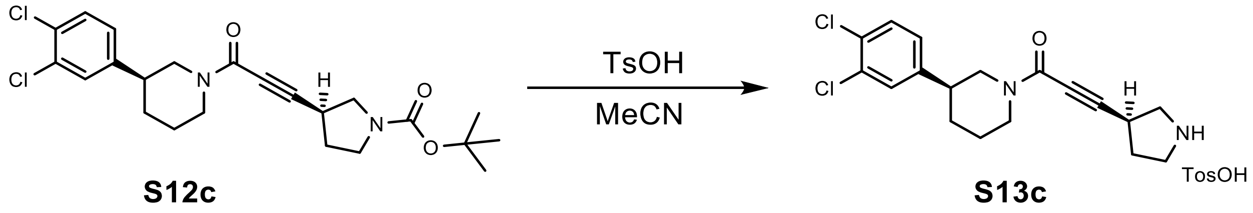


**1-[(3S)-3-(3,4-Dichlorophenyl)-1-piperidyl]-3-[(3S)-pyrrolidin-3-yl]prop-2-yn-1-one 4-methyl benzene sulfonate salt (S13c)**

To a solution of tert-butyl (3S)-3-[3-oxo-3-[(3S)-3-(3,4-dichlorophenyl)-1-piperidyl]prop-1-ynyl]pyrrolidine-1-carboxylate (**S12c**) (300.0 mg, 0.665 mmol, 1 eq.) in MeCN (3 mL) was added 4-methylbenzenesulfonic acid (114.5 mg, 0.665 mmol, 1 eq.) at 20 °C and the mixture was stirred at 65 °C for 6 hrs. TLC (0:1 petroleum ether:ethyl acetate; Rf:starting material 0.70, product 0.00) showed the starting material was consumed completely. The reaction was concentrated under reduced pressure to give 1-[(3S)-3-(3,4-dichlorophenyl)-1-piperidyl]-3-[(3S)-

pyrrolidin-3-yl]prop-2-yn-1-one 4-methyl benzene sulfonate salt (**S13c**) (233.0 mg) as a colorless oil which was carried forward crude without further purification.

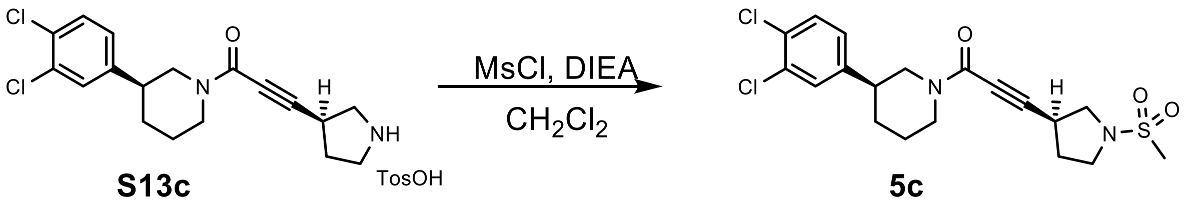


**S13c** **5c**

**1-[(3S)-3-(3,4-Dichlorophenyl)-1-piperidyl]-3-[(3S)-1-methylsulfonylpyrrolidin-3-yl]prop-2-yn-1-one (5c)**

To a solution of 1-[(3S)-3-(3,4-dichlorophenyl)-1-piperidyl]-3-[(3S)-pyrrolidin-3-yl]prop-2-yn-1-one 4-methylbenzene sulfonate salt (**S13c**) (233.0 mg, 0.663 mmol, 1 eq.) in $CH_2Cl_2$ (3 mL) was added DIEA (257 mg, 1.99 mmol, 3 eq.) and MsCl (91.2 mg, 0.796 mmol, 1.2 eq.) at 0 °C and the mixture was stirred at 0 °C for 0.5 hr. TLC (0:1 petroleum ether:ethyl acetate; Rf starting material 0.00, product 0.45) showed the starting material was consumed completely. The reaction was poured into $H_2O$ (6 mL) and extracted with $CH_2Cl_2$ (3 x 2 mL). The organic layers were washed with brine (2 mL), dried over $Na_2SO_4$, filtered and concentrated under reduced pressure to give a residue. The residue was purified by flash chromatography (10-70% v/v ethyl acetate in petroleum ether) and chiral SFC (ChiralPak IH column, liquid phase: [A-CO2; B-isopropanol] 42% B isocratic) to give 1-[(3S)-3-(3,4-dichlorophenyl)-1-piperidyl]-3-[(3S)-1-methylsulfonylpyrrolidin-3-yl]prop-2-yn-1-one (**5c**) (117.9 mg, 0.275 mmol, 41.4% yield) as a white solid (ee 99%).

**$^1$H NMR** (600 MHz, CDCl3) δ 7.36 (d, J = 8.3 Hz, 0H), 7.30 (dd, J = 10.3, 2.2 Hz, 1H), 7.06 (ddd, J = 12.4, 8.3, 2.2 Hz, 1H), 4.63 – 4.52 (m, 1H), 4.31 (dddd, J = 14.9, 7.5, 3.4, 1.6 Hz, 1H), 3.63 (ddd, J = 19.6, 10.2, 7.0 Hz, 1H), 3.50 (ddd, J = 9.9, 7.6, 6.4 Hz, 1H), 3.47 – 3.39 (m, 2H), 3.36 (dd, J = 10.2, 5.9 Hz, 1H), 3.22 (p, J = 6.6 Hz, 1H), 3.17 (q, J = 6.6 Hz, 1H), 3.13 – 3.03 (m, 1H), 2.88 (s, 2H), 2.84 (s, 1H), 2.67 (dddd, J = 16.0, 12.5, 8.5, 3.5 Hz, 1H), 2.64 – 2.59 (m, 1H), 2.34 – 2.21 (m, 1H), 2.14 (ddt, J = 12.7, 7.7, 6.3 Hz, 1H), 2.07 – 2.01 (m, 1H), 1.85 (ddq, J = 30.7, 13.2, 3.1 Hz, 1H), 1.72 – 1.49 (m, 1H). 22/22 protons observed/expected.

**$^{13}$C NMR** (101 MHz, DMSO-$d_6$) δ = 151.44, 144.28, 144.11, 131.18, 131.10, 130.68, 130.61, 129.27, 129.21, 129.18, 127.70, 127.64, 91.87, 91.76, 74.70, 74.60, 52.22, 52.14, 46.77, 46.74, 46.59, 46.09, 41.89, 40.83, 40.15, 39.94, 39.73, 39.52, 39.31, 39.10, 38.89, 33.69, 33.59, 31.54, 31.52, 31.03, 30.88, 29.19, 29.14, 25.51, 24.48. Apparent mixture of two amide rotational isomers

**HRMS** (*m/z*): $[M+H]^+$ calculated for $C_{19}H_{23}Cl_2N_2O_3S$, 429.0801; found, 429.0791.

$[\alpha]_D^{20}$ = +110 (c 0.01, $CHCl_3$)

Preparation of compound **5d**

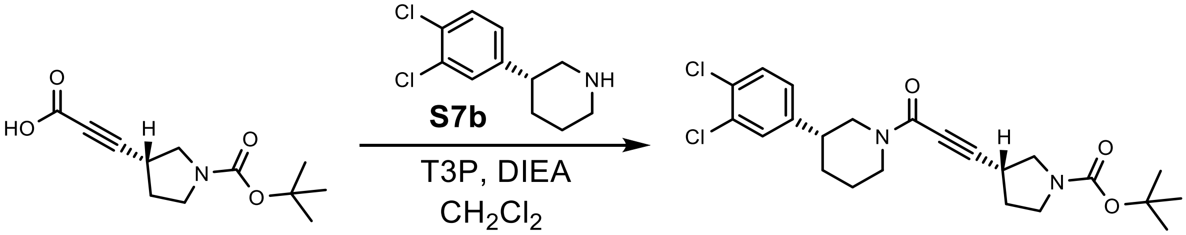


**S11a** **S12d**

**tert-Butyl (3R)-3-[3-oxo-3-[(3R)-3-(3,4-dichlorophenyl)-1-piperidyl]prop-1-ynyl]pyrrolidine-1-carboxylate (S12d)**

To a solution of 3-[(3R)-1-tert-butoxycarbonylpyrrolidin-3-yl]prop-2-ynoic acid (**S11a**) (390.0 mg, 1.630 mmol, 1 eq.), (3R)-3-(3,4-dichlorophenyl)piperidine (**S7b**) (375.1 mg, 1.630 mmol, 1 eq.) and DIEA (337.0 mg, 2.607 mmol, 3 eq.) in $CH_2Cl_2$ (3 mL) was added T3P (1.38 g, 2.173 mmol, 2.5 eq.) at 20 °C and the reaction was stirred at 20 °C for 5 min. TLC (1:1 petroleum ether:ethyl acetate; Rf starting material 0.00, product 0.40) showed the starting material was consumed completely. The reaction was poured into $H_2O$ (15 mL) and extracted with $CH_2Cl_2$ (3 x 5 mL). The organic layers were washed with brine (5 mL), dried over $Na_2SO_4$, filtered and concentrated under reduced pressure to give a residue. The residue was purified by flash chromatography (10-50% v/v ethyl acetate in petroleum ether) to give tert-butyl (3R)-3-[3-oxo-3-[(3R)-3-(3,4-dichlorophenyl)-1-piperidyl]prop-1-ynyl]pyrrolidine-1-carboxylate (**S12d**) (300.0 mg, 0.665 mmol, 76.5% yield) as a yellow solid.

**$^1$H NMR** (400 MHz, $CDCl_3$) δ = 7.36-7.47 (m, 1 H), 7.29-7.35 (m, 1 H), 7.07 (dd, *J*=8.3, 1.8 Hz, 1 H), 4.53-4.72 (m, 1 H), 4.35 (br d, *J*=13.1 Hz, 1 H), 3.24-3.81 (m, 4 H), 2.98-3.22 (m, 2 H), 2.51-2.81 (m, 2 H), 2.12-2.33 (m, 1 H), 2.09 (br d, *J*=3.9 Hz, 1 H), 1.79-2.04 (m, 2 H), 1.53-1.76 (m, 2 H), 1.46 (d, *J*=13.0 Hz, 9 H) ppm. 28/28 protons observed/expected.

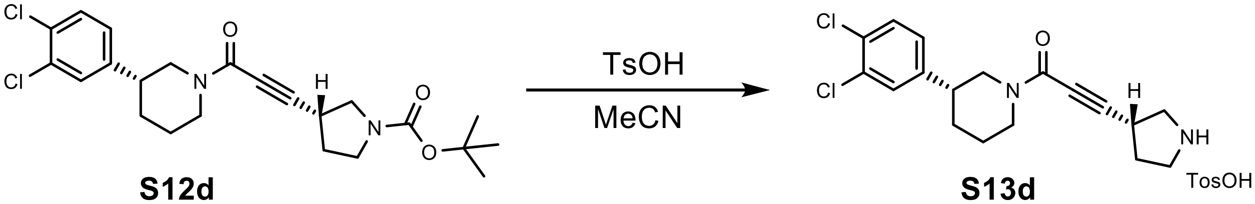


**S12d** **S13d**

**1-[(3R)-3-(3,4-Dichlorophenyl)-1-piperidyl]-3-[(3R)-pyrrolidin-3-yl]prop-2-yn-1-one 4-methylbenzene sulfonate salt (S13d)**

To a solution of tert-butyl (3R)-3-[3-oxo-3-[(3R)-3-(3,4-dichlorophenyl)-1-piperidyl]prop-1-ynyl]pyrrolidine-1-carboxylate (**S12d**) (300.0 mg, 0.665 mmol, 1 eq.) in MeCN (3 mL) was added TsOH (114.5 mg, 0.665 mmol, 1 eq.) at 20 °C and the mixture was stirred at 65 °C for 1.5 hrs. TLC (0:1 petroleum ether:ethyl acetate; Rf starting material 0.70, product 0.00) showed the starting material was consumed completely. The reaction was concentrated under reduced pressure to give 1-[(3R)-3-(3,4-dichlorophenyl)-1-piperidyl]-3-[(3R)-pyrrolidin-3-yl]prop-2-yn-1-one 4-methylbenzene sulfonate salt (**S13d**) (230.0 mg) as a white solid which was carried forward crude without further purification.

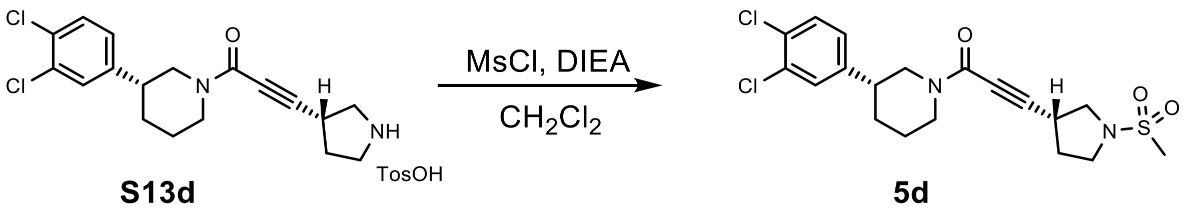

**1-[(3R)-3-(3,4-Dichlorophenyl)-1-piperidyl]-3-[(3R)-1-methylsulfonylpyrrolidin-3-yl]prop-2-yn-1-one (5d)**

To a solution of 1-[(3R)-3-(3,4-dichlorophenyl)-1-piperidyl]-3-[(3R)-pyrrolidin-3-yl]prop-2-yn-1-one (**S13d**) (233.0 mg, 0.663 mmol, 1 eq.) in $CH_2Cl_2$ (3 mL) was added DIEA (257.2 mg, 1.990 mmol, 3 eq.) and MsCl (91.2 mg, 0.796 mmol, 1.2 eq.) at 0 °C and the mixture was stirred at 0 °C for 0.5 hr. TLC (0:1 petroleum ether:ethyl acetate; Rf starting material 0.00, product 0.45) showed the starting material was consumed completely. The reaction was poured into $H_2O$ (6 mL) and extracted with $CH_2Cl_2$ (3 x 2 mL). The organic layers were washed with brine (2 mL), dried over $Na_2SO_4$, filtered and concentrated under reduced pressure to give a residue. The residue was purified by flash chromatography (10-70% v/v ethyl acetate in petroleum ether) and chiral SFC (ChiralPak IH column, liquid phase: [A-CO2; B-isopropanol] 42% B isocratic) to give 1-[(3R)-3-(3,4-dichlorophenyl)-1-piperidyl]-3-[(3R)-1-methylsulfonylpyrrolidin-3-yl]prop-2-yn-1-one (**5d**) (25.5 mg, 0.059 mmol, 8.9% yield) as a white solid.

**$^1$H NMR** (600 MHz, CDCl3) δ 7.40 (dd, J = 31.9, 8.3 Hz, 1H), 7.31 (dd, J = 7.6, 2.1 Hz, 1H), 7.07 (ddd, J = 10.5, 8.3, 2.2 Hz, 1H), 4.66 – 4.55 (m, 1H), 4.36 – 4.29 (m, 1H), 3.65 (ddd, J = 20.3, 10.2, 7.0 Hz, 1H), 3.55 – 3.46 (m, 1H), 3.48 – 3.41 (m, 2H), 3.37 (dd, J = 10.2, 6.0 Hz, 1H), 3.21 (dp, J = 25.3, 6.7 Hz, 1H), 3.14 – 3.05 (m, 1H), 2.90 (s, 2H), 2.85 (s, 1H), 2.73 – 2.59 (m, 2H), 2.29 (tq, J = 20.0, 6.9 Hz, 1H), 2.22 – 2.03 (m, 2H), 1.88 (dddd, J = 26.7, 13.6, 6.1, 3.0 Hz, 1H), 1.73 – 1.51 (m, 2H). 22/22 protons observed/expected.

**$^{13}$C NMR** (101 MHz, DMSO-*d₆*) δ = 151.44, 144.28, 144.11, 131.18, 131.10, 130.68, 130.61, 129.27, 129.21, 129.18, 127.70, 127.64, 91.87, 91.76, 74.70, 74.60, 52.22, 52.14, 46.77, 46.74, 46.59, 46.09, 41.89, 40.83, 40.15, 39.94, 39.73, 39.52, 39.31, 39.10, 38.89, 33.69, 33.59, 31.54, 31.52, 31.03, 30.88, 29.19, 29.14, 25.51, 24.48. Apparent mixture of two amide rotational isomers

**HRMS** (*m/z*): $[M+H]^+$ calculated for $C_{19}H_{23}Cl_2N_2O_3S$, 429.0801; found, 429.0801.

$[\alpha]_D^{20}$ = -40 (c 0.01, $CHCl_3$)

<u>Preparation of compound **6**</u>

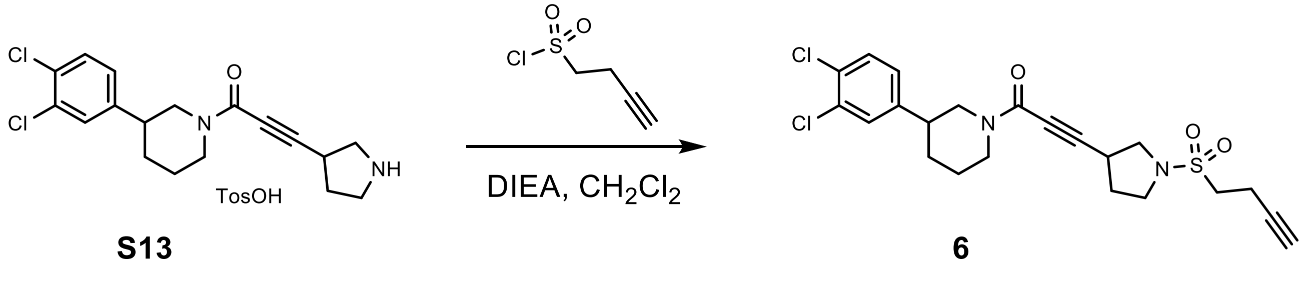

**3-(1-but-3-ynylsulfonylpyrrolidin-3-yl)-1-[3-(3,4-dichlorophenyl)-1-Piperidyl]prop-2-yn-1-one (6)**

But-3-yne-1-sulfonyl chloride (139.9 mg, 0.917 mmol, 1.2 eq.) was added to a mixture of 1-[3-(3,4-dichlorophenyl)-1-piperidyl]-3-pyrrolidin-3-yl-prop-2-yn-1-one 4-methylbenzenesulfonate salt (**S13**) (400.0 mg, 0.764 mmol, 1 eq.) and DIEA (296.3 mg, 2.292 mmol, 3 eq.) in $CH_2Cl_2$ (5 mL, 0.15 M) at 0 °C. The resulting mixture was stirred at 0 °C for 30 min. The reaction mixture was poured into water (10 mL) and extracted with $CH_2Cl_2$ (3 x 10 mL). The combined organic layer was washed with brine (10 mL), dried over $Na_2SO_4$, filtered and concentrated under reduced pressure to give a residue. The residue was purified by MPLC (ISCO; 4 g SepaFlash Silica Flash Column, Eluent of 0~50% Ethyl acetate/Petroleum ether gradient @ 18 mL/min) and prep-HPLC: column: Phenomenex Luna C18 75*30mm*3um; mobile phase: [water (0.1% TFA) - ACN]; B%: 35% - 65%, 8 min. 3-(1-but-3-ynylsulfonylpyrrolidin-3-yl)-1-[3-(3,4-dichlorophenyl)-1-piperidyl]prop-2-yn-1-one (**6**) (55 mg, 0.118 mmol, 15.4% yield) was obtained as pale yellow solid (ee 99%).

**$^1$H NMR** (400 MHz, DMSO-*d₆*) δ 7.65 – 7.56 (m, 2H), 7.33 (ddd, J = 10.5, 8.3, 2.1 Hz, 1H), 4.36 – 4.16 (m, 2H), 3.70 – 3.56 (m, 1H), 3.49 – 3.14 (m, 7H), 3.00 (dt, J = 12.8, 2.7 Hz, 1H), 2.93 – 2.76 (m, 1H), 2.80 – 2.62 (m, 1H), 2.65 – 2.58 (m, 1H), 2.57 (ddd, J = 10.0, 5.0, 2.0 Hz, 1H), 2.34 – 2.16 (m, 1H), 2.11 – 1.92 (m, 1H), 1.96 – 1.88 (m, 1H), 1.79 (dddd, J = 20.9, 18.7, 9.4, 5.9 Hz, 2H), 1.59 – 1.35 (m, 1H). 24/24 protons observed/expected.

**$^{13}$C NMR** (101 MHz, DMSO-$d_6$) δ 151.44, 144.28, 144.08, 131.19, 131.10, 130.67, 130.60, 129.28, 129.22, 129.18, 127.69, 127.67, 127.65, 91.73, 91.71, 91.68, 81.11, 81.08, 74.66, 74.56, 72.64, 72.60, 52.17, 51.91, 51.78, 46.75, 46.73, 46.66, 46.64, 46.60, 46.45, 46.10, 41.89, 40.85, 40.83, 40.15, 39.94, 39.73, 39.52, 39.31, 39.10, 38.89, 31.60, 31.04, 30.84, 29.31, 29.23, 25.52, 24.47, 13.07, 13.02. Apparent mixture of two amide rotational isomers.

**HRMS** (*m/z*): $[M+H]^+$ calculated for $C_{22}H_{25}Cl_2N_2O_3S$, 467.0957; found, 467.0949.

#### Small Molecule Crystallography

Crystals of compound **S7a** were grown from slow evaporation of a solution of the compound (20 mg) in a 1:3 mixture of methanol:acetonitrile. Data were collected on a Rigaku Oxford Diffraction XtaLAB Synergy four-circle diffractometer equipped with a HyPix-6000HE area detector and Oxford Cryostream 800 cooling system, generating Cu-Kα radiation ($\lambda$ = 1.54184 Å, 50 W) with a micro focus source with multi-layer mirror (μ-CMF), at a distance 35 mm, tube voltage 50 kV and tube current (1 mA). A total of 11703 reflections were collected in the 2θ range from 9.686 to 133.132. The limiting indices were: $-11 \leq h \leq 9$, $-8 \leq k \leq 8$, $-12 \leq l \leq 12$; which yielded 2285 unique reflections ($R_{int}$= 0.0472). Structures were solved using SHELX-T[5] then refined using SHELX-L[5]. The total number of refined parameters was 136, compared with 2285 data. All reflections were included in the refinement. The goodness of fit on $F^2$ was 1.099 with a final R value for $[I > 2\sigma (I)]$ $R_1$= 0.0410 and $wR_2$= 0.1036. The largest differential peak and hole were 0.32 and -0.25 Å$^{-3}$, respectively.

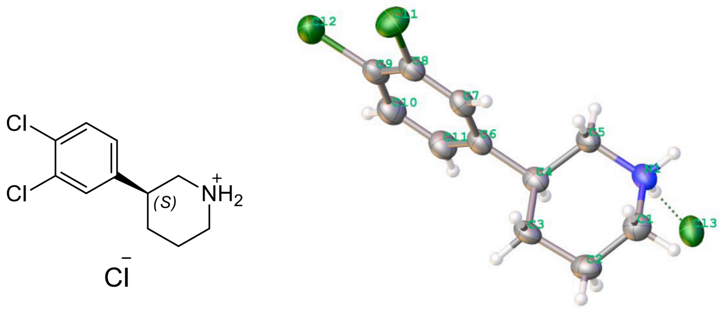


**Figure 1.** Compound **7a** absolute configuration (left) and ORTEP structure (right).

**Table 1.** Small molecule crystallography data collection and refinement statistics for **7a**

| | |
|---|---|
| Crystal size/mm$^3$ | 0.10 × 0.10 × 0.10 |
| Radiation Type | CuKα (λ = 1.54184) |
| Crystal system | monoclinic |
| Space group | $P2_1$ |
| a/Å | 9.8233(6) |
| b/Å | 7.3199(3) |
| c/Å | 10.3323(6) |
| α/° | 90 |
| β/° | 117.889(8) |
| γ/° | 90 |
| Cell Volume/Å$^3$ | 656.66(8) |
| Cell Formula Units Z | 2 |
| Crystal Density $_{calc}$ g/cm$^3$ | 1.348 |
| Crystal F(000) | 276.0 |
| Absorption Coefficient μ/mm$^{-1}$ | 6.058 |
| Index ranges | -11 ≤ h ≤ 9, -8 ≤ k ≤ 8, -12 ≤ l ≤ 12 |
| Cell Measurement Temperature/K | 150.00(10) |
| 2θ range for data collection/° | 9.686 to 133.132 |
| Goodness-of-fit on $F_2$ | 1.099 |
| Final R indexes [I>=2σ (I)] | $R_1$= 0.0410, $wR_2$= 0.1036 |
| Final R indexes [all data] | $R_1$= 0.0417, $wR_2$= 0.1040 |
| Largest diff. peak/hole/e Å$^{-3}$ | 0.32/-0.25 |
| Reflections collected/unique | 11703/2285 [$R_{int}$= 0.0472] |
| Flack parameter | 0.015(16) |

#### NMR Spectra of Final Compounds (1, 2, 3, 4, 5, 5a, 5b, 5c, 5d, 6)

##### Compound 1: $^{1}H$ and $^{13}C$ NMR in DMSO-$d_6$

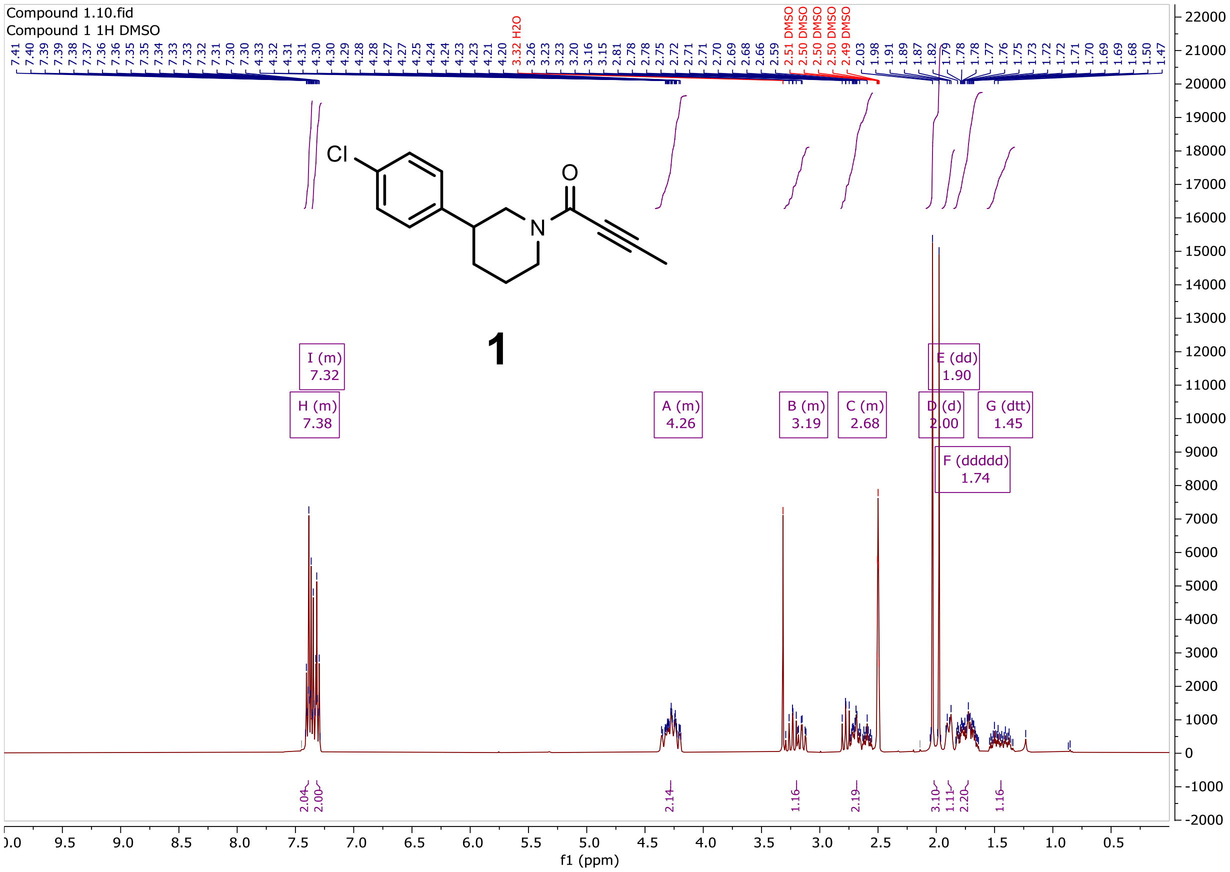

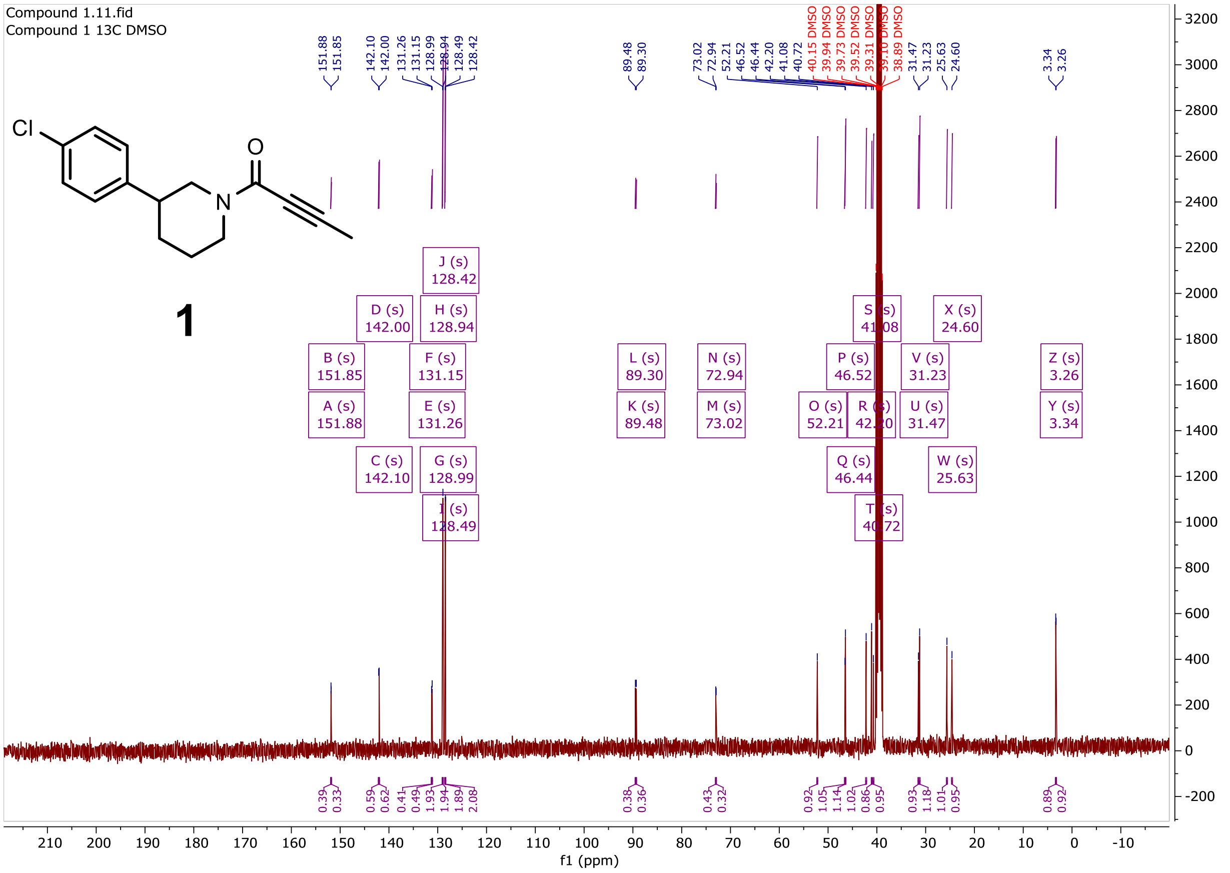
Compound 1.11.fid
Compound 1 13C DMSO
1
f1 (ppm)

#### Compound 2a: $^{1}$H and $^{13}$C NMR in DMSO-$d_6$

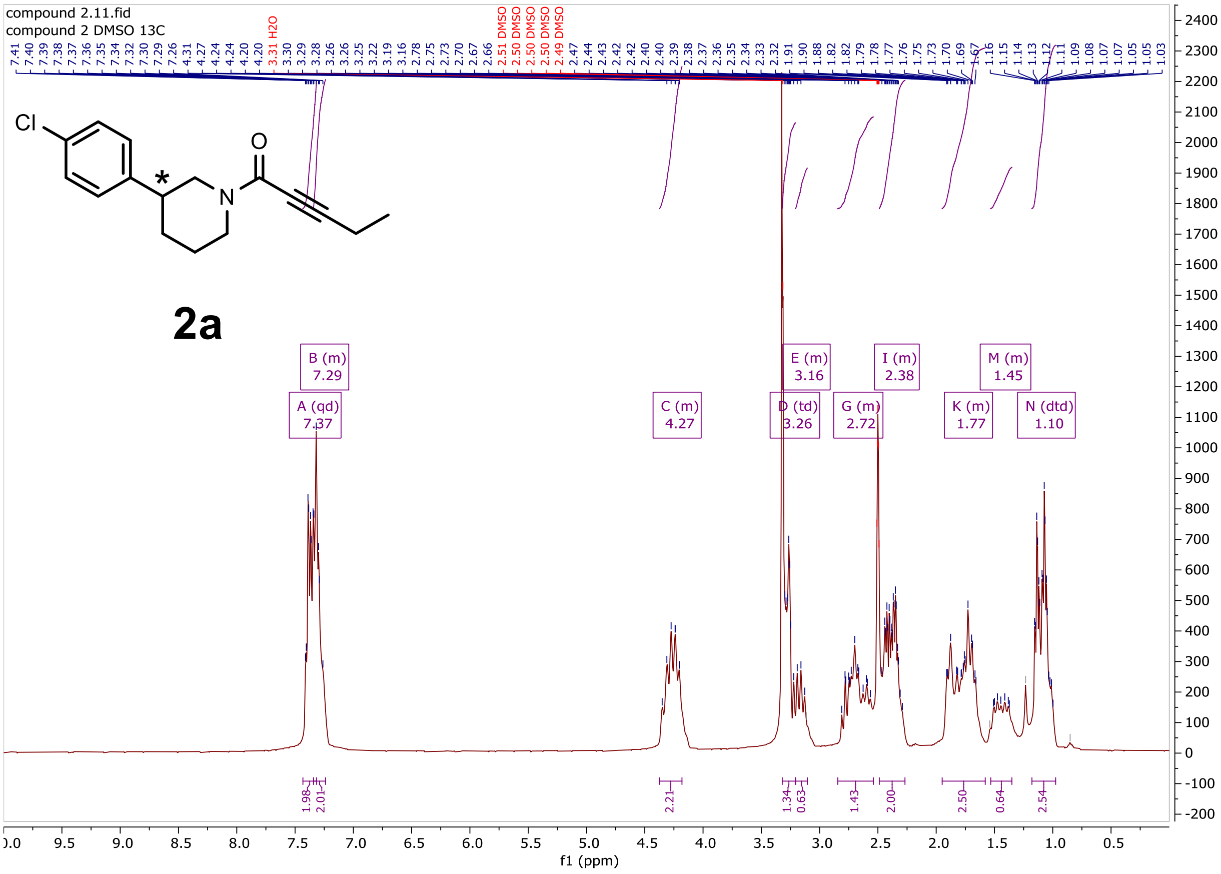

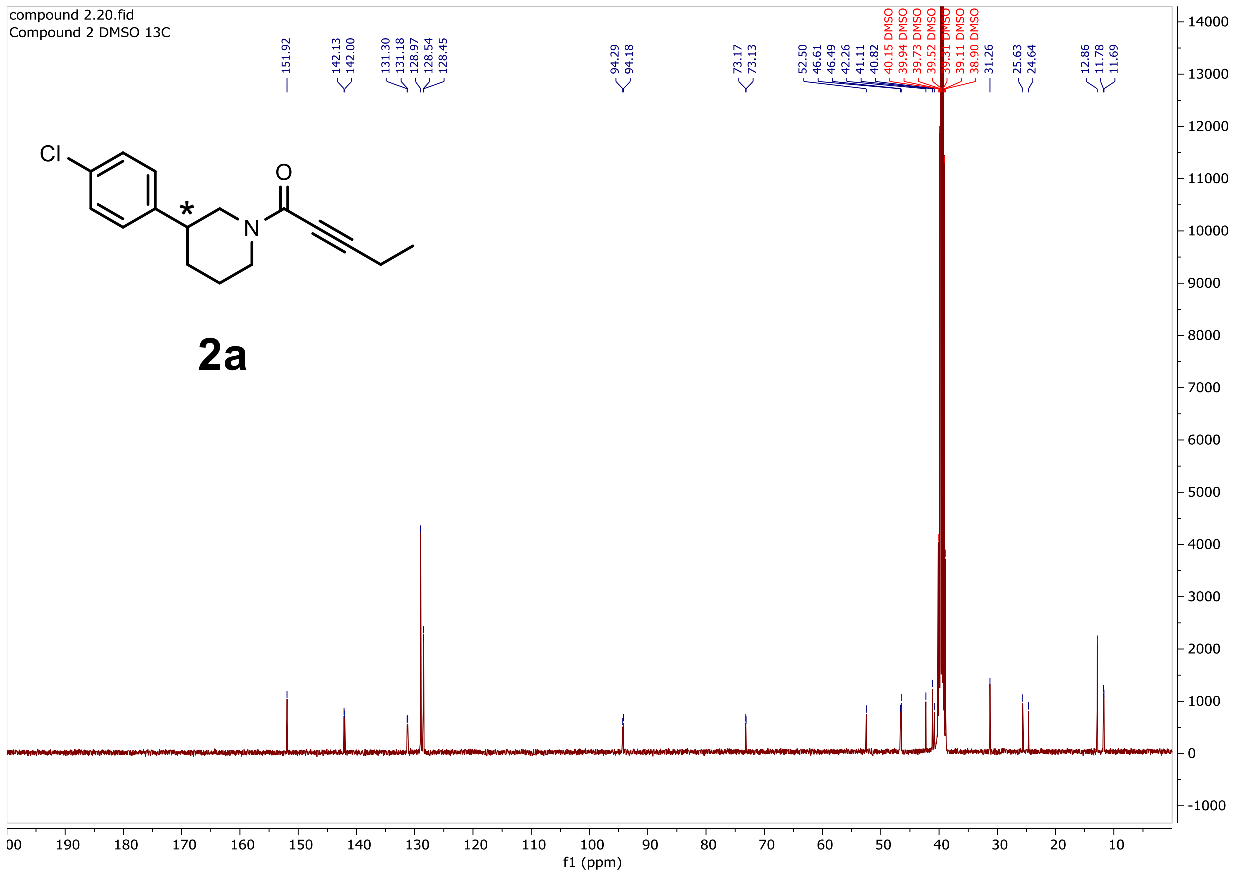
compound 2.20.fid
Compound 2 DMSO 13C
151.92
142.13
142.00
131.30
131.18
128.97
128.54
128.45
94.29
94.18
73.17
73.13
52.50
46.61
46.49
42.26
41.11
40.82
40.15 DMSO
39.94 DMSO
39.73 DMSO
39.52 DMSO
39.31 DMSO
39.11 DMSO
38.90 DMSO
31.26
25.63
24.64
12.86
11.78
11.69
Cl
O
N
*
f1 (ppm)

**2a**

Compound **2a** SFC trace

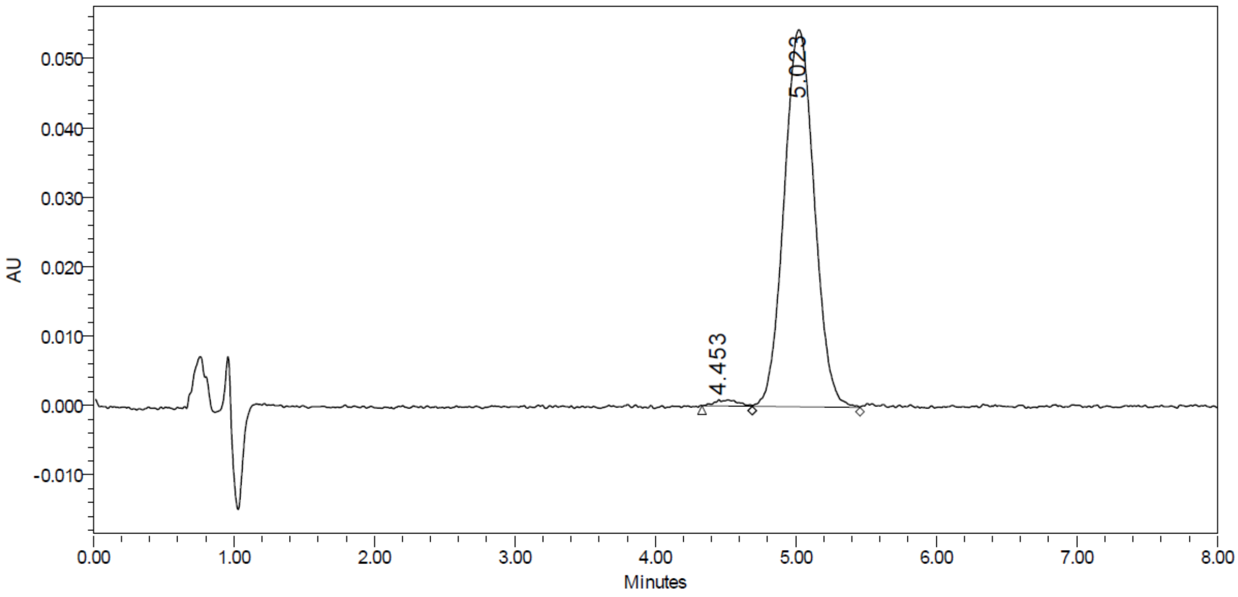


| | RT | Area | % Area | Height |
|---|---|---|---|---|
| 1 | 4.453 | 10154 | 1.24 | 884 |
| 2 | 5.023 | 811693 | 98.76 | 54286 |

#### Compound 3a: $^1$H and $^{13}$C NMR in DMSO-$d_6$

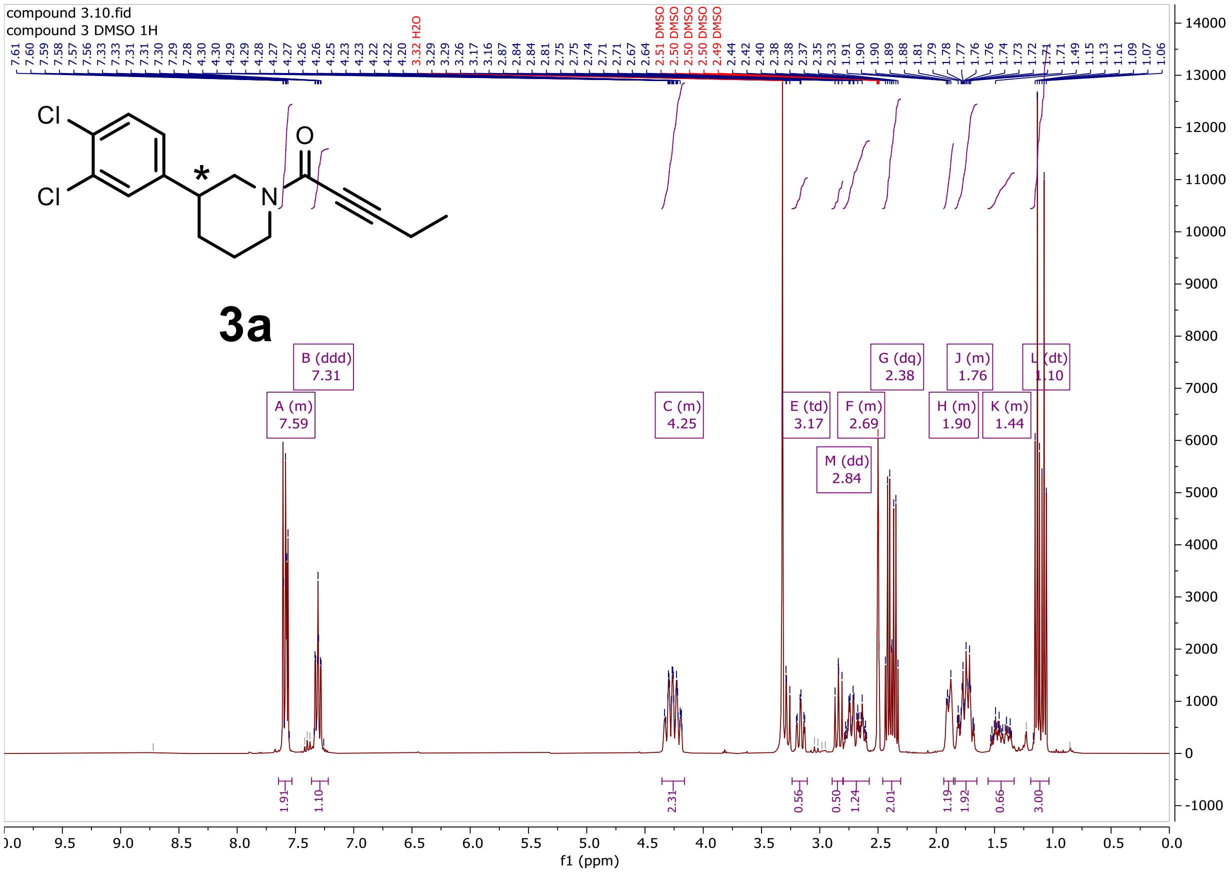

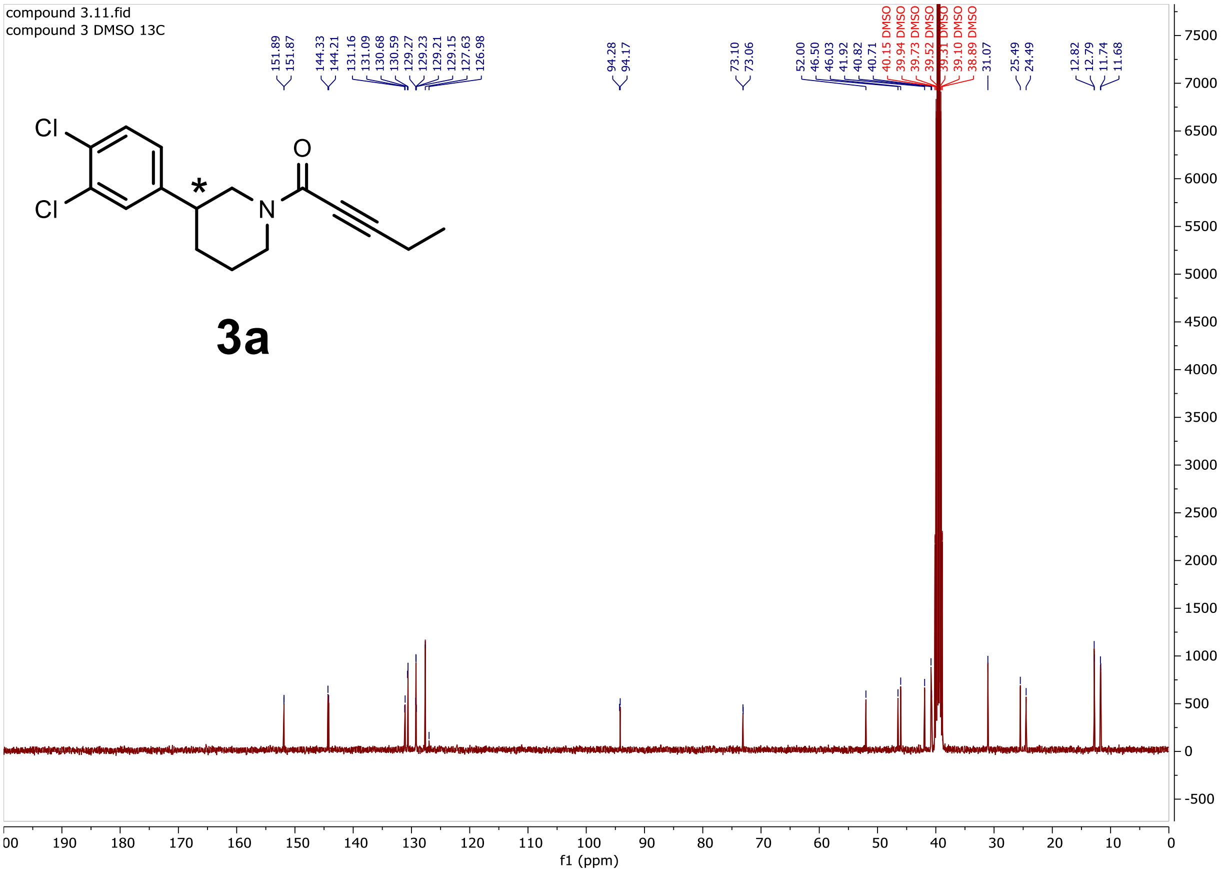
compound 3.11.fid
compound 3 DMSO 13C
151.89
151.87
144.33
144.21
131.16
131.09
130.68
130.59
129.27
129.23
129.21
129.15
127.63
126.98
94.28
94.17
73.10
73.06
52.00
46.50
46.03
41.92
40.82
40.71
40.15 DMSO
39.94 DMSO
39.73 DMSO
39.52 DMSO
39.31 DMSO
39.10 DMSO
38.89 DMSO
31.07
25.49
24.49
12.82
12.79
11.74
11.68
Cl
Cl
O
N
*
3a
f1 (ppm)

SFC of compound **3a**:

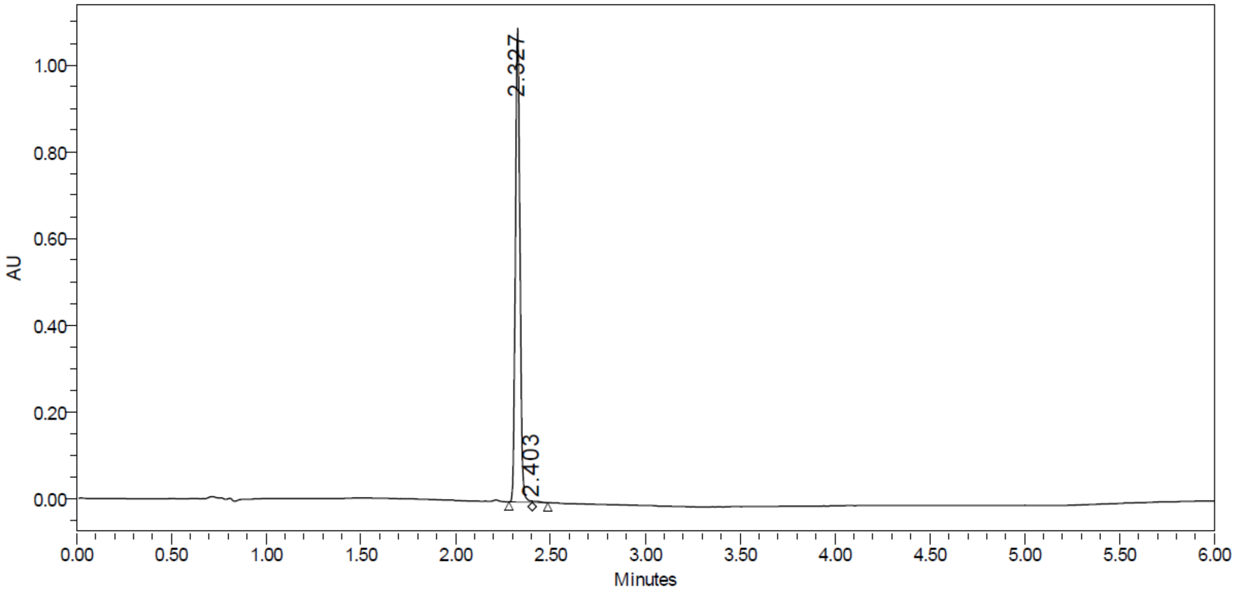


| | RT | Area | % Area | Height |
|---|---|---|---|---|
| 1 | 2.327 | 1824677 | 99.57 | 1090756 |
| 2 | 2.403 | 7947 | 0.43 | 2919 |

#### Compound 4: $^{1}H$ and $^{13}C$ NMR in $CDCl_3$

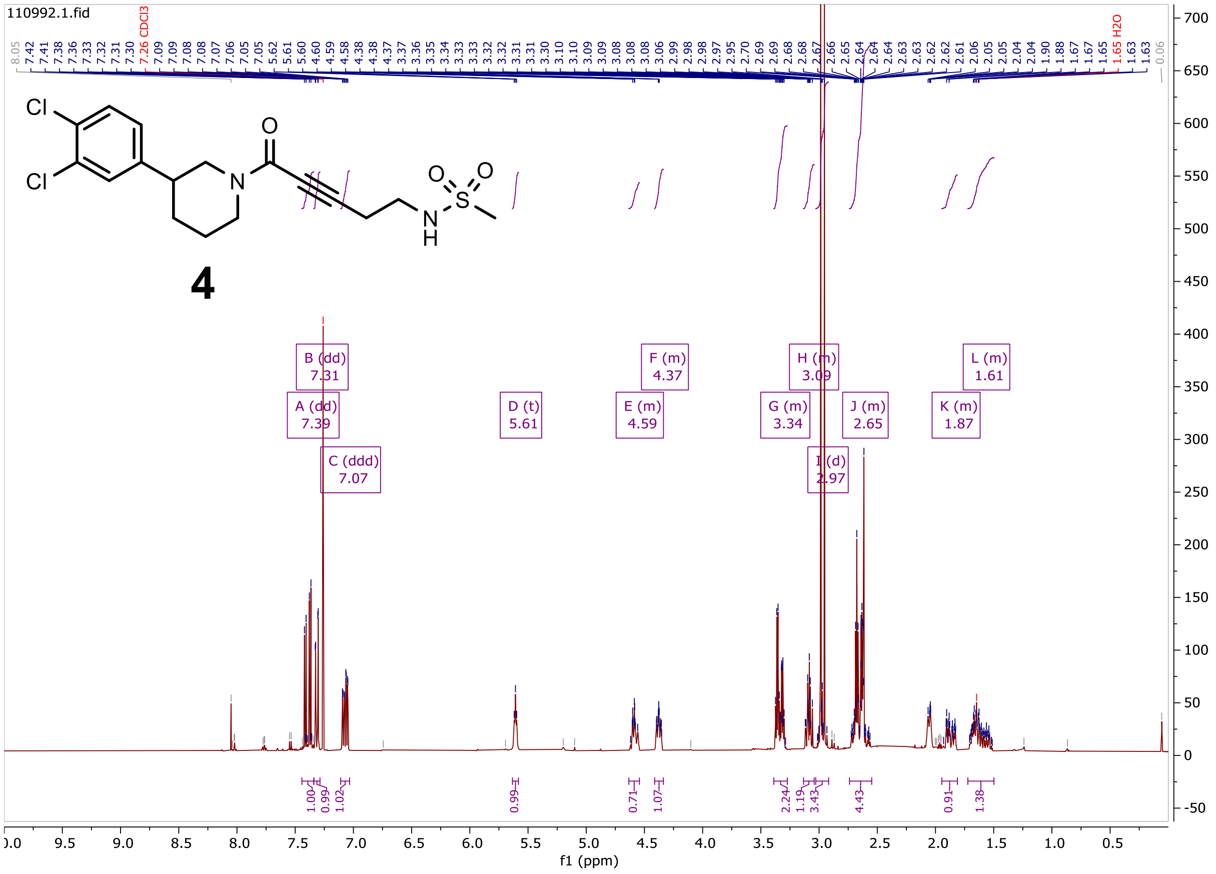

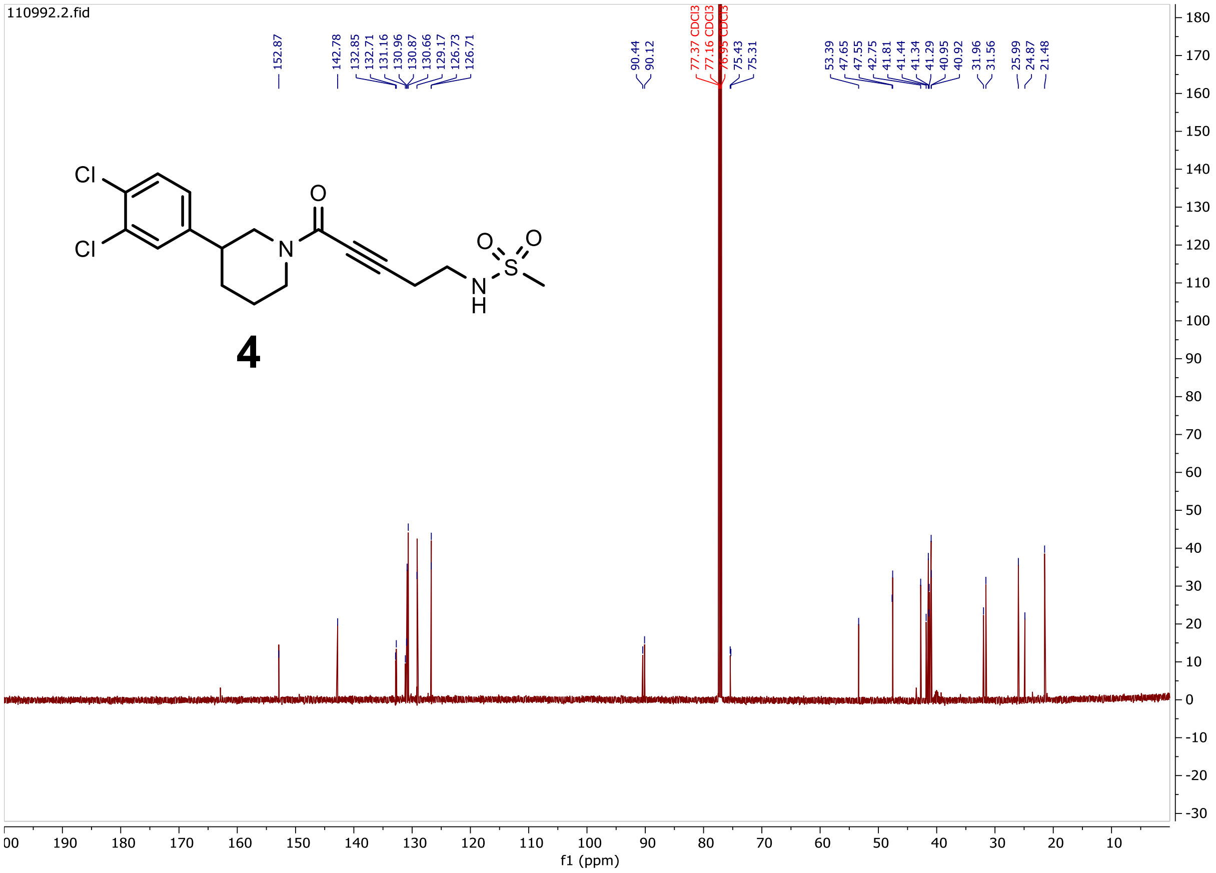

110992.2.fid
152.87
142.78
132.85
132.71
131.16
130.96
130.87
130.66
129.17
126.73
126.71
90.44
90.12
77.37 CDCl3
77.16 CDCl3
76.95 CDCl3
75.43
75.31
53.39
47.65
47.55
42.75
41.81
41.44
41.34
41.29
40.95
40.92
31.96
31.56
25.99
24.87
21.48
Cl
Cl
N
O
N
H
S
O
O
4
f1 (ppm)

#### Compound 5: $^{1}H$ and $^{13}C$ NMR in DMSO-$d_6$

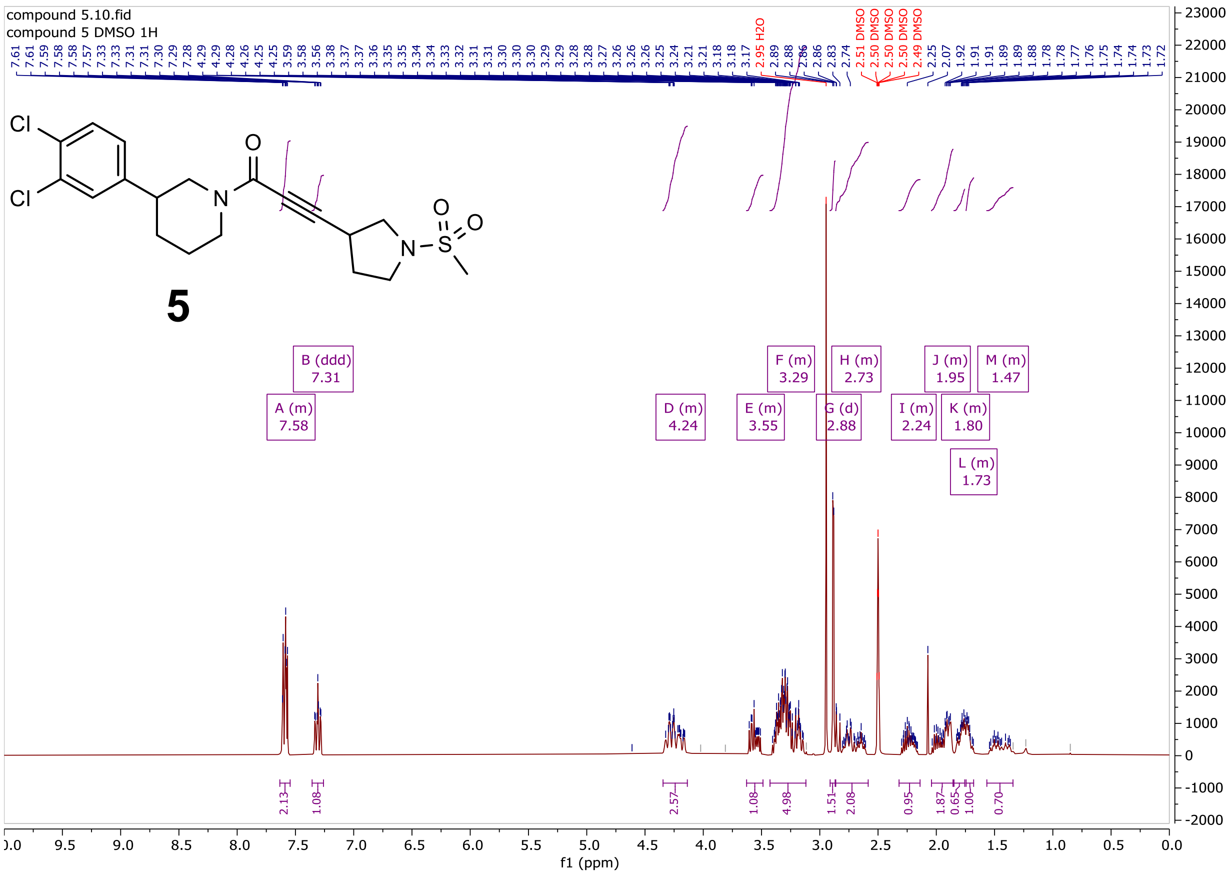

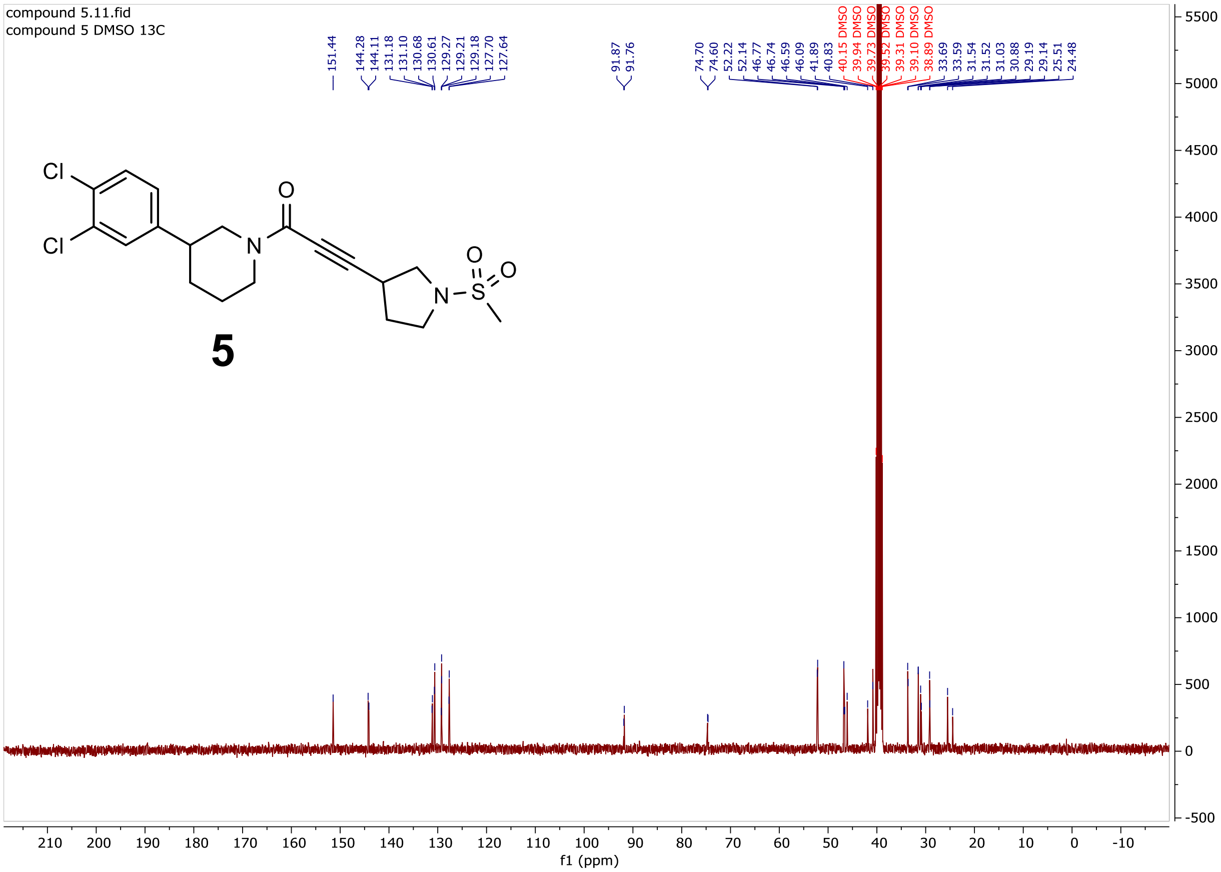
compound 5.11.fid
compound 5 DMSO 13C
151.44
144.28
144.11
131.18
131.10
130.68
130.61
129.27
129.21
129.18
127.70
127.64
91.87
91.76
74.70
74.60
52.22
52.14
46.77
46.74
46.59
46.09
41.89
40.83
40.15 DMSO
39.94 DMSO
39.73 DMSO
39.52 DMSO
39.31 DMSO
39.10 DMSO
38.89 DMSO
33.69
33.59
31.54
31.52
31.03
30.88
29.19
29.14
25.51
24.48
5
f1 (ppm)

**Compound 5a: $^{1}$H and $^{13}$C NMR in DMSO-$d_6$**

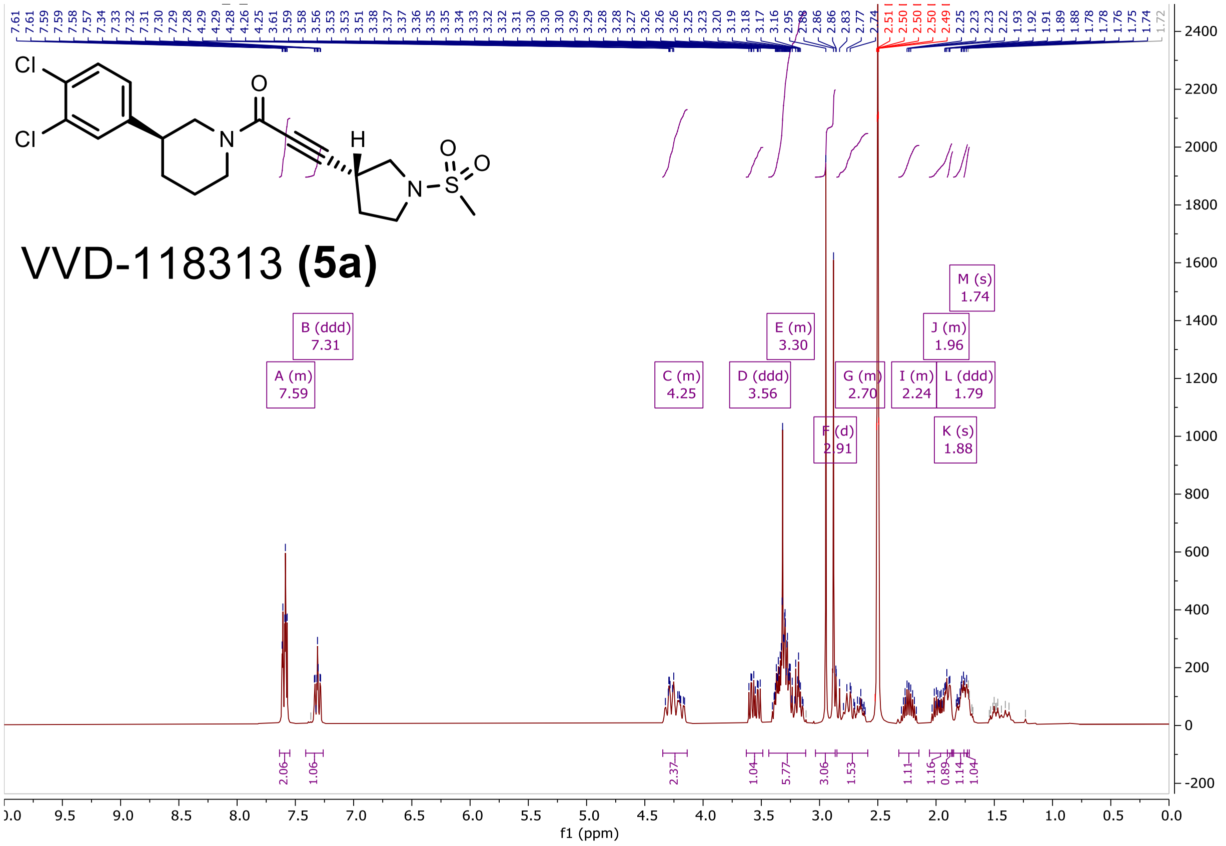

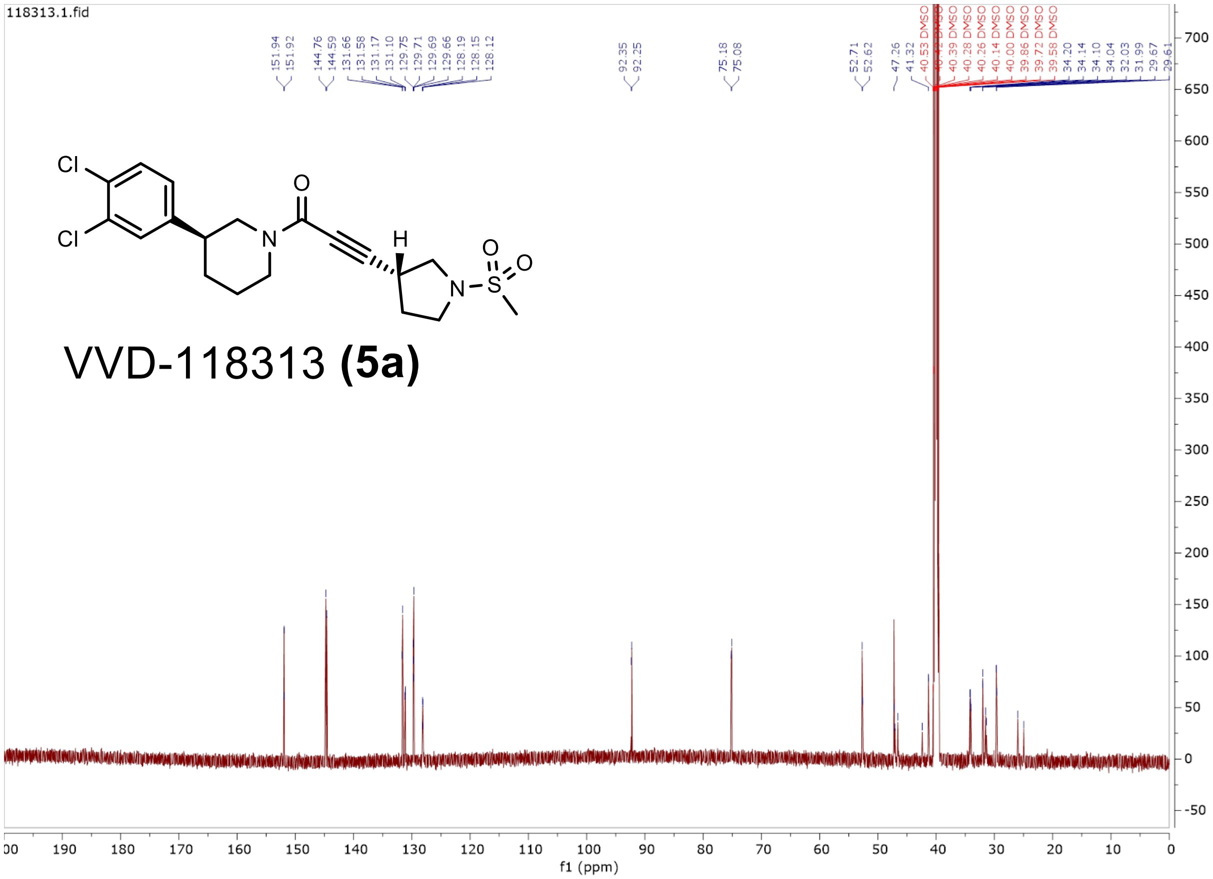


SFC of compound **5a**:

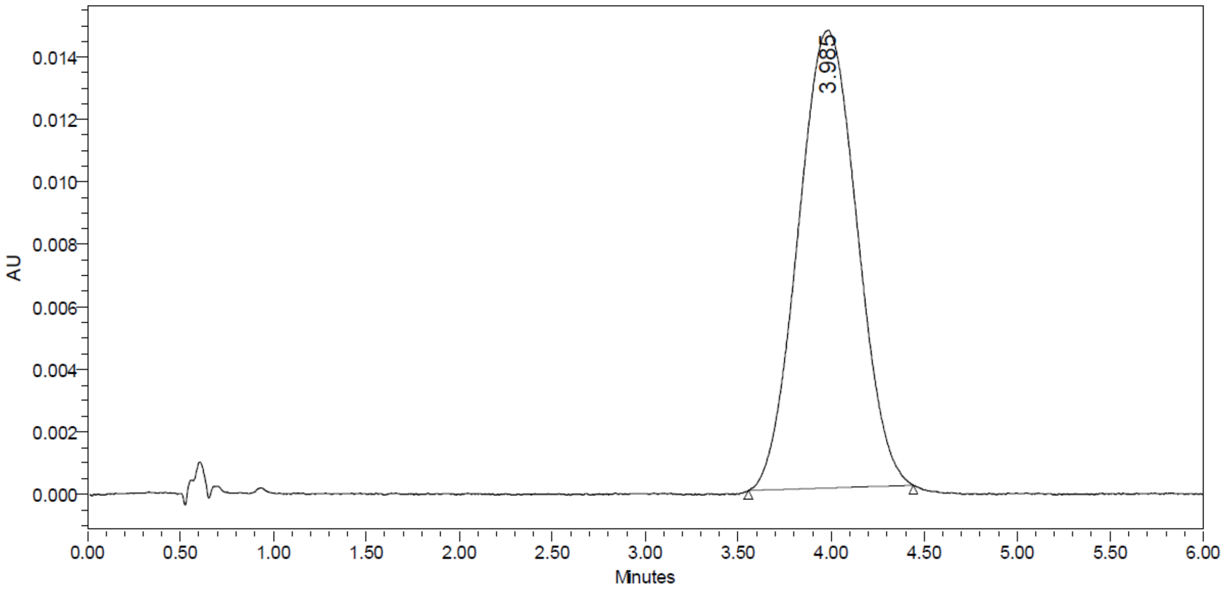


| | RT | Area | % Area | Height |
|---|---|---|---|---|
| 1 | 3.985 | 322393 | 100.00 | 14650 |

#### Compound 5b: $^{1}H$ and $^{13}C$ NMR in $CDCl_3$

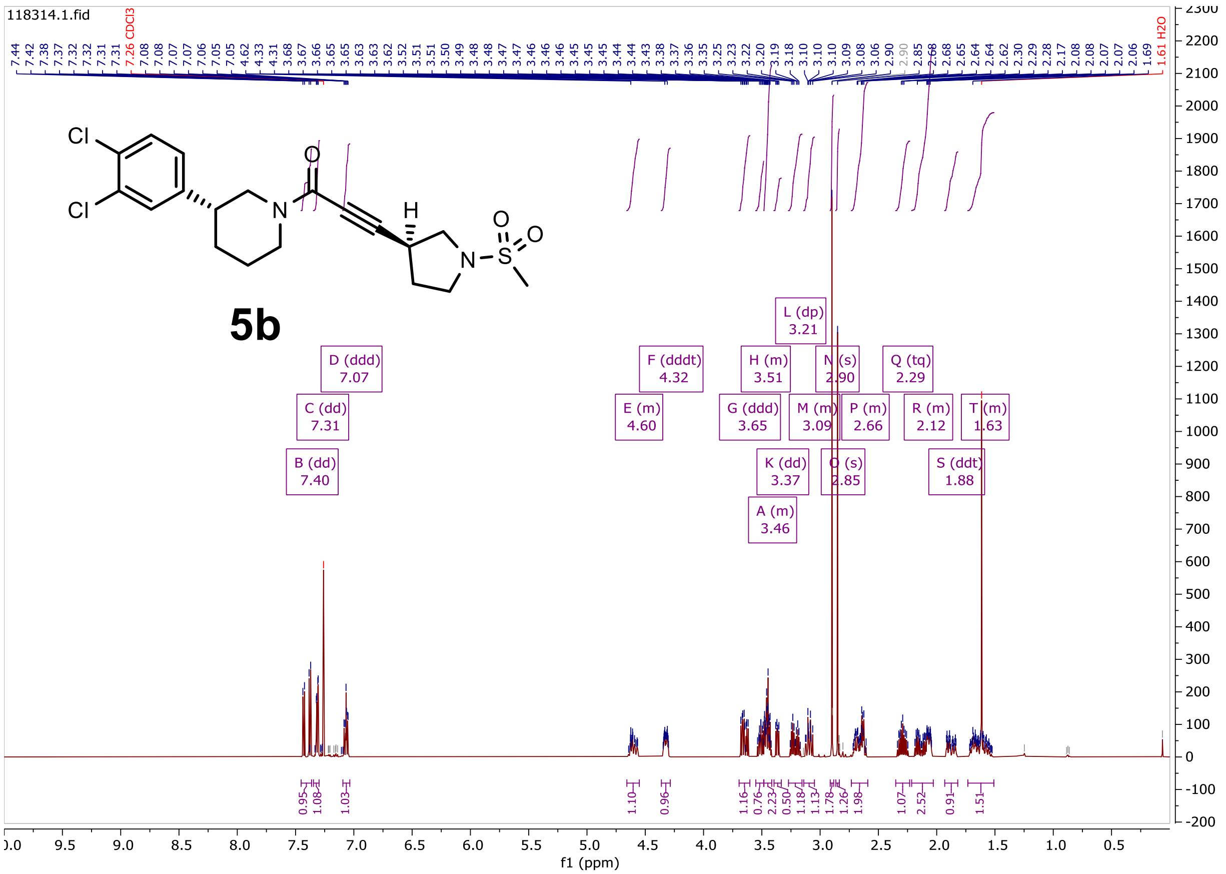

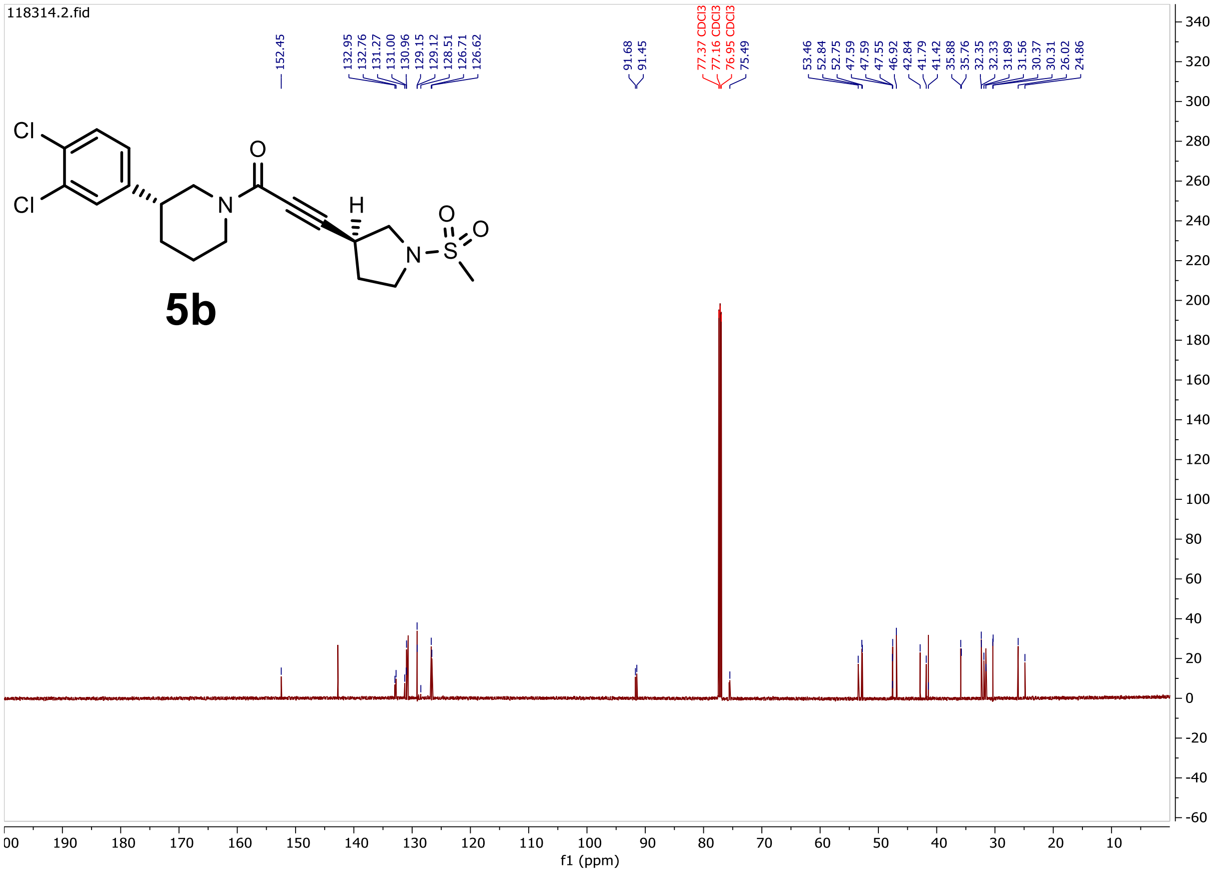


SFC of compound **5b**:

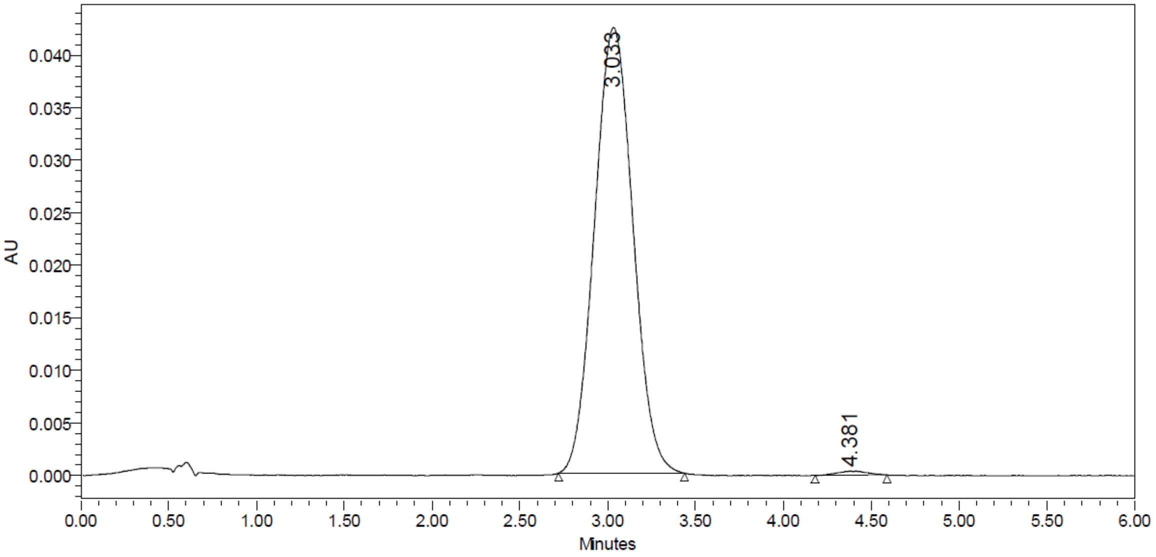


| | RT | Area | % Area | Height |
|---|---|---|---|---|
| 1 | 3.033 | 646597 | 99.32 | 42413 |
| 2 | 4.381 | 4453 | 0.68 | 370 |

### Compound 5c: $^{1}H$ and $^{13}C$ NMR in $CDCl_3$

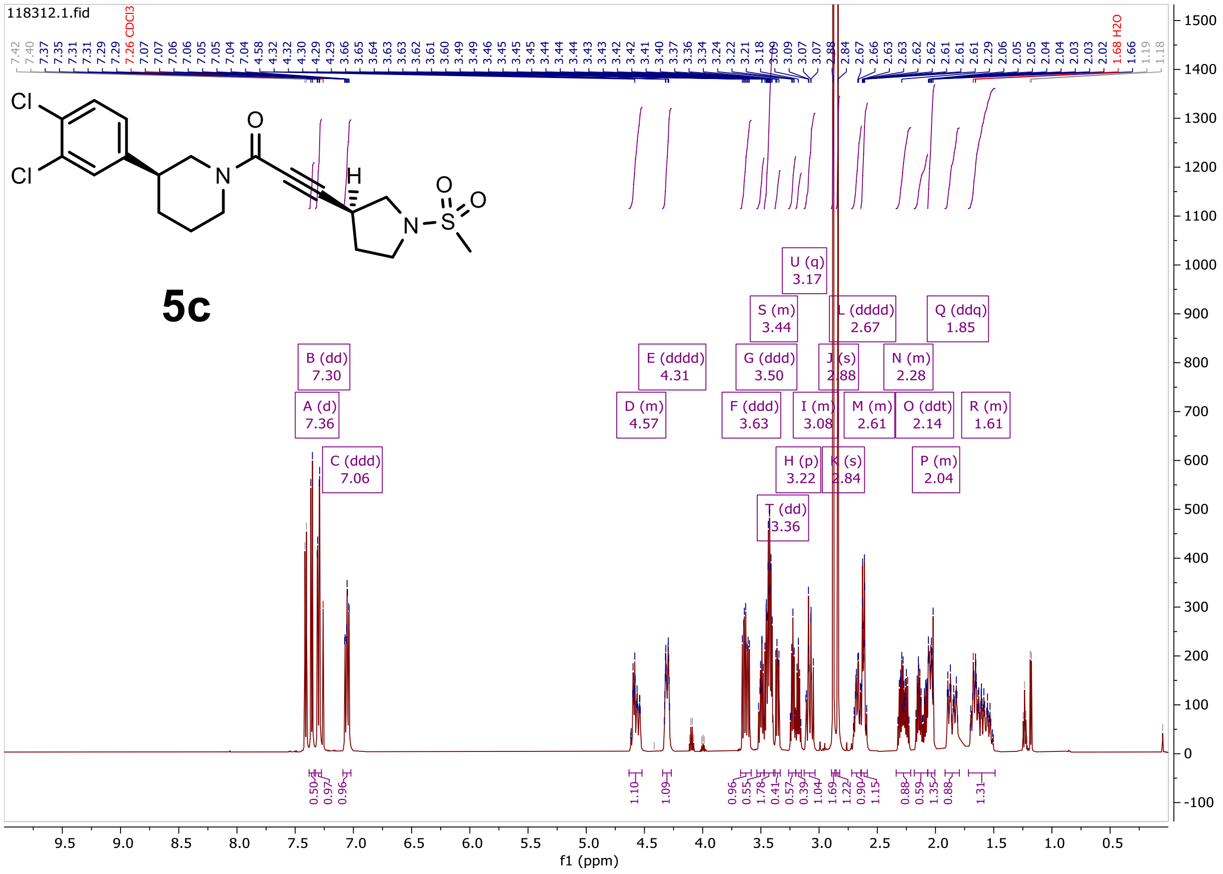

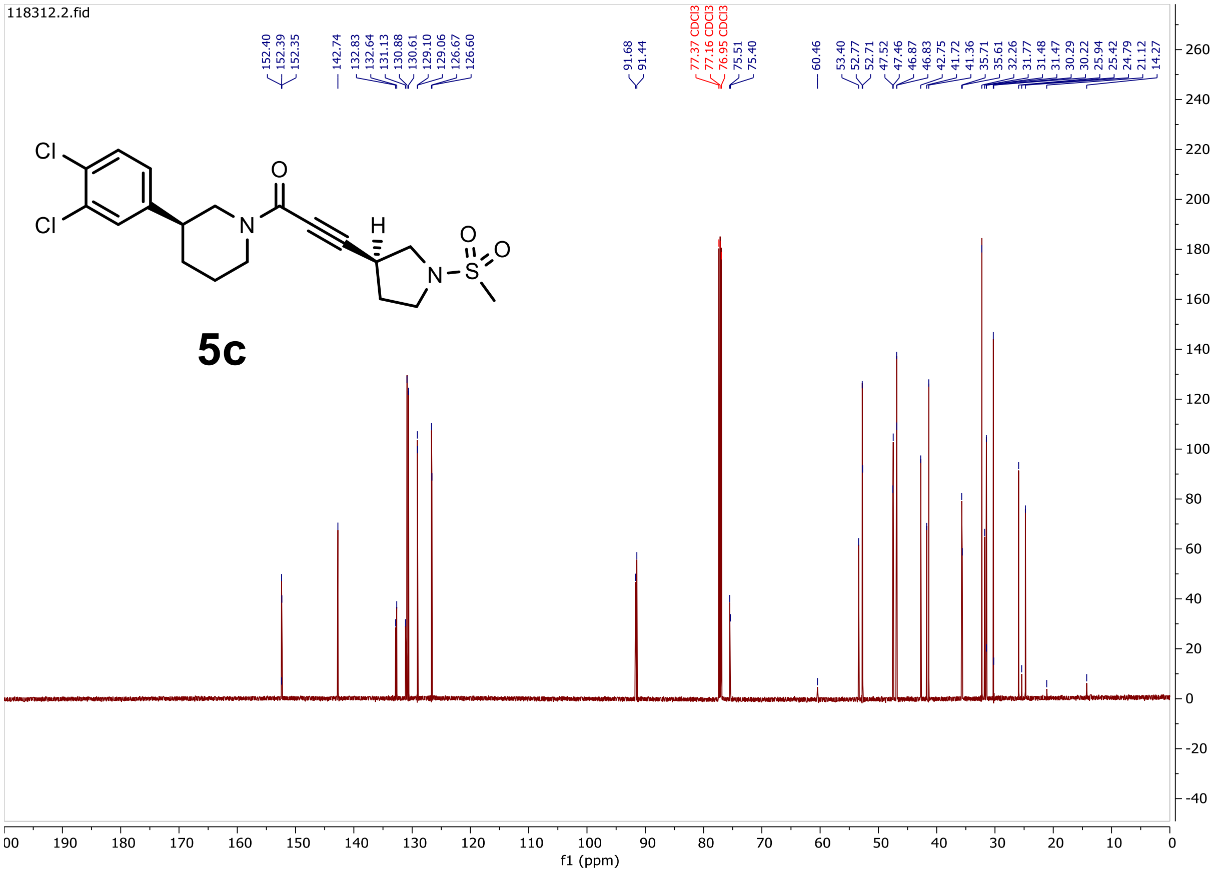


SFC of compound **5c**:

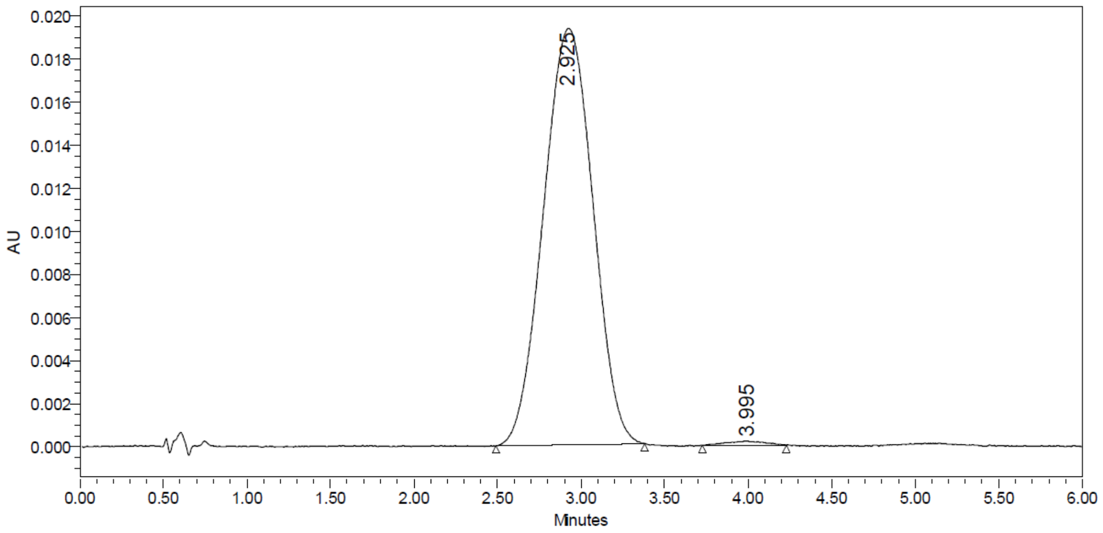


| | RT | Area | % Area | Height |
|---|---|---|---|---|
| 1 | 2.925 | 398789 | 99.26 | 19336 |
| 2 | 3.995 | 2979 | 0.74 | 194 |

### Compound 5d: [1]H and [13]C NMR in DMSO-$d_6$

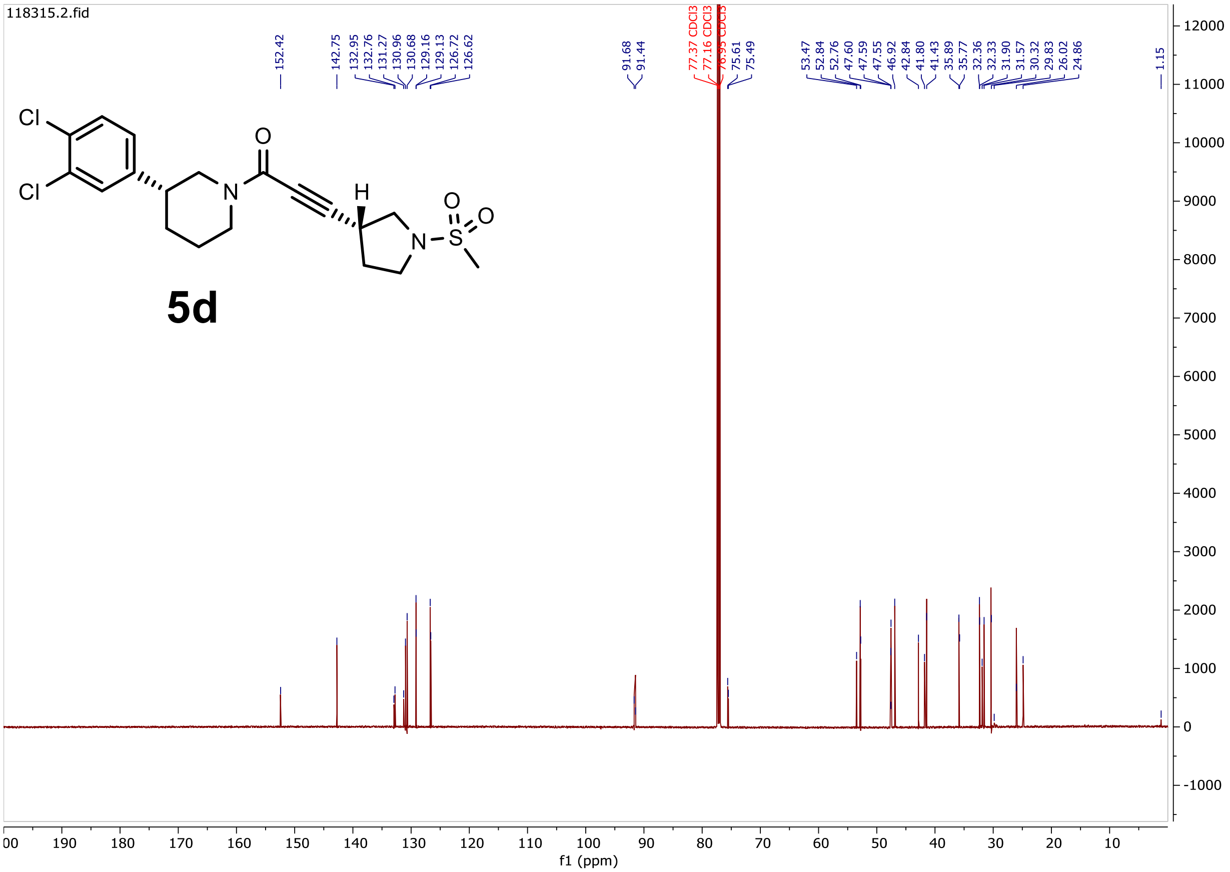

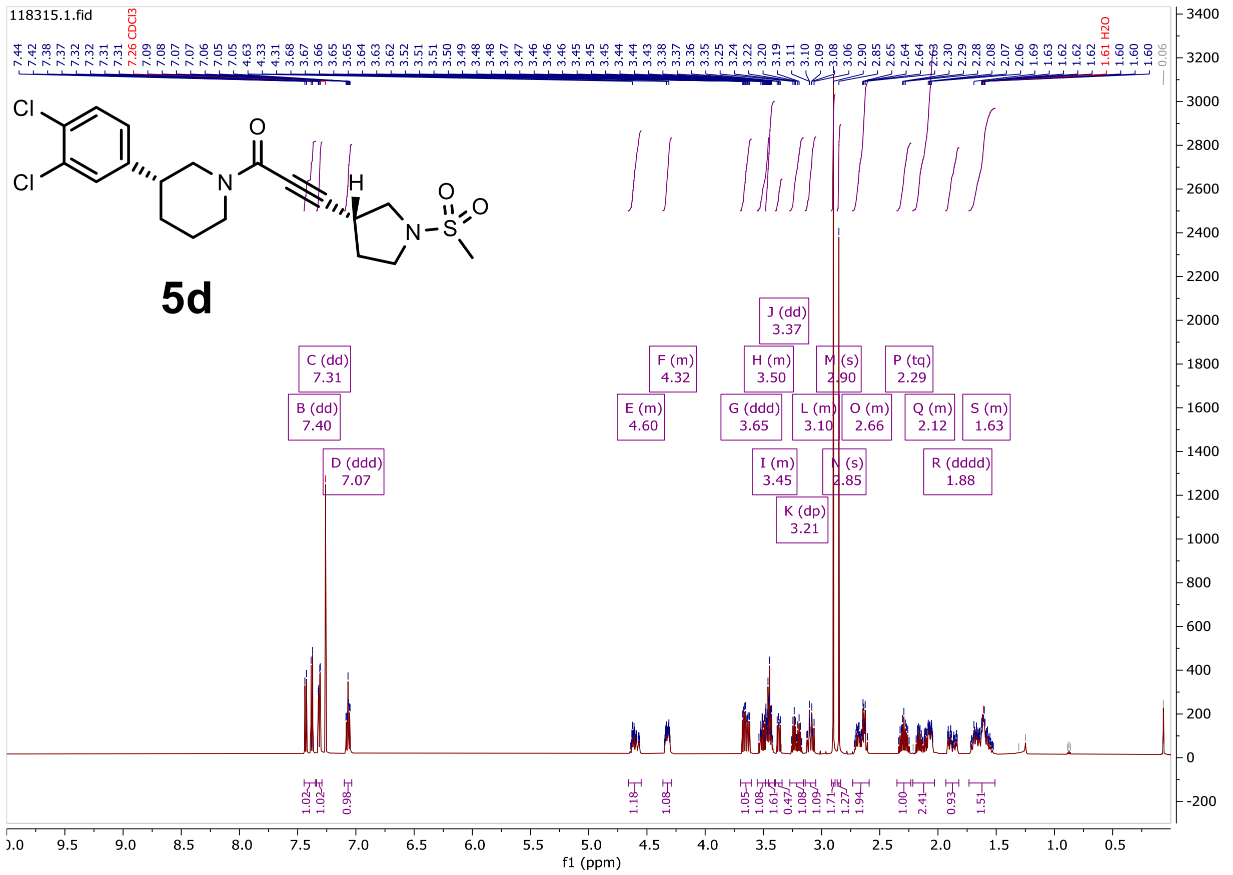

SFC of compound **5d**:

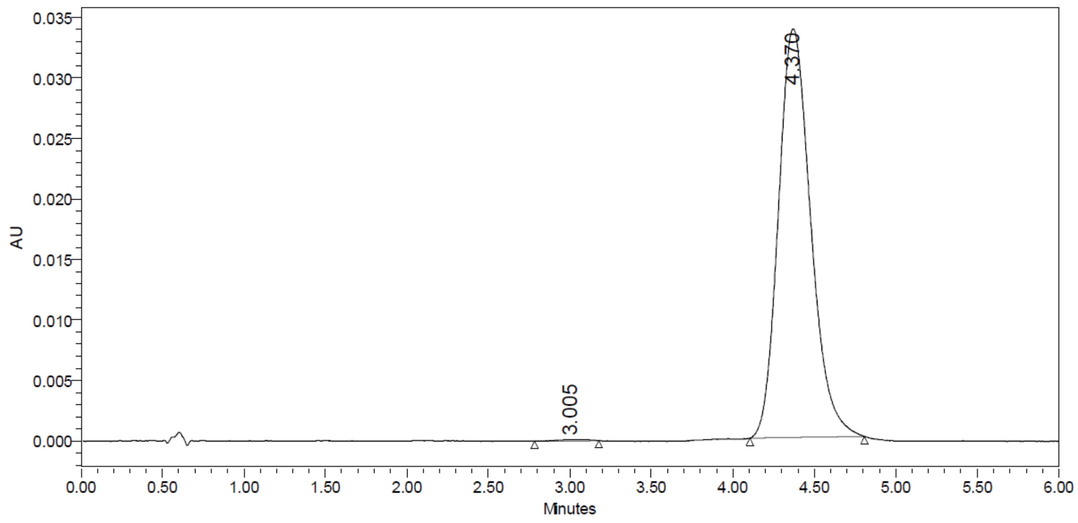

| | RT | Area | % Area | Height |
|---|---|---|---|---|
| 1 | 3.005 | 1550 | 0.33 | 124 |
| 2 | 4.370 | 465526 | 99.67 | 33763 |

#### Compound 6: $^{1}H$ and $^{13}C$ NMR in DMSO-$d_6$

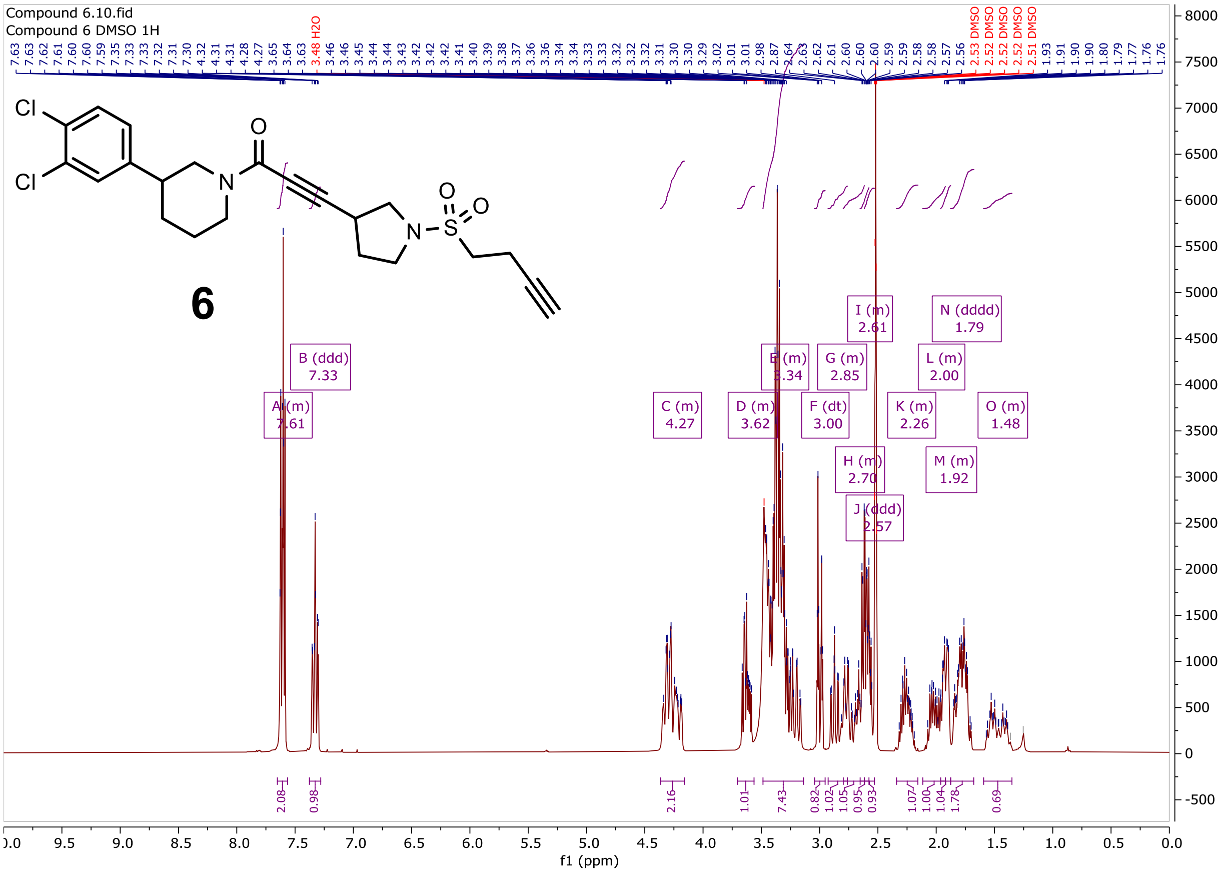

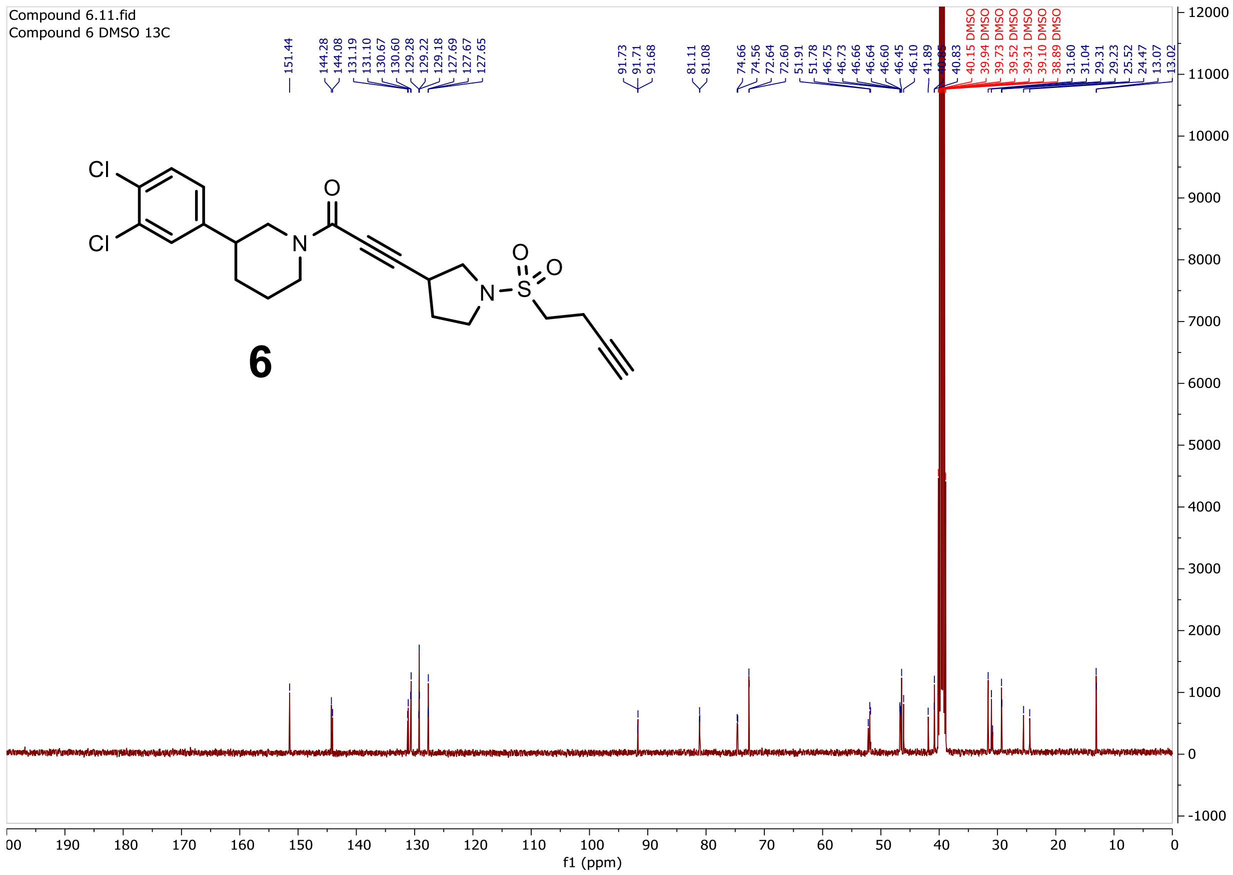

Compound 6.11.fid
Compound 6 DMSO 13C
151.44
144.28
144.08
131.19
131.10
130.67
130.60
129.28
129.22
129.18
127.69
127.67
127.65
91.73
91.71
91.68
81.11
81.08
74.66
74.56
72.64
72.60
51.91
51.78
46.75
46.73
46.66
46.64
46.60
46.45
46.10
41.89
40.83
40.15 DMSO
39.94 DMSO
39.73 DMSO
39.52 DMSO
39.31 DMSO
39.10 DMSO
38.89 DMSO
31.60
31.04
29.31
29.23
25.52
24.47
13.07
13.02
Cl
Cl
N
O
N
S
O
O
6
f1 (ppm)
