## Supplementary Figures and Biology Methods for "Selective inhibitors of JAK1 targeting a subtype-restricted allosteric cysteine"

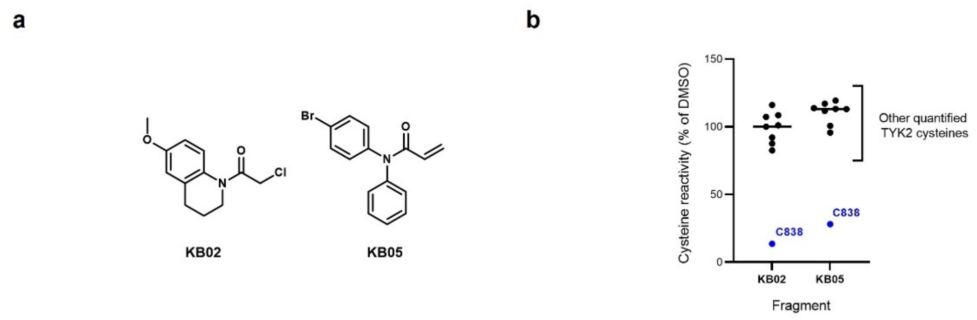


**Extended Data Fig. 1. Discovery of a ligandable cysteine in the JAK1/TYK2 pseudokinase domain. a**, Chemical structures of broadly reactive electrophilic fragments KB02 and KB05 evaluated previously for covalent reactivity with cysteines in the human T-cell proteome (Vinogradova, E. V. *et al*, *Cell* **182**, 1009–1026 e29 (2020), **b**, Relative MS3 signal intensity values for all quantified IA-DTB-labeled, cysteine-containing peptides in TYK2 in KB02- or KB05-treated T cells compared to DMSO-treated T cells. The KB02- and KB05-liganded cysteine in TYK2 (C838) is highlighted in blue. Horizontal black bars indicate the median signal intensity for all other quantified TYK2 cysteines. Data are mean values of two (KB02) or three (KB05) independent replicates analyzed in a single MS-ABPP experiment.

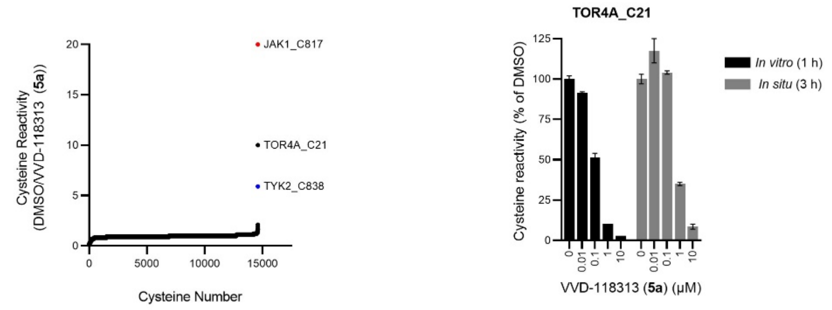


**Extended Data Fig. 2. Cysteine reactivity profile for VVD-118313 in PBMC lysates.** *Left*, Global cysteine reactivity profile for VVD-118313 (1 µM, 1 h, *in vitro*) in primary human PBMC lysates (2 mg/mL proteome). Data represent mean ratio values (DMSO/VVD-118313) for IA-DTB-labeled, cysteine-containing peptides quantified from two replicate cell treatment experiments analyzed in a single MS-ABPP experiment. Ratio values for JAK1_C817 (red) and TYK2_C838 (blue) are highlighted. Quantified cysteines with ratios ≥ 4 (≥ 75% engagement) are marked. *Right*, Concentration-dependent reactivity profile for VVD-118313 reactivity with TOR4A_C21 in human PBMCs (0.01-10 µM, 3 h, *in situ*) or PBMC lysates (0.01-10 µM, 1 h, *in vitro*) Data are mean values from VVD-118313-treated samples shown as a percentage of DMSO-treated samples from two replicate cell treatment experiments analyzed in a single MS-ABPP experiment.

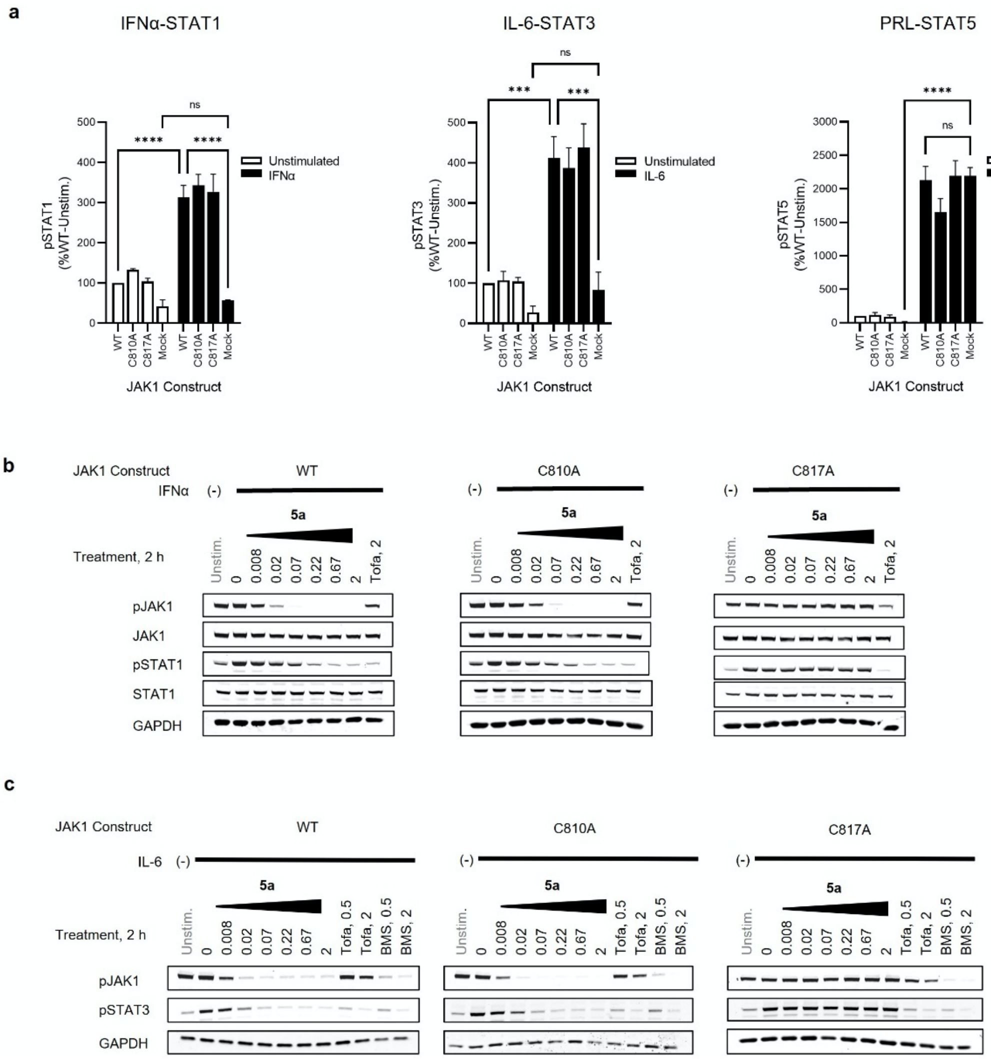


**Extended Data Fig. 3 Characterization of VVD-118313 inhibitory activity against JAK1 in 22Rv1 cells. a,** Quantification of western blotting data measuring cytokine-stimulated STAT phosphorylation in 22Rv1 cells expressing WT-, C810A-, or C817A-JAK1 variants compared to mock-transfected 22Rv1 cells (see **Fig. 3b** for representative western blots). Cells were treated with IFNα (100 ng/mL, 30 min), IL-6 (50 ng/mL, 30 min) or prolactin (PRL, 15 ng/mL, 15 min) after which the indicated phosphorylated STATs (pSTATs) were measured. Signal intensities were normalized to unstimulated 22Rv1 cells expressing WT-JAK1. Data are mean values ± S.E.M. from three independent experiments. Significance was determined by two-way ANOVA with Tukey's post-hoc test and reported for select comparisons. ****P<0.0001, ***P<0.001, ns – non-significant. IFNα and IL-6-stimulated STAT1/3 phosphorylation was significantly enhanced by expression of any of the three JAK1 variants (P<0.0001), while prolactin-stimulated STAT5 phosphorylation was unaffected by JAK1 expression. **b**, **c**, Representative western blots showing concentration-dependent effects of VVD-118313 (**5a**) on IFNα-stimulated STAT1 phosphorylation (**b**) and IL-6-stimulated STAT3 phosphorylation (**c**) in 22Rv1 cells expressing WT-, C810A-, or C817A-JAK1 variants.

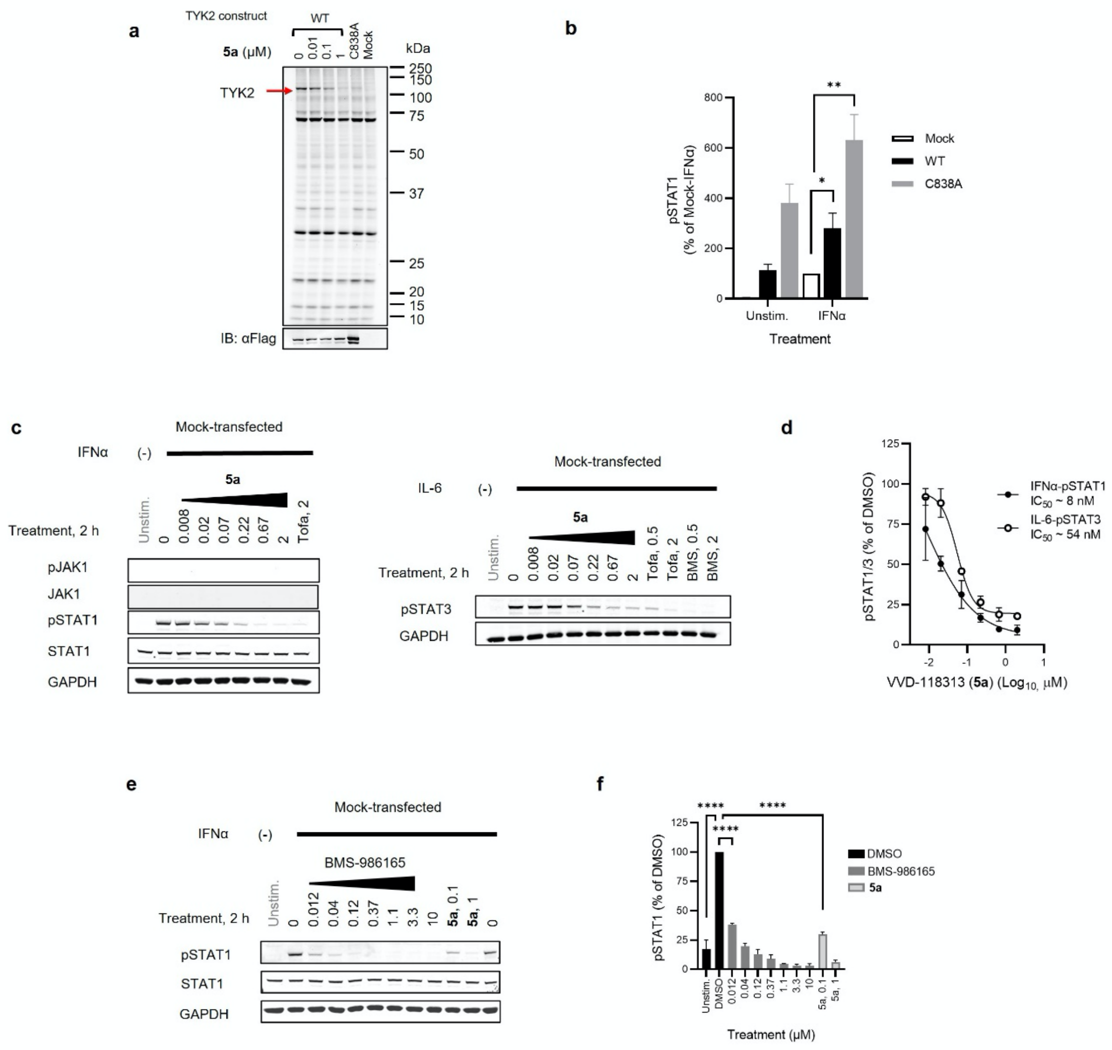


**Extended Data Fig. 4. Engagement of TYK2_C838 and inhibition of TYK2-dependent signaling in 22Rv1 cells. a,** gel-ABPP experiment showing labeling of recombinant WT-TYK2, but not C838A-TYK2, expressed in 22Rv1 cells by alkyne probe **6** (0.1 μM, 2 h*, in situ*). The labeling of WT-TYK2 was blocked by pretreatment with VVD-118313 (**5a**) (0.01-1 μM, 2 h, *in situ*). We noted that the C838A-TYK2 mutant consistently expressed at higher levels than WT-TYK2, as revealed by the anti-TYK2 immunoblot (bottom). Data are from a single experiment representative of two independent experiments. **b,** Quantification of IFNα-stimulated STAT1 phosphorylation in TYK2 (WT or C838A)-transfected 22Rv1 cells compared to mock-transfected cells. Signal intensities were normalized to IFNα-treated (100 ng/mL, 30 min) mock-transfected cells. Data are mean values ± S.E.M. from four independent experiments. Significance was determined using a two-tailed Student's *t*-test. *P<0.05, **P<0.01. **c-f**, Western blots (**c, e**) and quantification (**d, f**) of the effect of VVD-118313 (**5a**), tofacitinib (Tofa), and BMS-986165 (BMS) on IFNα-stimulated STAT1 and IL-6-stimulated STAT3 phosphorylation in mock-transfected 22Rv1 cells, which lack JAK1. Unstim, unstimulated controls. **d, f**, Quantification of pSTAT signals shown as a percent of the stimulated DMSO-treated control cells for each assay. Data are mean values ± S.D. from two independent experiments (**c, d**), or mean values ± S.E.M. from three independent experiments (**e, f**). Significance was determined by one-way ANOVA with Tukey’s post-hoc test and reported for selected comparisons. ****P<0.0001.

a

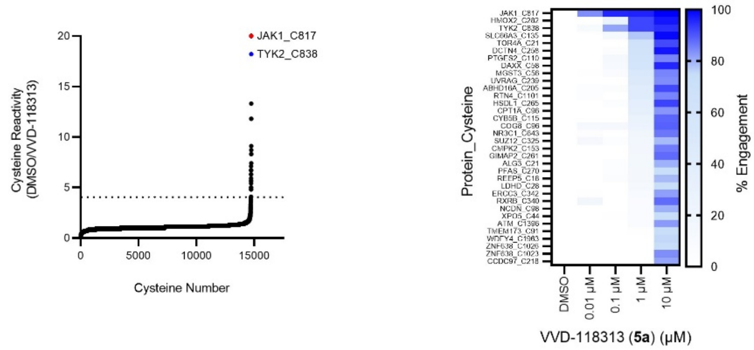


b

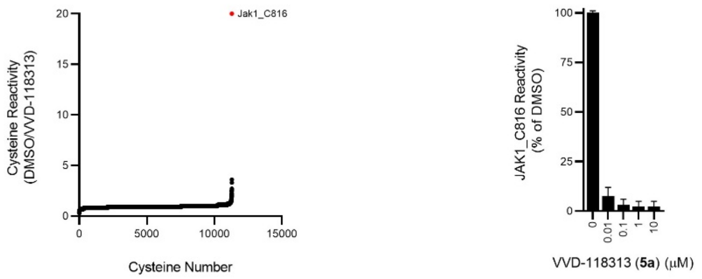


**Extended Data Fig. 5. Global cysteine reactivity profiles for VVD-118313 in primary human and mouse immune cells. a,** *Left,* Global cysteine reactivity profile for VVD-118313 (**5a**; 10 µM, 3 h, *in situ*) in primary human PBMCs. Reactivity values for JAK1_C817 (red) and TYK2_C838 (blue) are highlighted, and dashed horizontal line marks boundary for > 75% engagement by VVD-118313 at 10 µM. *Right*, heat map showing the reactivity profiles for cysteines in PBMCs treated with the indicated concentrations of VVD-118313**.** Only cysteines that were engaged >75% by VVD-118313 at 10 µM are shown. Data in both panels represent mean ratio values (DMSO/VVD-118313) for IA-DTB-labeled, cysteine-containing peptides quantified from two replicate cell treatment experiments analyzed in a single MS-ABPP experiment. **b**, *Left,* Global cysteine reactivity profile for VVD-118313 (1 µM, 1 h, *in vitro*) in mouse splenocyte lysates. Jak1_C816 shown in red. *Right*, Reactivity of JAK1_C816 in mouse splenocyte lysates treated with the indicated concentrations of VVD-118313 (**5a**; 1 h). Data in both panels represent mean ratio values (DMSO/VVD-118313) for IA-DTB-labeled, cysteine-containing peptides quantified from two replicate cell treatment experiments analyzed in a single MS-ABPP experiment.

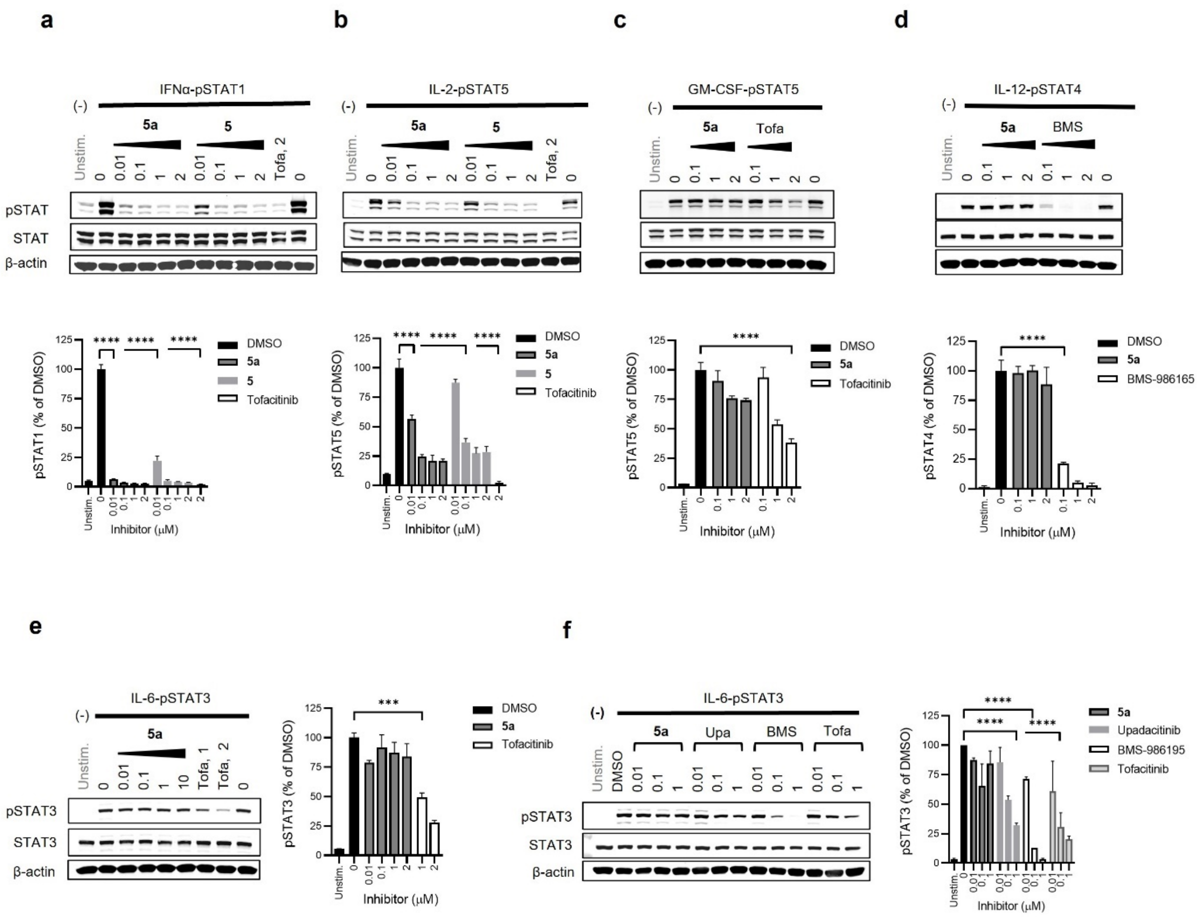


**Extended Data Fig. 6. Characterization of the inhibitory activity of VVD-118313 in mouse splenocytes**. **a-e**, *Top*, Western blots showing concentration-dependent effects of VVD-118313 (**5a**) and/or compound **5** on IFNα-stimulated STAT1 phosphorylation (**a**), IL-2-stimulated STAT5 phosphorylation (**b**), GM-CSF-stimulated STAT5 phosphorylation (**c**), IL-12-stimulated STAT4 phosphorylation (**d**), and IL-6-stimulated STAT3 phosphorylation (**e**) in mouse splenocytes. Tofacitinib (Tofa) and BMS-986165 were also tested where indicated. Splenocytes were treated with compounds at indicated concentration for 2 hours prior to stimulation with IFNα (100 ng/mL, 30 min), IL-6 (10 ng/mL, 30 min), IL-2 (20 U/mL, 15 min), GM-CSF (10 mg/mL, 15 min) or IL-12 (12.5 ng/mL, 15 min). *Bottom*, Quantification of pSTAT signals. Signal intensities were normalized relative to stimulated DMSO-treated controls in each assay. Data are mean values ± S.E.M. from three-four independent experiments. Significance determined by one-way-ANOVA with Dunnett's post-hoc test. P-values are shown for the lowest concentration of compound to inhibit S.I. ≥50%. ****P<0.0001, ***P<0.001, **P<0.01, *P<0.05. **f**, *Left*, Western blot showing effects of a panel of JAK inhibitors on IL-6-stimulated STAT3 phosphorylation in mouse splenocytes. *Right*, quantification of pSTAT3 signals performed and analyzed as described in **a-e**. Data are mean values ± S.D. from two independent experiments.

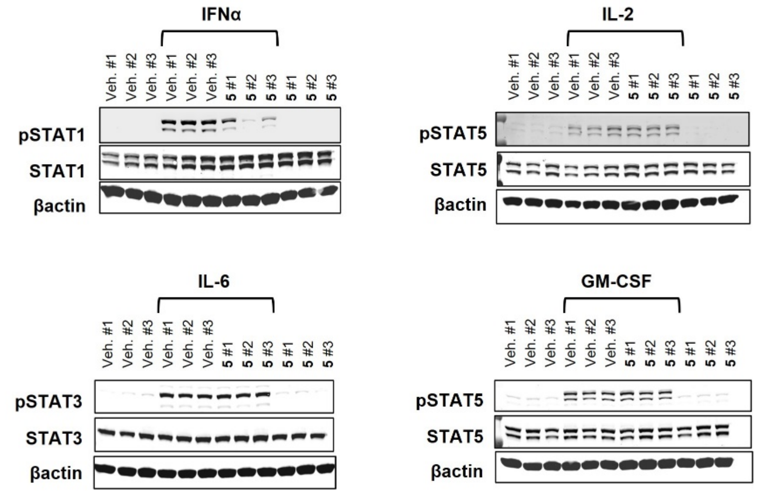


**Extended Data Fig. 7. VVD-118313 inhibits JAK1-dependent signaling *ex vivo*.** Western blots containing the results quantified in **Fig. 4i**, which represents *ex vivo* cytokine-stimulated STAT phosphorylation assays performed in splenocytes from mice treated with vehicle or compound **5** (25 mg/kg, 2 x 4 h). Splenocytes were stimulated with IFNα (1000 U/mL, 30 min), IL-2 (20 U/mL, 15 min), IL-6 (10 ng/mL, 30 min) or GM-CSF (10 ng/mL) prior to analysis of indicated STAT phosphorylation signals. #1-3 correspond to three individual mice in each treatment groups.

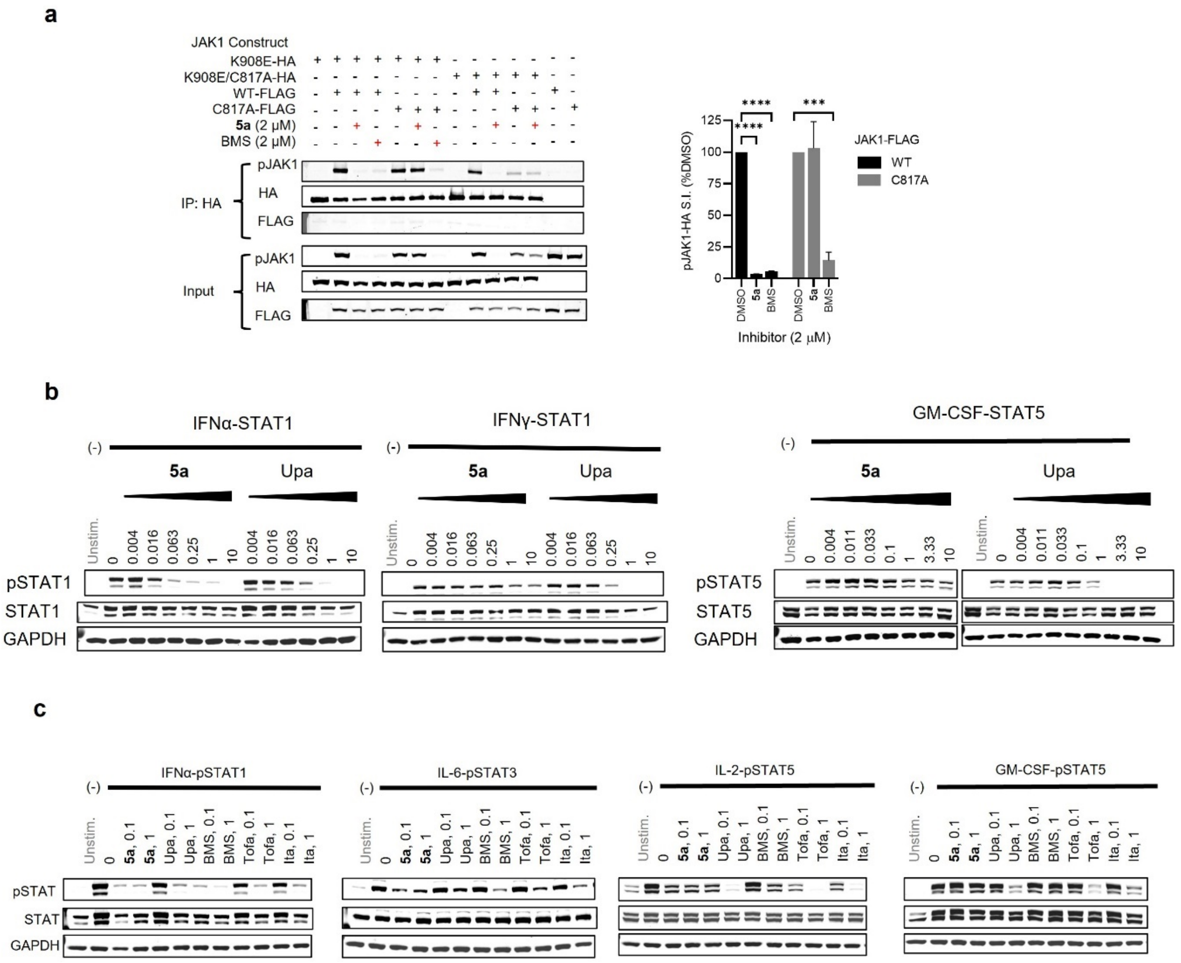


**Extended Data Figure 8. Mechanistic properties and distinct activity profile of allosteric JAK1 inhibitors. a**, *Left*, Western blots measuring effects of VVD-118313 (**5a**) and BMS-986165 (BMS) (2 μM, 2 h) on JAK1 phosphorylation (pJAK1) from anti-HA immunoprecipitations (IPs) of HA-tagged kinase dead (K908E) JAK1 (WT or C817A mutant) expressed in 22Rv1 cells alongside catalytically active FLAG-tagged JAK1 (WT or C817A mutant). *Right*, quantification of pJAK1 data, where pJAK1 signals in HA-immunoprecipitation eluent were normalized as a % of the respective DMSO-treated controls. Data are mean values ± S.E.M. from three independent experiments. Significance determined by two-way ANOVA with Dunnett's post-hoc test. ****P<0.0001, ***P<0.001. **b**, Western blots related to **Fig. 5g** showing concentration-dependent effects of VVD-118313 (**5a**) and upadacitinib (Upa) on the indicated cytokine (IFNα, IFNγ or GM-CSF)-stimulated STAT (STAT1/5) phosphorylation pathways. **c**, Western blots related to **Fig. 5i** showing effects of the indicated JAK inhibitors on the indicated cytokine-STAT phosphorylation pathways. Human PBMCs were treated with the compounds – VVD-118313 (**5a**), upadactinib (Upa), BMS-986165 (BMS), tofacitinib (Tofa. and itacitinib (Ita) – at the indicated concentrations (μM) for 2 h and then stimulated with IFNα (100 ng/mL, 309 min), IL-6 (25 ng/mL, 30 min), IL-2 (20 U/mL< 15 min) or GM-CSF (0.5 ng/mL, 15 minutes).

**Extended Data Table 1. In vivo pharmacokinetic properties of compound 5.** CD-1 mice (ICR) were dosed by intravenous (i.v.) or subcutaneous (s.c.) administration with compound **5**, prepared in 5% DMSO/95%(20%HPβCD). PK parameters were calculated by non-compartmental analysis of the plasma concentration–time profiles and reported as the mean of n = 3 mice/group.

| ID | Route | Dose (mg/kg) | CL (mL/min/kg) | $T_{1/2}$ (h) | F% | $C_0$ or $C_{max}$ (ng/mL) | $AUC_{(0\text{-}last)}$ (h·ng/mL) |
|---|---|---|---|---|---|---|---|
| **5** | i.v. | 1 | 112 | 0.153 | NA | 1608 | 149 |
| | s.c. | 25 | NA | 0.360 | 78.3 | 3377 | 2926 |

### Experimental Section

#### Antibodies, cytokines and inhibitors

Phospho-JAK1 (Tyr1034/1035) (D7N4Z) (74129), phospho-TYK2 (Tyr1054/1055) (D7T8A) (68790), TYK2 (D4I5T) (14193), phospho-STAT1 (Tyr701) (58D6) (9167), phospho-STAT3 (Tyr705) (D3A7) XP (9145), phospho-STAT4 (Tyr693) (D2E4) (4134), phospho-STAT5 (Tyr694) (C11C5) (9359), STAT1 (D1K9Y) (14994), STAT3 (79D7) (4904), STAT4 (C46B10) (2653), STAT5 (D2O6Y) (94205), HA-Tag (C29F4) (3724) and β-actin (13E5) (4970) rabbit monoclonal antibodies were obtained from Cell Signaling Technologies. Anti-JAK1 antibody (610231, BD Transduction Laboratories), anti-Flag M2 antibody (F1804, Sigma) and anti-GAPDH (sc-47724, Santa Cruz) mouse antibodies were sourced as indicated. Secondary IRDye antibodies for western blot were purchased from Li-Cor under product numbers: 926-32210, 926-32213, 926-68020 and 926-68023.

Commercial JAK inhibitors BMS-986165 (HY-117287, MedChemExpress), tofacitinib (S2789, Selleckchem), upadacitinib (NC1927829, Fisher Scientific) and itacitinib (INCB39110) (501948171, Selleckchem) were obtained from commercial vendors.

Recombinant human IFNα (11101-2) and mouse IFNα (12100-1) were purchased from PBL Assay Sciences. Recombinant IL-2 for human and mouse experiments was purchased from Hoffman-La Roche (TECIN Teceleukin, Bulk Ro 23-6019). All other cytokines were purchased from R&D Biosystems under the following catalogue numbers: recombinant human IFNγ (285-IF-100/CF), IL-6 (206-IL-010/CF), IL-12 (219-IL/CF), GM-CSF (7954-GM-010/CF), prolactin (682-PL-050), and recombinant mouse IL-6 (406-ML-005/CF), IL-12 (419-ML-010/CF), GM-CSF (415-ML-005/CF).

All cell lysates for western blotting were prepared using mPER mammalian protein extraction reagent (78501, ThermoFisher), unless otherwise noted. All lysis buffers were supplemented with cOmplete™ EDTA-free Protease Inhibitor Cocktail tablets (11873580001, Roche) and PhosSTOP phosphatase inhibitor cocktail tablets (4906837001, Roche).

#### DNA constructs and transfection

pCMV6-JAK1-Myc-Flag (RC213878) and pCMV6-TYK2-Myc-Flag (RC204351) vectors were obtained from Origene. Mutant JAK1 (C810A, C817A, K908E, K908E/C817A) and TYK2 (C838A) constructs were generated by site-directed mutagenesis, and HA-tagged constructions were generated by epitope-tag insertion using the Q5 Site-Directed Mutagenesis kit (New England BioLabs, E0552S). All constructs were confirmed by Sanger sequencing (Genewiz). Transfections were performed using PEI-MAX

#### Cell lines, primary cells, mice

22Rv1 human prostate cancer cells were purchased from the American Type Culture Collection (ATCC, #CRL-2505) and cultured at 37 °C and 5% $CO_2$. in complete RPMI-1640 (supplemented with ʟ-glutamine (2 mM), penicillin (100 U $ml^{-1}$), streptomycin (100 µg $ml^{-1}$) and 10% fetal bovine serum (FBS)). Cells were passaged every 3 days using trypsin and seeded into new 6-12 well tissue culture plates at the stated densities for assays. Compound treatments were performed in serum free RPMI.

Human peripheral blood mononuclear cells (PBMCs) were obtained from healthy donors (aged 18-50 yrs) recruited through the Scripps Normal Blood Donor service and used according to protocols approved by The Scripps Research Institute Institutional Review Board (protocol #IRB-187252). PBMCs were isolated from heparinized blood by Ficoll gradient (Lymphoprep, #7861, Stemcell Technologies) and red blood cells were lysed with 1x Red Blood Cell Lysis buffer (#00-4300-54, eBiosciences). Purified PBMCs were then washed with PBS (1x) and RPMI (1x), before being resuspended in serum free RPMI (supplemented with L-glutamine (2 mM), penicillin (100 U ml$^{-1}$), and streptomycin (100 µg ml$^{-1}$)) for phospho-STAT assays. For IL-12 assays, PBMC-derived T-blasts were generated by stimulating freshly isolated PBMCs with phytohaemagglutinin (PHA-P, 10 µg ml$^{-1}$) in complete RPMI for 72 h, followed by IL-2 (100 U ml$^{-1}$) in fresh RPMI for 24 hours. Cells were then washed and rested overnight in serum free RPMI prior to performing IL-12-phospho-STAT4 assays.

All animals used in this study were adult (8-12 weeks) C57BL/6 mice, except for pharmacokinetic studies, where male CD-1 (ICR) mice were used. Mice were maintained in pathogen-free conditions and handled in accordance with requirements of the National Institutes of Health and the Institutional Animal Care and Use Committee at The Scripps Research Institute. Splenocytes were isolated from fresh spleens by passing through a 70 µm sieve (#130-110-916, Miltenyi Biotec Inc). Debris and red blood cells were removed by lysis in 1x RBC Lysis buffer (#00-4300-54, eBiosciences), followed by sequential washes with PBS (x2), and serum free RPMI (1x). Splenocytes were then resuspended at 6 x $10^6$ cells ml$^{-1}$ in serum free RPMI for phospho-STAT assays.

**Gel-based ABPP fluorescence**

22Rv1 cells were seeded (2.5 million/10 cm plate) in complete RPMI media (10% FBS) 24 hours prior to PEI transfection with JAK1/TYK2 constructs (5 µg/plate). 24 hours after transfection, cells were serum starved overnight, then treated with DMSO or **5a** (0.01, 0.1 or 1 µM) for 2 hours at 37 °C, followed by alkyne **6** (0.1 µM) for 2 hours at 37 °C. After treatments, cells were harvested by scraping in cold PBS, centrifuged (1,400 g, 2 min), pellets were washed with PBS (1X) and then frozen at -80 °C until use.

To label alkyne **6** with a rhodamine reporter tag, cells were thawed on ice, resuspended in PBS (350 µl) PBS containing complete protease inhibitors, and lysed by sonication (2 x 8 pulses). The protein concentration of whole cell lysates was normalized to 1.2 mg ml$^{-1}$ and 50 µl was used for copper-catalyzed azide–alkyne cycloaddition (CuAAC) reactions. The CuAAC reaction mixture was prepared using a 1:1:1:3 ratio of rhodamine azide (1.25 mM in DMSO; 25 µM final), 50 mM $CuSO_4$ (aq., 1 mM final), 50 mM TCEP (aq., 1 mM final) and 1.7 mM tris(benzyltriazolylmethyl)amine (4:1 *t*-BuOH/DMSO; 100 µM final). Lysates were treated with 6 µl of Rh-CuAAC reaction mixtures, vortexed and incubated for 1 hour at room temperature (vortexing 1-2 x). Reactions were then quenched with 18 µl 4x Laemmli sample buffer and either frozen at -20 °C or immediately resolved on 10% Tris-glycine polyacrylamide gels, loading 20 µg protein per lane. Rhodamine fluorescence was detected using a BioRad Imager flatbed scanner and images were processed in manufacturer software.

**Phospho-STAT western Blot assays in 22Rv1 cells**

22Rv1 cells (0.6 x $10^6$ cells/well) were seeded in 12 well plates in complete RPMI (10% FBS). After 24 hours, cells were transfected with JAK1/TYK2 constructs (500-1500 ng DNA/well) using PEI-max (24765-1, Polysciences) diluted in OptiMEM (31985062, Gibco) and incubated at 37 °C for 24 hours. Media was then replaced and cells were serum-starved overnight prior to compound treatments. Cells were treated with DMSO or JAK inhibitors in serum free RPMI for 2 h at 37 °C, then stimulated with IFNα (100 ng ml$^{-1}$, 30 min), IFNγ (1000 U ml$^{-1}$, 30 min), IL-6 (50 ng ml$^{-1}$, 30 min) or prolactin (15 ng ml$^{-1}$, 15 min). Media was removed, cells were washed with PBS (1x) and lysed in mPER buffer (150 µL), supplemented with protease and phosphatase inhibitors for 10-20 minutes at room temperature. Lysates were collected, cleared by centrifugation (5 min, 16,000 g, 4 °C) and supernatants were combined with 4x Laemmli sample buffer for western blotting.

**Phospho-STAT western blot assays in primary immune cells**
Cytokines were used at a concentration that generated ~ 50–90% of the maximal cytokine-induced signal in each assay ($EC_{50-90}$). Freshly isolated PBMCs (or PHA/IL-2 activated T-blasts for IL-12 assays) were resuspended in serum free RPMI (6 million cells ml$^{-1}$) and treated with DMSO or JAK inhibitors for 2 hours at 37 °C, then stimulated with cytokines as follows: IFNα (100 ng ml$^{-1}$, 30 min), IFNγ (1000 U ml$^{-1}$, 30 min), IL-6 (25 ng ml$^{-1}$, 30 min), IL-2 (20 U ml$^{-1}$, 15 min), GM-CSF (0.5 ng ml$^{-1}$, 15 min) or IL-12 (12.5 ng ml$^{-1}$, 15 min). Cells were then pelleted by centrifugation (1.5 min, 16,000 g), media was removed, and cells were lysed in 90 µL mPER buffer, containing protease and phosphatase inhibitors for 15-20 minutes at room temperature. Lysates were cleared by centrifugation (5 min, 16,000 g, 4 °C) and supernatant were combined with 4x Laemmli sample buffer for western blotting.

Western blot assays with splenocytes were performed as described above, with the following cytokine concentrations: murine IFNα (1000 U ml$^{-1}$, 30 min), IL-6 (10 ng ml$^{-1}$, 30 min), IL-2 (20 U ml$^{-1}$, 15 min), GM-CSF (10 ng ml$^{-1}$, 15 min) or IL-12 (12.5 ng ml$^{-1}$, 15 min). Murine IL-12 assays were performed on freshly isolated splenocytes and did not require *in vitro* activation to prior to assays to stimulate receptor expression.

***In vivo* compound treatment**
Compound **5** was formulated at 2.5 mg ml$^{-1}$ in 5% DMSO/20% hydroxy-propyl-β-cyclodextrin for all *in vivo* experiments. Adult (8-12 weeks), age and sex matched mice (n = 3) were administered 2 doses of **5** (25 mg kg$^{-1}$) or 5% v/v DMSO vehicle by subcutaneous injection at 4 hour intervals. Mice were sacrificed 4 hours after the second dose according to approved protocols.

***Ex vivo* phospho-STAT assays**
Splenocytes from mice treated with compound **5** or DMSO vehicle were isolated as described above and seeded at 6 x $10^6$ cells ml$^{-1}$ in serum free RPMI for *ex vivo* stimulation with IFNα (1000 U ml$^{-1}$, 30 min), IL-6 (10 ng ml$^{-1}$, 30 min), IL-2 (20 U ml$^{-1}$, 15 min) or GM-CSF (10 ng ml$^{-1}$, 15 min). Cells were collected by centrifugation, lysed with mPER buffer supplemented with protease and phosphatase inhibitors (60 µL, r.t., 10-20 min), cleared (5 min, 16,000 g, 4 °C) and then supernatants were combined with 4x 1x Laemmli sample buffer to prepare western blotting samples.

**Western blotting**
Samples in 1x Laemmli sample buffer were boiled for 5-10 min at 95 °C, then resolved by electrophoresis on either 10% or 4-20% Novex WedgeWell Tris-Glycine mini-gels (XP00105BOX, XP04205BOX, Invitrogen), and transferred to nitrocellulose (45004011, Amersham). Membranes were blocked with 5% milk in Tris-buffered saline (20 mM Tris-HCl 7.6, 150 mM NaCl) supplemented with 0.1% Tween-20 (TBST) buffer for 1 hour at room temperature and then probed overnight at 4 °C with primary antibodies (1:1000) in 5% BSA/TBST. Membranes were washed 3 x 5 min with TBST, then probed for 1 hour at room temperature with IRDye secondary antibodies (1:10,000) in 5% BSA/TBST, washed a further 3 x 5 min with TBST

and then visualized using the Odyssey Infrared Imaging System (Li-Cor Biosciences). Densitometry was performed using Odyssey software, subtracting any background fluorescence and normalize channels as a percentage of the relevant control channels in each experiment.

**HTRF phospho-STAT assays**

Homogeneous time-resolved fluorescence (HTRF) assays to detect IFNα-stimulated STAT1 (Tyr701) phosphorylation and IL-6-stimulated STAT3 (Tyr705) phosphorylation in human PBMCs were performed using CisBio assay kits #63ADK026PEG and #62AT3PEG. Assays were performed using a modified version of the manufacturer's two-plate assay protocol, except that phospho-total protein lysis buffer #2 (64KL2FDF) was used for both pSTAT1 and pSTAT3 assays. In brief, stock compound plates were prepared by a 7-point serial dilution of compounds in 100% DMSO. Working plates were prepared immediately prior to assays by diluting stock plates serum free RPMI to give working solutions containing 0.3% v/v DMSO. Freshly isolated PBMCs in serum free RPMI (20 x $10^6$ cells/mL) were aliquoted (50 µL) into duplicate 96-well microplates (655098, Greiner Bio-one) and treated with diluted compounds (25 µL) for 2 hours at 37 °C (0.1% v/v final DMSO, 1 nM – 2 µM, 10 µM or 50 µM final compound concentration). Cells were then stimulated with diluted cytokines (25 µL) to give a final concentration of IFNα (100 ng $mL^{-1}$) or IL-6 (25 ng $mL^{-1}$). After 30 minutes at 37 °C, cells were lysed using 33 µL of phospho-total protein lysis buffer 2 supplemented with phospho-peptide blocking regent (4% v/v) and incubated at room temperature for 45 minutes. After pipetting to homogenize lysates, 16 µL was transferred to 384-well white microplates (784075, Greiner Bio-one) and treated with a premixed solution of anti-pSTAT d2:Eu Cryptate antibodies (1:1, 4 µL). Solutions were covered with a plate sealer, incubated overnight at room temperature, and then fluorescence intensity was measured using a BMG PHERAstar plate reader, with excitation set to 337 nm, and emission wavelengths at 665 nm and 620nm.

HTRF ratios were calculated as (Signal 665 nm/Signal 620 nm) x $10^4$. The basal HTRF ratio of unstimulated DMSO treated controls (2 per assay plate) was subtracted from cytokine treated samples and the data were normalized as a percentage of mean HTRF ratio of the cytokine stimulated-DMSO treated controls (5 per assay plate). At least two dose-response experiments were performed per compound. $IC_{50}$ values were estimated by fitting data to a 4-parameter logistic model in Graphpad Prism v 9.3.0, and the mean values ± S.D. of a minimum of two dose-response experiments, except where noted in Table 1.

***In vitro* kinase assay**

JAK1 biochemical activity assays were performed in 384-well microplates (784075, Greiner) using the Promega JAK1 Kinase Enzyme System (VA7207) and ADP-Glo Kinase Assay Kit (V6930). Assay conditions were optimized to quantify initial reaction rates and performed according to manufacturer instructions. Prior to assays, compounds were prepared as working solutions containing 5% v/v DMSO and added to assays to give a final concentration of 1% v/v DMSO. In brief, 30 ng of recombinant GST-JAK1 protein (residues 438-1154) was incubated with DMSO or VVD-118313 (**5a**) or tofacitinib in 1x kinase assay buffer (supplemented with 50 µM DTT) for 30 minutes at room temperature prior to the addition of 0.2 µg/mL IRS-1 peptide and 50 µM ATP. Kinase assays were incubated at room temperature for 60 min, then quenched with 5 µL ADP Glo reagent (40 min, r.t.), followed by 5 µL Kinase Detection Reagent (60 min, r.t.). ATP conversion was quantified by luminescence using a CLARIOstar (BMG Labtech) plate reader and data were normalized as a percent of the maximum (DMSO-treated) response. $IC_{50}$

values were calculated by fitting data to a 4-parameter logistic model in GraphPad Prism (v 9.3) software.

**Trans-phosphorylation assay**
22Rv1 cells (0.6 x $10^6$ cells/well) were seeded into 6-well TC plates in complete RPMI media 24 hours prior to transfection with HA-tagged kinase dead (K908E) JAK1 (WT or C817A) and Flag-tagged catalytically active JAK1 (WT or C817A) constructs (1:1 ratio, 3 µg total DNA). 24 hours after transfection, media was replaced and cells were serum starved overnight and then treated with DMSO, **5a**, BMS-986165 or tofacitinib (2 µM) for 2 hours at 37 °C. Cells were then washed with cold PBS and lysed on ice for 30 minutes in 400 µL immunoprecipitation (IP) buffer (50 mM Tris, pH 8, 150 mM NaCl, 1% NP-40, 1 mM EDTA), supplemented with protease and phosphatase inhibitors. Lysates were collected, cleared (10,000 g, 10 min, 4 °C), the protein concentration of supernatants was normalized to ~1.5 mg $ml^{-1}$, and ~ 480 µg was aliquoted to LoBind SafeLock 1.5 mL Eppendorf tubes for HA-tag immunoprecipitation. Remaining supernatant was combined with 4x Laemmli sample buffer and used to quantify IP input. Anti-HA agarose (26181, Pierce) was equilibrated with IP buffer (3 x washes, 1 min, 2000 g), and 7.5 µL/sample of packed resin was used to immunoprecipitate kinase dead JAK1 (2 h, 4 °C). Immunoprecipitated proteins were collected by pelleting resin (2 min, 2000 g, 4 °C), washing 3 x 500 µL with IP buffer, and then eluting proteins using 50 µL 2x Laemmli sample buffer. Samples were boiled (5-10 minutes) and then resolved on 10% or 4-20% Tris-glycine gels and transferred to nitrocellulose as described in “Western Blotting”.

The effect of C817A mutation on transphosphorylation efficiency was calculated by normalizing the phospho-JAK1 signal intensity of DMSO-treated samples to transfection conditions where both kinase dead (K908E) and active JAK1 constructs contained the native C817. As trans-phosphorylation efficiency was lower for C817A mutants, phospho-JAK1 signal intensity from compound-treated samples was normalized to the respective DMSO-treated sample for a given pair of JAK1 constructs. All trans-phosphorylation assays were performed in at least triplicate.

***In vivo* pharmacokinetic analysis**
Male CD-1 (ICR) mice (n = 3/group) were dosed by intravenous (i.v.) or subcutaneous (s.c.) administration with compound **5**, prepared in 5% DMSO/95%(20%HPβCD). Plasma samples for pharmacokinetic analysis were collected in a composite manner with three animals per time point. The plasma concentration of compound **5** was measured by a liquid chromatography tandem mass spectrometry method using positive electrospray ionization in multiple reaction monitoring mode. Plasma samples were extracted by protein precipitation using acetonitrile containing an internal standard. After vortexing and centrifugation, the supernatant was injected into an API6500 (AB SCIEX) liquid chromatography tandem mass spectrometry system for quantification. Pharmacokinetic parameters were calculated by non-compartmental analysis of the plasma concentration–time profiles and reported as mean values in Extended Data Table 1.

**Data Analysis and Statistics**
Quantitative data are expressed as mean ± S.E.M. in bar charts and as mean ± S.D. in dose-response curves used to calculate $IC_{50}$ values. In western blot quantification, pSTAT or pJAK S.I. was normalized as a percent of the cytokine-stimulated DMSO-treated sample on the same membrane, or the respective WT control in recombinant 22Rv1 experiments. Statistical analysis was performed in Graphpad Prism Software v9.2.0 using one-way ANOVA or two-way ANOVA with either Dunnett’s, Turkey’s or Šidák’s post-hoc test as noted in the corresponding figure legend. Significant P-values are only noted for the lowest concentration of compound to inhibit

STAT/JAK phosphorylation ≥50%, unless otherwise noted (*P<0.05, **P<0.01, ***P<0.001, ****P<0.0001). $IC_{50}$ values were estimated by fitting data using a 4-parameter logistic model (GraphPad Prism, v9.2.0).

**Proteome wide cysteine ligandability profiling by MS-ABPP**
*In vitro treatments:* PBMCs isolated from a single blood donor and flash frozen were thawed on ice and resuspended in PBS, containing protease inhibitors, and lysed by sonication on ice (3 x 10 pulses, 40% output). Protein concentration in whole cell lysate was estimated using a DC protein assay (Bio-Rad), normalized to 2 mg $ml^{-1}$ and then 500 µl was aliquoted to 10 x LoBind SafeLock 1.5 ml Eppendorf tubes. Duplicate samples were treated with 5 µl DMSO or compound **5a** (final concentration 0.01, 0.1, 1 and 10 µM), vortexed and incubated for 1 h at r.t. Samples were then processed according to the "*Cysteine MS-ABPP*" procedure. Freshly isolated splenocytes (red blood cells removed) from 6 adult C57BL/6 mice were washed with PBS (3x), then resuspended in PBS containing protease inhibitors and processed as above for PBMCs, except that protein concentration was normalized to 1.7 mg $ml^{-1}$.

*In situ treatments*: Freshly isolated PBMCs from a single blood donor were resuspended at 5 million cells/ml in serum free RPMI and 40 million cells were aliquoted to 10 x TC flasks. Cells were treated in duplicate with DMSO or compound **5a** (0.01, 0.1, 1, or 10 µM), incubated for 2 h at 37 °C, then harvested. Cell pellets were washed 2-3 x PBS, then frozen at -80 °C until processing. Frozen pellets were thawed on ice, resuspended in PBS containing protease inhibitors and lysed by sonication 3 x 10 pulses, 40% output. The proteome concentration of each sample was normalized to 2 mg $ml^{-1}$ and then 500 µl was taking forward for labeling with IA-DTB.

*Cysteine ABPP-MS:* Samples were treated with 5 µl of 10 mM iodoacetamide desthiobiotin (IA-DTB) (in DMSO), vortexed and incubated for 1 h at room temperature. Protein was then precipitated by the addition of 600 µl ice-cold MeOH, 200 µl CHCl3 and 100 µl $H_2O$, vortexed, and centrifuged (10 min, 10,000 g, 4 °C). Without disrupting the protein disk, both top and bottom layers of solvent were aspirated, and the protein disk was washed with 1 ml ice-cold MeOH, vortexed, and centrifuged (10 min, 16,000 g, 4 °C). Solvent was aspirated and the pellets were allowed to air dry (5-10 min, r.t.). Pellets were then re-suspended in 90 µl reducing buffer (9 M urea, 10 mM DTT, 50 mM TEAB, pH 8.5) and heated at 65 °C for 15 min, vortexing several times to solubilize proteins. Solutions were allowed to cool briefly, spun down, and then alkylated by treatment with 10 µl of 500 mM iodoacetamide and incubated for 30 min, 37 °C with shaking. Samples were then diluted with 300 µl 50 mM TEAB, pH 8.5 to reach final concentration of 2 M urea, and probe-sonicated (10 pulses) to ensure homogenous resuspension of all proteins. Samples were then digested overnight at 37 °C using 1 µg trypsin (resuspended at 0.25 µg $\mu l^{-1}$ in trypsin resuspension buffer supplemented with 25 mM $CaCl_2$) (V5111, Promega). Desthiobiotin labeled peptides were enriched from digested samples using 25 µl of packed streptavidin agarose resin per sample. Streptavidin was initially washed 3x with enrichment buffer (50 mM TEAB pH 8.5, 150 mM NaCl, 0.2% NP-40), and then 300 µl of resuspended agarose was added per sample. Samples were rotated at room temperature for 2-3 h, and then pelleted by centrifugation (2000 g, 2 min). Samples were transferred to BioSpin columns using 2 x 500 µl wash buffer (50 mM TEAB pH 8.5, 150 mM NaCl, 0.1% NP-40), then washed 2 x 1 ml wash buffer, 3 x 1 ml PBS, 3 x ml $H_2O$. Peptides were eluted by gravity into LoBind 1.5 ml Eppendorf tubes using 2 x 200 µL of 50% acetonitrile with 0.1% formic acid and evaporated to dryness in a SpeedVac vacuum concentrator. Dried samples were then resuspended in 100 μl of 70% 200 mM EPPS (pH 8), containing 30% acetonitrile, vortexed, and water bath sonicated (5 min). Samples were treated with 3 μl of a resuspended 10-plex TMT tag (20 μg $\mu l^{-1}$ in dry acetonitrile) and incubated at room temperature for 1 h with intermittent vortexing. Reactions were then quenched by the addition of 5 μl hydroxylamine (5% v/v in water, 15 min, r.t.), acidified with 5 μL formic acid, combined and dried using a SpeedVac.

Samples were resuspended in 500 μl Buffer A (5% MeCN, 0.1% formic acid), supplemented with 20 μL additional formic acid, and desalted using Sep-Pak C18 cartridge (WAT054955, Waters). In brief, cartridges were conditioned 3 x 1 ml 100% MeCN, equilibrated 3 x 1 ml Buffer A and then samples were flowed through 2 x under ambient pressure. Samples were washed 3x 1 ml Buffer A, then eluted into LoBind Eppendorf tubes using 1 ml Buffer B (80% MeCN, 0.1% formic acid), and evaporated to dryness in a SpeedVac.

Samples were resuspended in 500 μl Buffer A and fractionated by HPLC at a flow rate of 0.5 ml $min^{-1}$ using a capillary column (ZORBAX 300 Extend-C18, 3.5 μm) and the following gradient: 100% buffer A from 0-2 min, 0%–13% buffer B from 2-3 min, 13%–42% buffer B from 3-60 min, 42%–100% buffer B from 60-61 min, 100% buffer B from 61-65 min, 100%–0% buffer B from 65-66 min, 100% buffer A from 66-75 min, 0%–13% buffer B from 75-78 min, 13%–80% buffer B from 78-80 min, 80% buffer B from 80-85 min, 100% buffer A from 86-91 min, 0%–13% buffer B from 91-94 min, 13%–80% buffer B from 94-96 min, 80% buffer B from 96-101 min, and 80%–0% buffer B from 101-102 min (buffer A: 10 mM aqueous $NH_4HCO_3$; buffer B: acetonitrile). Peptides were eluted as 1 ml fractions into 96-well (deep-well) plates (Aglient) which contained 20 μl of 20% formic acid/well. The eluent was evaporated to dryness in the plate using SpeedVac, and then peptides were resuspended in Buffer B (80% MeCN, 0.1% formic acid) buffer (100 μl/well) and columns were combined to give 12 fractions. Samples were dried using SpeedVac, then re-suspended in 8 μl buffer A (5% acetonitrile, 0.1% formic acid) for mass spectrometry.

**TMT liquid chromatography-mass-spectrometry (LC-MS) analysis**

Samples were analyzed by liquid chromatography tandem mass-spectrometry using an Orbitrap Fusion Tribrid Mass Spectrometer (Thermo Scientific) coupled to an UltiMate 3000 Series Rapid Separation LC system and autosampler (Thermo Scientific Dionex). The peptides were eluted onto a capillary column (75 μm inner diameter fused silica, packed with C18 (Waters, Acquity BEH C18, 1.7 μm, 25 cm)) or an EASY-Spray HPLC column (Thermo ES902, ES903) using an Acclaim PepMap 100 (Thermo 164535) loading column, and separated at a flow rate of 0.25 μl $min^{-1}$. Data was acquired using an MS3-based TMT method. Briefly, the scan sequence began with an MS1 master scan (Orbitrap analysis, resolution 120,000, 400−1700 m/z, RF lens 60%, automatic gain control [AGC] target 2E5, maximum injection time 50 ms, centroid mode) with dynamic exclusion enabled (repeat count 1, duration 15 s). The top ten precursors were then selected for MS2/MS3 analysis. MS2 analysis consisted of quadrupole isolation (isolation window 0.7) of precursor ion followed by collision-induced dissociation (CID) in the ion trap (AGC 1.8E4, normalized collision energy 35%, maximum injection time 120 ms). Following the acquisition of each MS2 spectrum, synchronous precursor selection (SPS) enabled the selection of up to 10 MS2 fragment ions for MS3 analysis. MS3 precursors were fragmented by HCD and analyzed using the Orbitrap (collision energy 55%, AGC 1.5E5, maximum injection time 120 ms, resolution was 50,000). For MS3 analysis, we used charge state–dependent isolation windows. For charge state z = 2, the MS isolation window was set at 1.2; for z = 3-6, the MS isolation window was set at 0.7. RAW files were uploaded to Integrated Proteomics

Pipeline (IP2) and converted to MS2 and MS3 files. Files were searched using the ProLuCID algorithm (publicly available at http://fields.scripps.edu/downloads.php) using a reverse concatenated, non-redundant variant of the Human UniProt database (release 2016) or Mouse UniProt database (release 2017). Cysteine residues were searched with a static modification for carboxyamidomethylation (+57.02146 Da) and a dynamic modification for IA-DTB labeling (+398.25292 Da) (maximum 2 differential modifications per peptide). N-termini and lysine residues were also searched with a static modification corresponding to the respective TMT tag (+229.1629 Da). Peptides were required to be at least 6 amino acids long, to be fully tryptic (K or R cleavage sites), and to have a maximum of 2 mis-cleavage sites. ProLuCID data was filtered through DTASelect (version 2.0) to achieve a peptide false-positive rate below 1%. The MS3-based peptide quantification was performed with reporter ion mass tolerance set to 20 ppm with Integrated Proteomics Pipeline (IP2).

**Data processing**
Cysteine engagement (DMSO vs inhibitor) was calculated for each peptide-spectra match by dividing each TMT reporter ion intensity by the average intensity for the channels corresponding to DMSO treatment. Peptide-spectra matches were then grouped based on protein ID and residue number (e.g., JAK1_C817). Data were filtered to exclude peptides with summed reporter ion intensities for the DMSO channels < 10,000, a coefficient of variation for DMSO channels > 0.5, non-tryptic or reverse peptide sequences. Cysteine engagement is reported as either a % of peptide signal intensity (S.I.) in DMSO-treated channels, or a competition ratio (S.I DMSO/S.I. compound) and are the mean value of the 2 replicate channels in each TMT 10-plex sample. Competition ratios ≥4 (corresponding to a ≥75% loss in peptide S.I. compared to DMSO) were considered significantly engaged by inhibitors.

***In vitro* and *in vivo* target engagement**
*In vitro* $TE_{50}$ values for JAK1_C817 and TYK2_C838 were obtained by treating 500 µg of cell lysate generated from primary human PBMCs (AllCells), Jurkat T-cells (ATCC –TIB-152), or MDA-MB-468 breast cancer cells (ATCC HTB-132), with DMSO or compound for 1 hour at room temperature, followed by the addition of 200 µM iodoacetamide desthiobiotin (IA-DTB) (in DMSO) for 1 hr at room temperature. Samples were then precipitated by the addition of 8x ice cold acetone and incubated at -80°C for two hours. Protein was then pelleted by centrifugation (4,200 RPM, 45 min, 4°C). The pelleted material was resuspended in 50 mM ammonium bicarbonate buffer containing 9M Urea, and proteins were reduced and alkylated by the addition of DTT (10 mM) and iodoacetamide (30 mM). Following reduction and alkylation, samples were exchanged into 2M urea (Zeba spin desalting plates, Thermo Fisher) and digested with trypsin. IA-DTB-labeled peptides were isolated with by enrichment with streptavidin agarose resin. Target engagement was determined from isolated peptides by PRM or tTMT as described below.

*In vivo* engagement of JAK1 C817 by compound **5** was determined from whole spleen lysate of mice (n = 4 per group) that had been treated subcutaneously with 2 doses of compound **5** (25 mg $kg^{-1}$) or DMSO vehicle (prepared as 5% DMSO/20% hydroxy-propyl-β-cyclodextrin). Mice were sacrificed 4 hours after the second dose. Harvested spleens were homogenized in 700 µL of PBS by bead beating (1.4 mM ceramic beads, Omni Bead Ruptor Elite) for 30 seconds. Samples were further homogenized by sonication in a water bath and clarified by centrifugation (1,000 x g for 10 mins.). Protein concentration was determined and 1.5 mgs of lysate was treated with 200 μM IA-DTB (in DMSO) for 1 hour at room temperature, and then samples were processed and analyzed by PRM as described below.

**Parallel reaction monitoring**
Peptides were eluted from an EASY-Spray C18 loading column (5 μm particle size, 100 μm × 2 cm; Fisher Scientific, DX164564) and resolved on a custom analytical column (2 μM particle size, 75 μm x 15 cm) using a Dionex Ultimate 3000 nano-LC (Thermo Fisher Scientific). Peptides were separated over an 11-min gradient of 6 - 33% acetonitrile (0.1% v/v formic acid) and analyzed by parallel reaction monitoring (PRM) on a Q-Exactive instrument (Thermo Fisher Scientific). Selected precursor ions targeting JAK1_C817 (amino acids 810 - 818, +3 charge state) and TYK2_C838 (amino acids 833 - 860, +3 charge state) were isolated and fragmented by high-energy collision-induced dissociation and fragments were detected in the Orbitrap at 17,500 resolution. The resulting PRM data were analyzed using Skyline (v.21.1.0.278) and quantification was performed by summing the peak areas corresponding to six fragment ions from each peptide. The peptides and fragment ions were pre-selected from in-house reference spectral libraries acquired in data-dependent acquisition mode to identify authentic spectra for each peptide.

**Targeted TMT**
Targeted TMT measurements were collected using an Orbitrap LumosTribrid Mass Spectrometer (Thermo Scientific) coupled to an UltiMate 3000 Series Rapid Separation LC system and autosampler (Thermo Scientific Dionex). Peptides were eluted onto a custom C18 capillary analytical column (75 μm inner diameter fused silica, packed with Acclaim PepMap C18 resin (Thermo Scientific)) using an Acclaim PepMap 100 (Thermo 164535) loading column, and separated at a flow rate of 1.0 μl $min^{-1}$. Data were acquired using a specific MS3-based TMT method targeting peptides containing JAK1_C817 (amino acids 810 - 818, +3 charge state) and TYK2_C838 (amino acids 830 – 860, +5 charge state), where MS2 peptide fragmentation is triggered upon detection of the peptide precursor ion. Subsequent MS3 analysis was then performed using pre-selected peptide fragment ions that were isolated for fragmentation using synchronous precursor selection. RAW files were converted to MZXML format and searched with the SEQUEST algorithm using the MassPike software package. TMT quantitation was performed with a filter requiring at least 10 summed signal-to-noise for control channels.
